# Genetically Encoded Red Fluorescent Indicators for Imaging Intracellular and Extracellular Potassium Ions

**DOI:** 10.1101/2024.12.20.629597

**Authors:** Lina Yang, Vishaka Pathiranage, Shihao Zhou, Xiaoting Sun, Hanbin Zhang, Cuixin Lai, Chenglei Gu, Fedor V. Subach, Alice R. Walker, Kiryl D. Piatkevich

**Affiliations:** Fudan University, Shanghai, China; School of Life Science, Westlake University, Hangzhou, Zhejiang, China; Westlake Laboratory of Life Sciences and Biomedicine, Hangzhou, Zhejiang, China; Institute of Basic Medical Sciences, Westlake Institute for Advance Study, Hangzhou, Zhejiang, China; Department of Chemistry, Wayne State University, Detroit, MI, 48202; Complex of NBICS Technologies, National Research Center “Kurchatov Institute”, Moscow, 123182, Russia

**Keywords:** genetically encoded red potassium indicators, directed molecular evolution, intracellular potassium imaging, extracellular potassium imaging

## Abstract

Potassium ion (K^+^) dynamics are vital for various biological processes. However, the limited availability of detection tools for tracking intracellular and extracellular K^+^ has impeded a comprehensive understanding of the physiological roles of K^+^ in intact biological systems. In this study, we developed two novel red genetically encoded potassium indicators (RGEPOs), RGEPO1 and RGEPO2, through a combination of directed evolution in *E. coli* and subsequent optimization in mammalian cells. RGEPO1, targeted to the extracellular membrane, and RGEPO2, localized in the cytoplasm, exhibited positive K^+^-specific fluorescence response with affinities of 3.55 mM and 14.81 mM in HEK293FT cells, respectively. We employed RGEPOs for real-time monitoring of subsecond K^+^ dynamics in cultured neurons, astrocytes, acute brain slices, and the awake mouse in both intracellular and extracellular environments. Using RGEPOs, we were able, for the first time, to visualize intracellular and extracellular potassium transients during seizures in the brains of awake mice. Furthermore, molecular dynamics simulations provided new insights into the potassium-binding mechanisms of RGEPO1 and RGEPO2, revealing distinct K^+^-binding pockets and structural features. Thus, RGEPOs represent a significant advancement in potassium imaging, providing enhanced tools for real-time visualization of K^+^ dynamics in various cell types and cellular environments.

## Introduction

Potassium ion (K^+^) homeostasis and dynamics play critical roles in a multitude of biological activities in all domains of life^1,2^. For example, in neurons, K^+^ is vital for generating action potentials and maintaining the resting membrane potential, which is critical for proper signal transmission^3,4^. Astrocytes, on the other hand, regulate extracellular K^+^ levels through a process known as potassium buffering, which is essential for preventing neuronal hyperexcitability and maintaining overall ion homeostasis^5^. Under resting conditions in the mammalian cells, K^+^ has an intracellular concentration of 142 to 175 mM, while extracellular concentration is in the range of 3.5 to 5 mM, maintaining the resting membrane potential at -70 to -80 mV^6,7^. Therefore, sensitive ion probes with a high dynamic range at physiological concentrations are essential for accurately analyzing K^+^ homeostasis and its functional transients.

Traditionally, K^+^ levels have been measured using ion-sensitive electrodes^8,9^ or fluorescent dyes^10–12^. While ion-sensitive electrodes provide direct and precise measurements of absolute K^+^ concentrations^9^, they are highly invasive and typically offer limited spatial resolution and throughput of measurements, making them unsuitable for studying fine-scale cellular processes in intact cells and tissues. Fluorescent dyes, such as potassium-sensitive fluorophores, offer improved spatial resolution compared to electrodes and allow for non-invasive imaging of K^+^ levels in live cells^10,12^. However, they suffer from drawbacks such as poor ion selectivity, labeling specificity, limited dynamic range, cytotoxicity, and lack of targetability to specific cell types or subcellular compartments.

Genetically encoded potassium indicators (GEPOs) provide a promising alternative for K^+^ measurement, allowing for real-time, minimally invasive monitoring with high spatial and temporal resolution^13–16^. Recent achievements in developing GEPOs stemmed from using a small water-soluble potassium-binding protein from *E. coli*, known as Ec-Kbp (formerly YgaU)^17^. Among these indicators, GEPII^16^ and KIRIN1s^15^ are based on the Förster Resonance Energy Transfer (FRET) (Supplementary Table 1). The FRET-based GEPOs provide ratiometric output, which enhances the accuracy of concentration measurements. However, they require two specific emission color bands, limiting their application in multiplexed imaging experiments. Additionally, FRET-based indicators often suffer from weak signals and limited dynamic range. In contrast, single fluorescent protein (FP)-based indicators, although more challenging to engineer than FRET-based biosensors and typically intensiometric, require only single-color bands for excitation and emission. This makes them well-suited for multiparameter imaging^18^, and they are generally characterized by higher dynamic range and stronger signals. The green GEPOs, GINKO1 (ref.^15^), GINKO2 (ref.^13^), and KRaIONs^14^, based on the single GFP-like FPs, have been successfully applied to monitor intracellular K^+^ dynamics in mammalian cells and plants (Supplementary Table 1). Compared to green fluorescence proteins, red fluorescence proteins provide the advantage of deeper tissue penetration, low phototoxicity, and compatibility with a large variety of existing green indicators. Thus, the development of red fluorescent GEPOs (RGEPOs) is essential for advancing K^+^ research.

By integrating insights garnered from genomic mining and informed by structural guidance in potassium-binding protein architecture, we successfully developed two red fluorescent potassium indicators, designated as RGEPO1 and RGEPO2. These indicators were engineered by inserting a potassium-binding protein cloned from a hydrothermal vent bacterium into the fluorescent protein mApple at β-strand 7 next to the chromophore. RGEPO1 and RGEPO2 exhibited a ΔF/F in the range of 400-1000% in solution and 100-300% in HEK293FT cells in response to 150 mM and 200 mM K^+^, respectively. In HEK cells, potassium affinities were determined to be 3.55 mM for RGEPO1 and 14.81 mM for RGEPO2. Additionally, we discovered the molecular mechanisms underlying the different responses of RGEPO1 and RGEPO2 to potassium ions using molecular dynamic simulation. Furthermore, RGEPOs were applied to monitor intracellular and extracellular K^+^ dynamics in the culture neurons, astrocytes, acute brain slices, and mouse seizures *in vivo*. These findings substantiate the practical utility of indicators RGEPO1 and RGEPO2 as essential tools for the real-time visualization and comprehensive analysis of intracellular and extracellular potassium ion dynamics.

## Results

### Development and validation of red fluorescent potassium indicator

To engineer a single red fluorescent protein-based K^+^ indicator, we initiated our efforts by identifying a large diversity of putative naturally occurring potassium binding proteins, aiming to develop a sensor with a larger dynamic range that can be effectively applied across various biological contexts. For genome mining, we used protein BLAST on the ’env_nr’ dataset. This dataset includes proteins from whole genome shotgun sequencing (WGS) metagenomic projects, which was searched using the Hv-Kbp protein (see the original publication reporting its function^14^) as a query sequence. The search identified five proteins with putative functions that shared 40%-70% amino acid identity with the Hv-Kbp protein and had conserved K^+^ binding sites based on the primary sequence alignment with the Ec-Kbp protein, which has confirmed potassium binding pocket with 6 amino acids coordinating the cation (Supplementary Fig. 1). To validate their potassium binding properties, we swapped Ec-Kbp in the KRaION1 indicator^14^ with each of these homologs and expressed them in *E. coli* to assess sensitivity to K^+^ (Fig. 1a). All generated constructs exhibited positive fluorescence responses to K^+^, confirming their function as K^+^-binding proteins (Supplementary Fig. 2). Among them, Hv-Kbp-mNG and MNT-mNG displayed the largest ΔF/F values, reaching 59%, and 62%, respectively, while also exhibiting the highest baseline fluorescence (Fig. 1b) making them primary candidates, along with Ec-Kbp, for developing red potassium indicators.

**Figure 1.**
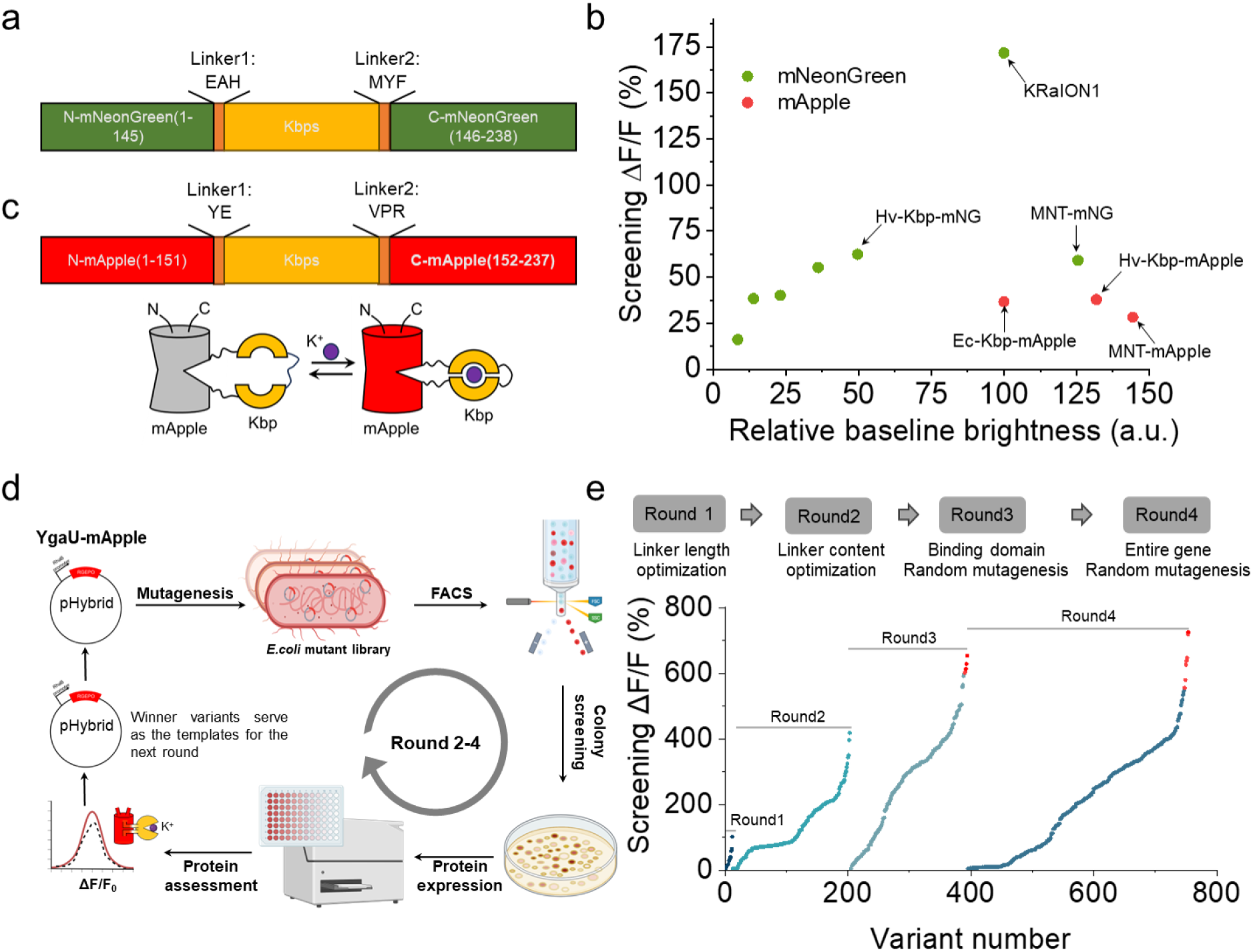
Development and optimization of red potassium biosensors in *E. coli*. a. Schematic representation of framework of Kbp’s homologues inserted into mNeoGreen. b. Screening ΔF/F vs. screening brightness rank plot representing the homologues of Kbp tested in *E. coli* based on mNeoGreen and mApple, respectively. c. Schematic representation of RGEPO and its putative mechanism of response to K^+^. d. Schematic depiction of the random mutagenesis workflow. Random mutagenesis was applied to the binding domain or the entire gene of the template, yielding a mutant library. Generated mutants were selected using FACS based on brightness. Protein extraction was carried out, and ΔF/F was determined upon the addition of 200 mM K^+^. The variants exhibiting the highest ΔF/F were employed as the template for the next round. FACS, fluorescence-activating cell sorting (created in BioRender.com). e. In vitro-stages in the identification of potential RGEPO candidates. The optimization process comprised four distinct steps, which involved genome mining for insertion site variation, the optimization of linker length, linker content screening, and random mutagenesis of both the binding domain and the entire gene. ΔF/F rank plot representing all variants assessed during the directed evolution. For each round, tested variants are ranked from lowest to highest ΔF/F value from left to right. The red highlight dots are the candidates for validation in cultured mammalian cells.

To construct a red fluorescence potassium sensor, we inserted these two homologs as well as Ec-Kbp into the mApple FP between amino acids S151 and V152 derived from the previously developed and characterized FRCaMP calcium sensor^19–21^ (following the previous design of red Ca^2+^ sensor FRCaMP^19^ but with inverted topology as described in ref.^20^; Fig. 1c) and expressed the constructs in *E. coli* for further characterization. All three variants were functional, although they exhibited limited ΔF/F below 30% with comparable baseline brightness (Fig. 1b). Considering the brightness and ΔF/F, we selected the Hv-Kbp-based chimera for further optimization through directed molecular evolution in *E. coli*, which allows for rapid and efficient high-throughput screening of libraries to enhance the prototype^13,22^ (Fig. 1d). Previous studies on high-performance biosensors revealed the critical role of linkers between sensing and reporting moieties in biosensor functionality and perfromance^23,24^. Therefore, we started by optimizing the length and amino acid composition of the linkers near the insertion site, ultimately identifying a prototype sensor with linker sequences Linker1: P and Linker2: SAP (Supplementary Fig. 3a). This prototype exhibited a robust fluorescence increase with ΔF/F of 418% in response to 230 mM K^+^ (Fig. 1e, Round 2). Further optimization using random mutagenesis, targeting both the binding domain and the entire sensor sequence, produced variants with maximum ΔF/F values exceeding 600% (Fig. 1e and Supplementary Fig. 3b).

Sensor properties in solution are not always readily translated to mammalian cells^22^. Since our goal was to develop a sensor suitable for imaging mammalian cells, we selected the top-performing variants identified through *E. coli* screening for side-by-side comparison in HEK293FT cells. To evaluate these candidates in a cellular environment, we expressed and assessed them in HEK293FT cells, titrating with 200 mM K^+^ in the presence of 15 μM valinomycin and 5 μM CCCP, which facilitated K^+^ influx and maintained a stable pH (Fig. 2a). This K^+^ concentration, slightly above typical levels in mammalian cells^25^, was chosen to enhance the interaction between potassium ions and the indicator, ensuring a robust and detectable response for effective screening. Among these, RGEPO0.5 produced a peak change in fluorescence of 53.37% (Fig. 2b), which was 13-fold lower than that in the solution. We attribute this discrepancy to three primary factors: first, the intracellular environment may alter protein conformation, affecting sensor responsiveness to K^+^. Second, despite the addition of K^+^-specific ionophores like valinomycin and CCCP, ion permeation remains a challenge, resulting in lower effective ion concentrations interacting with the sensor in cells compared to direct in vitro assays. Third, in solution titrations start from 0 mM, whereas intracellular conditions inherently contain potassium ions, affecting baseline fluorescence responses. These challenges necessitate further optimization to enhance ΔF/F in mammalian cells.

**Figure 2.**
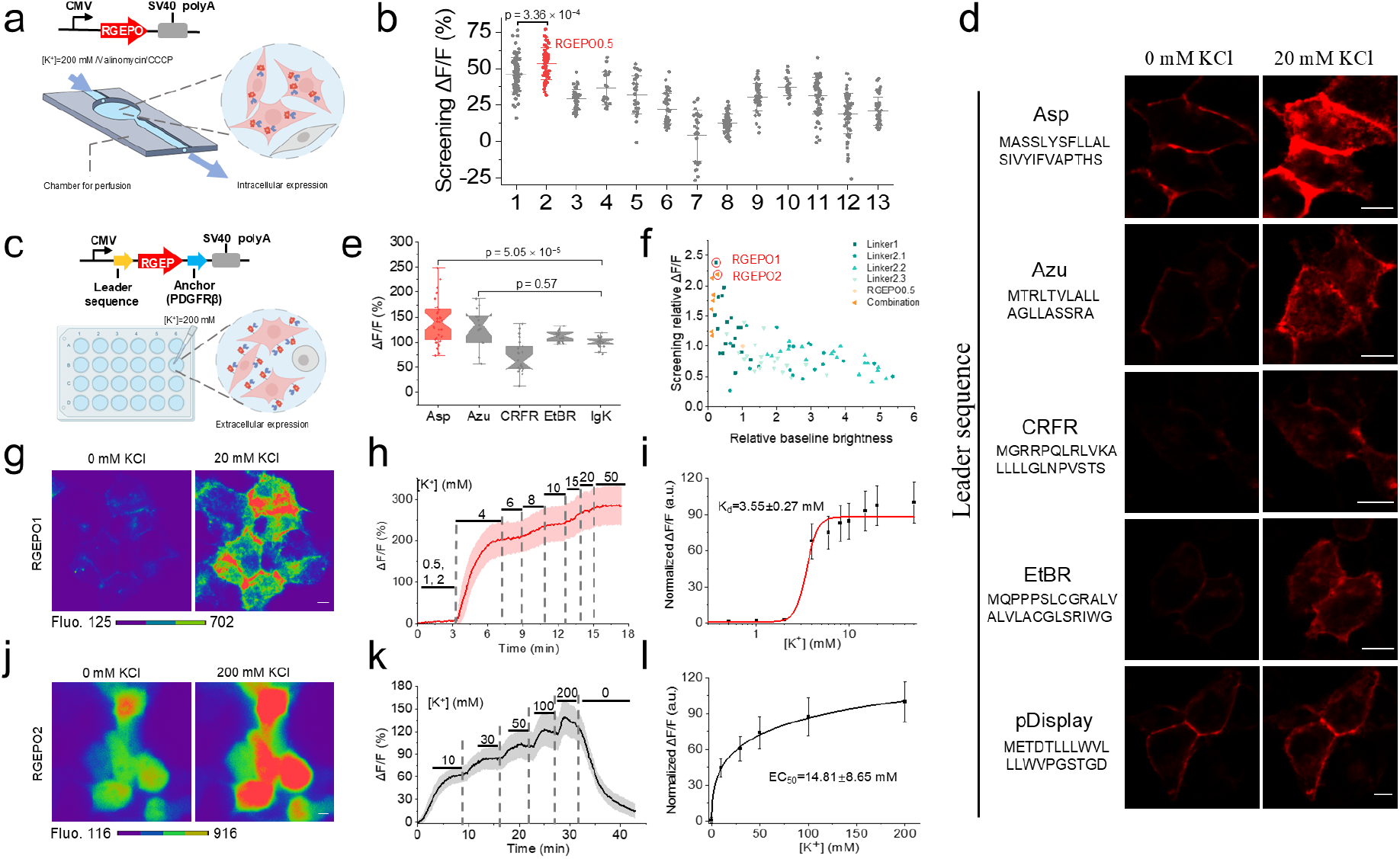
Optimization in HEK cells. a. The linear diagram of the expression cassette for RGEPO expression in the mammalian cells(top) and schematic representation of the automated perfusion system employed in buffer exchange experiments. b. Validation of candidates selected in E. coli using HEK cells via perfusing K^+^ free and 200 mM K^+^ containing buffers(n= 80, 50, 46, 22, 29, 50, 29, 54, 39, 26, 65, 66, 43 cells from 1 culture for 1st to 13th mutant). One-way ANOVA with Benferroni’s test was performed. p = 3.35775 × 10^−4^ between mutant1 and mutant2. This screening led to RGEPO0.5. c. A linear map of the expression cassette for RGEPO expression at the extracellular surface of the plasma membrane (top) and a schematic depiction of the detection system employed to assess the response of RGEPO. d. Representative single-plane confocal fluorescence images of leader-sequence variants (n= 3 cells from 2 FOVs over 1 culture for each). Scale bars, 10 µm. e. ΔF/F of a range of leader sequence variants in response to 200 mM K^+^ (n= 31, 13, 22, 22, 24 cells from 1 culture for Asp, Azu, CRFR, EtbR and Igκ, respectively). Box plots with notches are used: narrow part of notch, median; top and bottom of the notch, 95% confidence interval for the median; top and bottom horizontal lines, 25% and 75% percentiles for the data; whiskers extend 1.5 × the interquartile range from the 25th and 75th percentiles; horizontal line, mean; outliers, dots. One-way ANOVA with Benferroni’s test was performed. P = 5.0538 × 10^−5^ between Asp and Igκ. P = 0.05682 between Azu and Igκ. f. Saturation mutagenesis screening and recombination of linker content on the extracellular surface of cultured mammalian cells with the Asp leader sequence, continuing from the template in b. This screening led to RGEPO1 and RGEPO2 indicated with a red circle. g. Representative images of expression and fluorescence change of RGEPO1 in response to 20 mM K^+^. Scale bar, 10 µm. h. Fluorescence intensity change (ΔF/F) time course of RGEPO1 with stimulation by a series of K^+^ buffers on HEK293T cells (n= 18 cells from 1 culture), data are expressed as mean ± SD. i. Plot of normalized ΔF/F against different K^+^ concentrations fitted using nonlinear fitting (Hill) for the data shown in panel h. j. Representative images of expression and fluorescence change of RGEPO2 in response to 200 mM K^+^. Scale bar, 10 µm. k. Fluorescence intensity change (ΔF/F) time course of RGEPO1 with stimulation by a series of K+ buffer on HEK cells using valinomycin and CCCP (n= 32 cells from 1 culture), data are expressed as mean ± SD. l. Plot of normalized ΔF/F against different K^+^ concentrations fitted using nonlinear fitting (Hill) for the data shown in panel k.

To enhance the screening efficiency in HEK cells, we decided to express the sensor on the extracellular surface of the plasma membrane, which facilitates rapid buffer exchange to modulate sensor fluorescence and thus increase the throughput of the screening. Using RGEPO0.5 as a template for preliminary extracellular probe construction, we targeted it to the outer surface of HEK cells by fusing it to five N-terminal leader sequences from naturally occurring proteins (see Methods for details) and the platelet-derived growth factor receptor (PDGFRβ) transmembrane domain, respectively (Fig. 2c). While all generated constructs exhibited membrane localization and functional responses to K^+^, the Asp leader peptide variant demonstrated a superior response, with a 1.4-fold higher ΔF/F compared to the widely used leader sequence of immunoglobulin k-chain (Igk) from pDisplay system^26^ (Fig. 2d, e and Supplement Fig. 4). Additionally, compared to pDisplay-EGFP as a cell surface marker, the combination of Asp leader and PDGFRβ anchor also effectively targeted RGEPO0.5 to the cell surface (Supplement Fig. 5). Utilizing the Asp-PDGFRβ cell surface expression system, we performed saturation mutagenesis on the linker content. Stimulated with 200 mM K^+^, the P299C mutant exhibited the highest response among the single and recombined linker mutations and was therefore designated as RGEPO1 (Fig. 2f). Subsequently, we validated the top three mutants with the highest extracellular response by expressing them intracellularly and measuring fluorescence response through intracellular perfusion with 200 mM K^+^ in the presence of 15 μM valinomycin and 5 μM CCCP. Among them, P152L/A298N showed the highest response to K^+^ intracellularly and was thus designated as RGEPO2 (Fig. 2f and Supplementary Fig. 6). As a result, RGEPO1 and RGEPO2 had G264V mutations in the binding domain but differ only by linker content between each other (see Supplementary Fig. 7 for amino acid alignment of the prototype, RGEPO0.5, and RGEPO1,2).

To characterize the Kd and the dynamic range of RGEPOs and confirm the reversibility of the fluorescence response, we performed serial titrations with K^+^ in HEK293FT cells. The Asp- RGEPO1 sensor exhibited clear localization to the cell membrane. The addition of K^+^ triggered a robust fluorescence increase of cell surface-localized RGPEO1 (Fig. 2g), which returned to baseline after washing out K^+^ using a perfusion system (Supplement Fig. 8a). The fluorescence increase of RGEPO1 was dependent on K^+^ concentration locally applied with a micropipette, achieving a ΔF/F of 286% at 50 mM (Fig. 2g,h; for visualization of fluorescence changes see Supplementary Movie 1). The Kd at the cell surface was determined to be 3.55±0.27 mM enabling detection of [K^+^] in the range of 4-15 mM, with a Hill coefficient of 4.0±1.0 (Fig. 2i). In parallel, RGEPO2 was observed to localize evenly within the cytoplasm and nucleus without any visible aggregates (Supplementary Fig. 8b). We sequentially introduced solutions containing varying K^+^ concentrations from 0 to 200 mM along with valinomycin and CCCP. The fluorescence intensity changes of RGEPO2 were also dependent on the K^+^ concentrations, resulting in a ΔF/F of 139% at 200 mM with an affinity of ∼ 14.81±8.65 mM and a Hill coefficient of 0.49±0.1 (Fig. 2j,k,l; for visualization of fluorescence changes see Supplementary Movie 2). Fluorescence of RGEPO2 returned to the baseline after washing out K^+^, indicating full reversibility of the sensor (Fig. 2k, Supplementary Fig. 8c). As a result, we developed a pair of red GEPOs for detecting intracellular and extracellular K^+^ dynamics in the wide range of potassium concentrations from 4 to 200 mM. We utilized a hybrid workflow for evolution, combining high throughput large library screening in bacteria followed by optimization in HEK cells. We used a new domain, Hv-Kbp, which was not used for the sensor development before, although it has been reported^14^.

### Characterization of RGEPOs in solution

To evaluate the applicability of the sensor in complex mammalian systems, it is essential to assess its fundamental biophysical properties in highly controlled environments, such as solutions. To this end, we performed a comprehensive characterization of the spectral and biochemical properties of RGEPO1 (without Asp leader or PDGFβ anchor) and RGEPO2 in the physiological range of [K^+^] in mammalian cells, i.e., from 0 mM to 150 mM [K^+^] at pH 7.4 (ref.^25^). In a buffer without potassium, RGEPO1 and RGEPO2 exhibited two absorbance peaks at 448/446 and 576/578 nm, indicative of the protonated and the deprotonated chromophore forms, respectively (Fig. 3a). Upon adding 150 mM K^+^ to the purified proteins, we observed an increase in the absorption at 576/578 nm and a simultaneous absorption decrease at 448/446 nm (Fig. 3a), suggesting a K^+^ dependent chromophore deprotonation process. Correspondingly, excitation peaks were observed at 575 nm for RGEPO1 and 574 nm for RGEPO2, leading to emission maxima at 591 nm and 593 nm, respectively (Fig. 3b, Supplement Fig. 9). In the K^+^ binding state, the excitation maximum shifted slightly blue to 563 nm (Supplementary Fig. 9), most likely due to the influence of the potassium ion on the chromophore environment. Upon the addition of 150 mM K^+^, RGEPO1 emission showed a 4.7-fold intensiometric increase at its peak of 591 nm, and RGEPO2 exhibited a 3.1-fold at its peak of 593 nm (Fig. 3b). The enhanced fluorescence intensity was attributed to increases in both extinction coefficients and quantum yields (Table 1). For RGEPO1, the extinction coefficient increased from 10,700 ± 900 to 41,600 ± 500 M^-1^·cm^-1^, and for RGEPO2, it increased from 9,700 ± 1,200 to 26,850 ± 2,960 M^-1^·cm^-1^. Concurrently, the quantum yields (QYs) under these conditions increased by ∼1.7- fold for RGEPO1 and ∼1.4-fold for RGEPO2 (Table 1). Together, these findings highlight the significant potential of RGEPO1 and RGEPO2 as effective indicators for monitoring potassium ion concentrations, attributed to their substantial changes in spectral properties upon K^+^ binding. We also characterized the pH sensitivity of RGEPOs, which observed significant fluorescence changes in response to 150 mM K^+^ within a pH range of 6.5 to 9.0. RGEPO1 achieved a maximum dynamic range at pH 8.0 and RGEPO2 at pH 7.4, with respective pKa values of 8.55 and 8.55 in the K^+^-free state and pKa values of 7.36 and 7.82 in the K^+^-bound state for RGEPO1 and RGEPO2, respectively (Fig. 3c,d). Thus, the RGEPOs’ fluorescence was sensitive to physiologically relevant pH change. To investigate the metal ion selectivity, we titrated RGEPOs with a variety of ions: Li^+^, Na^+^, Rb^+^, Cs^+^, NH4^+^, Mg^2+^, Ca^2+^, and Zn^2+^, along with K^+^. Among all tested ions, only Rb^+^, Ca^2+^, and Zn^2+^ induced considerable fluorescence increases of RGEPO1 / RGEPO2 by 109.7%/43.6%, 45.5%/72.7%, and 127.1%/100.0%, respectively (Fig. 3e). At the highest concentrations of the tested ions typically found in mammalian cells^27–34^, ΔF/F for K^+^ were the highest at 844% for RGEPO1 and 584% for RGEPO2 significantly surpassing responses to other ions. Median effective concentrations (EC50) were determined as 14 mM for RGEPO1 and 59 mM for RGEPO2 under isotonic conditions with varying K^+^ concentrations up to 245 mM (Fig. 3f). The corresponding Hill coefficients were 0.88 ± 0.08 for RGEPO1 and 1.1 ± 0.07 for RGEPO2. According to these results, the estimated dynamic range of K^+^ detection in solution was from ∼0.1 mM to ∼225 mM. Overall, these results demonstrate the strong potential of RGEPOs as selective and sensitive indicators for potassium ions in varying physiological environments.

**Figure 3.**
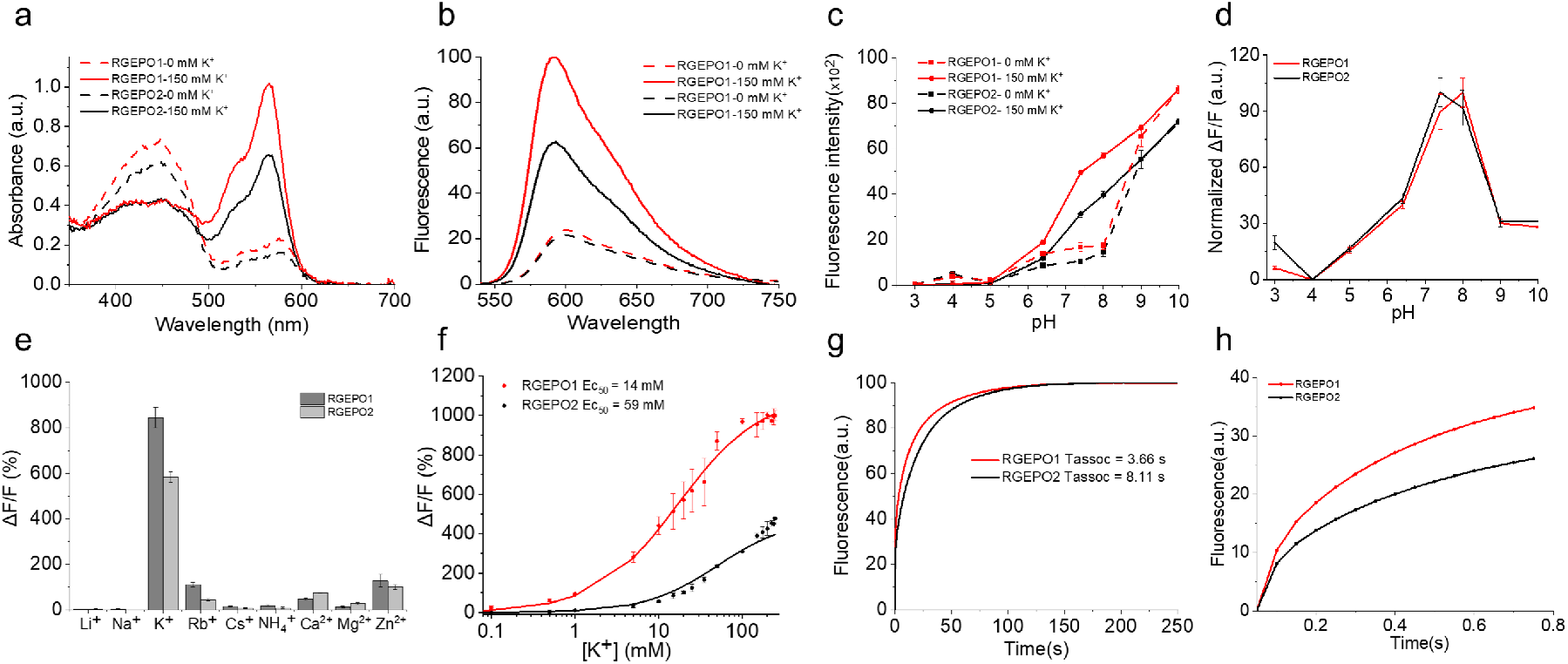
In vitro characterization of RGEPOs. **a.** Absorbance spectra of RGEPOs at 0 and 150 mM potassium at pH = 7.4. **b.** Steady-state fluorescence spectra of RGEPOs at 0 and 150 mM potassium at pH = 7.4. **c.** Fluorescence intensity of RGEPOs at 0 and 150 mM potassium as a function of pH (n = 3 technical replicates; mean ± SD). **d.** Normalized fluorescence changes upon the addition of 150 mM potassium as a function of pH. **e.** Ion selectivity of RGEPOs (n=3 technical replicates; mean ± SD). The concentrations of cations used were above their physiological concentrations. **f.** K^+^ titration of RGEPOs at isotonic conditions (n=3 technical replicates for RGEPO1 and RGEPO2, respectively). **g.** Association kinetics of RGEPOs at 150 mM K^+^ measured by stopped-flow fluorimetry (n=3 independent technical replicates each). h. the same association kinetics curve as g shown in the range of the 0−0.75 s time frame.

**Table 1.**
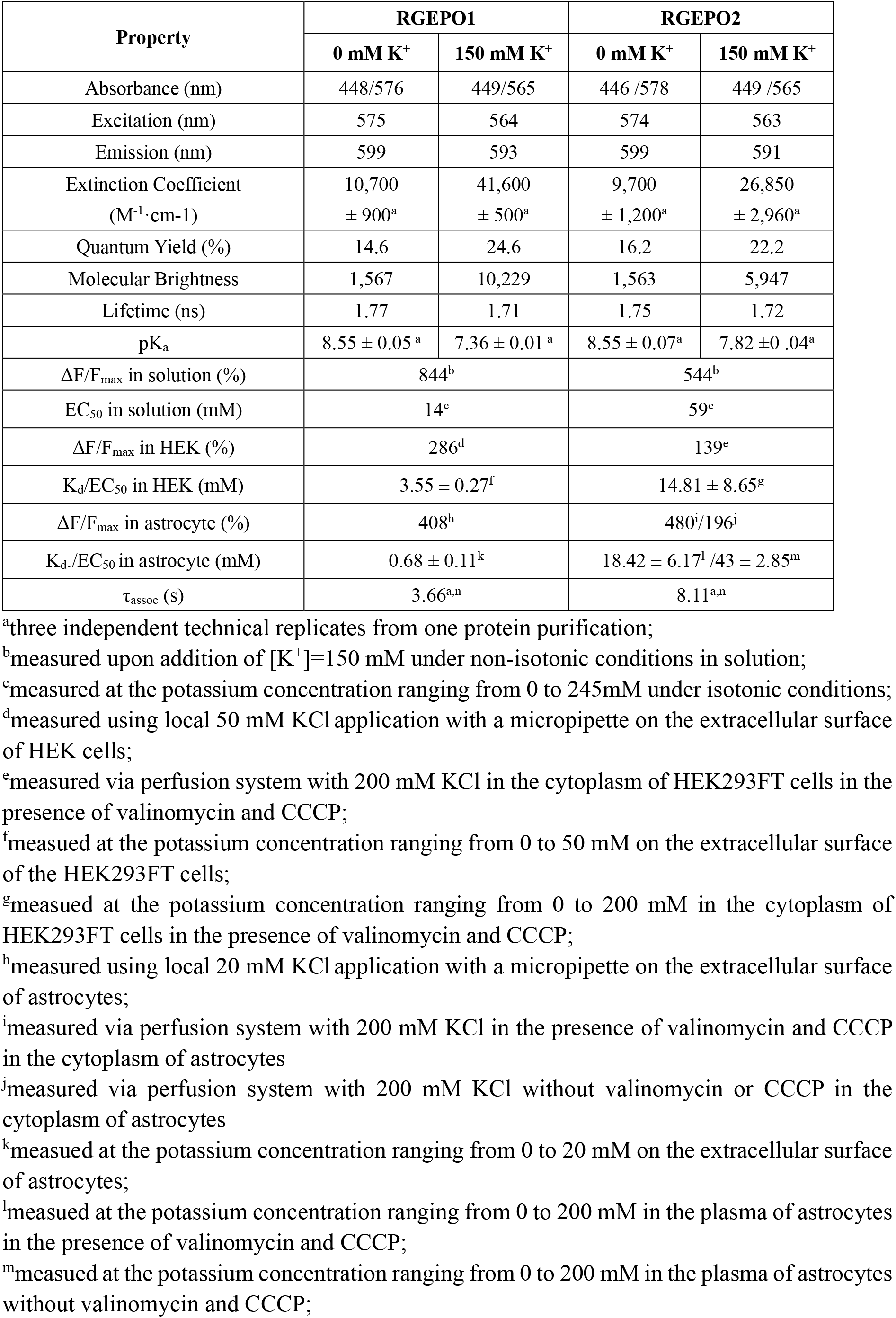
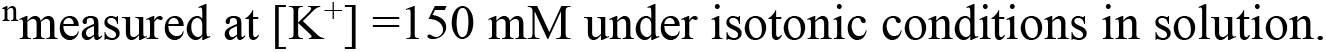
Spectral and biochemical properties of RGEPOs in solution.

| Property | RGEPO1 |  | RGEPO2 |  |
| --- | --- | --- | --- | --- |
|  | 0 mM K<sup>+</sup> | 150 mM K<sup>+</sup> | 0 mM K<sup>+</sup> | 150 mM K<sup>+</sup> |
| Absorbance (nm) | 448/576 | 449/565 | 446 /578 | 449 /565 |
| Excitation (nm) | 575 | 564 | 574 | 563 |
| Emission (nm) | 599 | 593 | 599 | 591 |
| Extinction Coefficient<br>(M<sup>-1</sup>·cm-1) | 10,700<br>± 900<sup>a</sup> | 41,600<br>± 500<sup>a</sup> | 9,700<br>± 1,200<sup>a</sup> | 26,850<br>± 2,960<sup>a</sup> |
| Quantum Yield (%) | 14.6 | 24.6 | 16.2 | 22.2 |
| Molecular Brightness | 1,567 | 10,229 | 1,563 | 5,947 |
| Lifetime (ns) | 1.77 | 1.71 | 1.75 | 1.72 |
| pK<sub>a</sub> | 8.55 ± 0.05 <sup>a</sup> | 7.36 ± 0.01 <sup>a</sup> | 8.55 ± 0.07<sup>a</sup> | 7.82 ± 0.04<sup>a</sup> |
| ΔF/F<sub>max</sub> in solution (%) | 844<sup>b</sup> |  | 544<sup>b</sup> |  |
| EC<sub>50</sub> in solution (mM) | 14<sup>c</sup> |  | 59<sup>c</sup> |  |
| ΔF/F<sub>max</sub> in HEK (%) | 286<sup>d</sup> |  | 139<sup>e</sup> |  |
| K<sub>d</sub>/EC<sub>50</sub> in HEK (mM) | 3.55 ± 0.27<sup>f</sup> |  | 14.81 ± 8.65<sup>g</sup> |  |
| ΔF/F<sub>max</sub> in astrocyte (%) | 408<sup>h</sup> |  | 480<sup>i</sup>/196<sup>j</sup> |  |
| K<sub>d</sub>/EC<sub>50</sub> in astrocyte (mM) | 0.68 ± 0.11<sup>k</sup> |  | 18.42 ± 6.17<sup>l</sup> /43 ± 2.85<sup>m</sup> |  |
| τ<sub>assoc</sub> (s) | 3.66<sup>a,n</sup> |  | 8.11<sup>a,n</sup> |  |

Potassium binding kinetics were further explored at 150 mM K^+^ using stopped-flow fluorimetry, which revealed an exponential association curve for both sensors. The components of this curve led to a 50% fluorescence change within approximately 3.66 seconds for RGEPO1 and 8.11 seconds for RGEPO2 (Fig. 3g,h). Additionally, we measured fluorescence lifetimes. The average fluorescence lifetimes for RGEPO1 in the K^+^-free and K^+^-bound states were 1.77 and 1.71 ns, respectively. For RGEPO2, the lifetimes were 1.75 and 1.72 ns in the K^+^-free and K^+^-bound states, respectively (Table 1). Altogether, we characterized the fluorescence spectra, brightness, affinity, pH sensitivity, metal ion specificity, and kinetics of RGEPOs, demonstrating that RGEPO1 and RGEPO2 can serve as sensitive and selective indicators for monitoring K^+^ dynamics across various physiological environments.

### Molecular mechanism of potassium sensitivity

To explore the molecular mechanism underlying potassium binding, we performed classical molecular dynamics (MD) simulations for both the RGEPO1 and RGEPO2 systems in solution at low (20 mM) and high (300 mM) potassium concentrations. Low concentration was selected to match the affinity of the sensors in the solution, and high concentration corresponds to the upper limit of the sensor dynamic range. The initial protein structures were generated using AlphaFold2, revealing two key structural differences in the linker regions of RGEPO1 and RGEPO2 that were maintained and stable in the MD (Supplementary Figure 10). Specifically, in RGEPO2, residues N298 and P299 exhibit a tendency towards an open conformation, which results in the loss of beta-sheet character below residue P299 (Supplementary Fig. 11). This structural variation renders the phenolate end of the chromophore in RGEPO2 more accessible to the solvent as compared to RGEPO1. This could explain some of the differences in molecular brightness in K^+^-bound states between them since the fluorescence mechanism likely requires a structural shift upon K^+^ binding to stabilize the phenolate and emissive form. RGEPO2, therefore, would require a larger conformational change as compared to RGEPO1, though they have similar recognition mechanisms, as discussed further below. The MD simulations of RGEPO1 revealed two distinct binding pockets for K^+^ ions (Fig. 4a). The first, referred to as the main binding pocket, was observed in both 20 mM and 300 mM ionic concentrations and exhibited significant K^+^ binding duration (∼140 ns; Fig. 4a upper inset). The second pocket, termed the transient binding pocket, showed a shorter binding time (∼100 ns) and was observed only at 20 mM (Fig. 4a lower inset). The main binding pocket was formed by 13 amino acids, including K240/E241/P242/V243/D294/L295/S297/A298/C299/V300/S301/V349/D350/R374Among them, D294 and D350 directly coordinate potassium ions via ionic interactions. R374 is highly mobile prior to K^+^ binding, and subsequently forms a key hydrogen bond with D294 that stabilizes K^+^ in the pocket for a significant amount of time. The transient pocket consisted of only 5 amino acids, namely D156/V158/K159/S228/K229. Among them, only V158 and K159 were shared with the binding pocket identified in the Ec-Kbp domain of the KRaION1 biosensor^14^.

**Figure 4.**
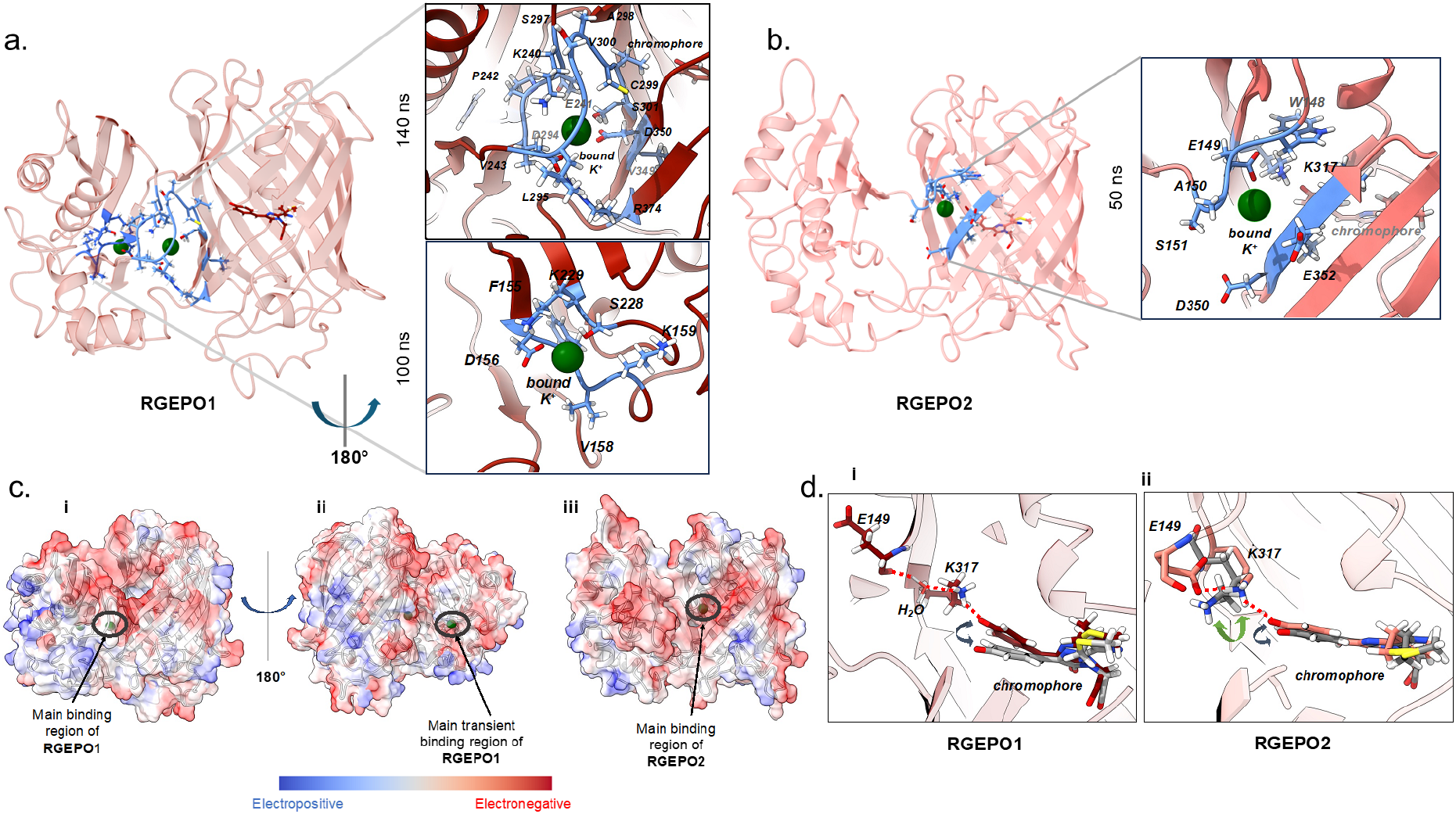
Molecular mechanism of potassium sensitivity. (a) The predicted 3D structure of RGEPO1 is shown in red, with K^+^ ions in green. The residues around the binding pockets are depicted as blue sticks. The binding regions are enlarged on the right with labels for the respective binding residues (upper main binding site and lower transient site). (b) The predicted 3D structure of RGEPO1 is shown in red, with K^+^ ions in green. The residues around the binding pockets are depicted as blue sticks. The binding region is enlarged on the right, and labels are placed for the respective binding residues. (c) Charge distribution of the surface of RGEPO1 (i and ii), RGEPO2 (iii). Surface color is assigned such that red is more electronegative and bluer is more electropositive. The green spheres represent the K^+^ ions. (d) Sensitivity mechanism of (i) RGEPO1 and (ii) RGEPO2. Left, the prominently observed hydrogen bonding network with O^-^ of phenolate ring of RGEPO1 upon K^+^ binding. Right, the prominently observed hydrogen bonding interactions with O^-^ of phenolate ring of RGEPO2 upon K^+^ binding. The hydrogen bonds are represented by thin red dashed lines, indicating the interaction points between the phenolate ring and surrounding residues/water (in sticks), and are labeled with their respective names. The chromophore’s conformational space in both (i) and (ii) is depicted by grey sticks and double-headed arrows, indicating the range of movement in the absence of bound K⁺. In the right panel (ii), the conformation of K317 in the absence of K⁺ is highlighted in grey sticks.

In contrast, RGEPO2’s MD simulations identified only one binding pocket under both ionic conditions (20 mM and 300 mM), in a similar location to the main binding pocket for RGEPO1 but all different amino acids in the binding pocket, which included W148/E149/A150/S151/D350/E352/K317 (Fig. 4b). The binding region is located closer to the fluorescent protein (Fig. 4b). The bound ion remained stable for ∼50 ns in this binding pocket. This positioning also caused an opening between two beta sheets near the phenolate O^-^ end, destabilizing the fluorescent protein’s beta-barrel structure and exposing the inner residues more to the solvent as compared to RGEPO1. This shorter timescale is consistent with the slightly lowered affinity of RGEPO2 observed experimentally in mammalian cells. The main K^+^-binding pockets are located either on the interface between the binding domain and FP in the case of RGEPO1 or in the β-barrel of FP next to the phenolate group of the chromophore. These binding pockets were not previously reported for any other single-FP-based biosensors that utilize other ion-binding domains. However, it was possible to engineer GFP-like FPs to have Zn^2+^ or Cu^2+^ binding pockets inside of β-barrel directly interacting with the chromophore^35^.

Further analysis of RGEPO1’s surface charge distribution revealed an electronegative region extending from the main binding site to the transient site, indicating a higher likelihood of retaining positively charged ions. The MD shows K^+^ ions transiently bound to this red region for a brief time (Fig. 4c, Supplementary Fig. 12). The more grooved areas, such as the main binding pocket and the transient binding regions, provide increased ion stability due to reduced solvent exposure, resulting in longer ion retention. Comparatively, RGEPO2’s surface lacked a continuous negatively charged region. Instead, it had more negatively charged residues clustered around the main binding pocket, forming a groove that stabilized the cations in that specific area (Fig. 4c). Collectively, these results suggest that RGEPO1 has a better performance as a fluorescent sensor than RGEPO2, which is consistent with our experimental results.

Even though RGEPO1 and RGEPO2 have completely different main binding pockets, further analysis of the MD results confirmed that both show the same mechanism related to fluorescence sensing. Upon binding of K^+^ to the main binding pocket of both RGEPO1 and RGEPO2, we observed a chain of hydrogen bonding interactions occurring with the O^-^ position of the phenolate ring of the chromophore. In RGEPO1, these interactions extend from the phenolate ring to K317 and through a water molecule to E149 (Fig. 4d and Supplementary Fig. 13). In RGEPO2, the same interactions occur without the solvent mediating the chain (Fig. 4d). The H bonding residence time between phenolate O^-^ and K317 is approximately 80% and 40% of the trajectory for RGEPO1 and RGEPO2, respectively. The analysis of variation of distance between K317 at the sidechain N atom and chromophore phenolate O^-^ atoms upon binding of K^+^ to the proposed main and transient sites that the distance is lowered (<3 Å) and hydrogen bonding between K317 and O^-^ of chromophore is more stabilized compared to unbound situations (Supplementary Fig. 14). Additionally, analysis of the water shell around the phenolate O^-^ end of all the systems shows a drop in the number of water molecules upon K^+^ binding to the binding domain (Supplementary Fig. 15).

To further investigate the binding regions, we examined the structure of the previously reported NMR-resolved K^+^-binding site in Ec-Kbp of the KRaION1 biosensor (PDB ID: 7PVC). The electrostatic distribution of the binding site clearly differed from that of RGEPO1 and RGEPO2 (Supplementary Fig. 16), suggesting that the binding regions were altered by the introduced mutations. Furthermore, the electronegative charge distribution introduced by mApple at the interface of the potassium binding domain and the fluorescent protein compared to that of green fluorescent protein (PDB ID: 7VCM) made the proposed binding sites more accessible and stable in RGEPO1 and RGEPO2. To determine if the previously reported Kbp binding pocket was still a possibility, we carried out MD simulations by placing the K^+^ ion at this previously reported Kbp binding pocket. In RGEPO1, the bound ion remained stable in this region throughout the simulation, although another K^+^ ion diffused freely to the proposed main binding pocket. The proposed transient pocket of RGEPO1 is located close to the previously reported site (∼7 Å) (Supplementary Fig. 16d), but importantly neither induces the necessary conformational change in the chromophore and K317 (Figure 4d). Additionally, in RGEPO2, the K^+^ binding at the previously reported site was unstable, lasting less than 20 ns. Analysis of the distance between the K317(NZ) atom and the O^-^ atom of the chromophore shows no significant difference between the bound and unbound states (Supplementary Fig. 14f). Taken together, this suggests that the observed main binding pockets in RGEPO1 and RGEPO2 are more likely to be involved in the fluorescence mechanism.

## Recording potassium transients in primary astrocyte and neuronal cultures

Potassium plays a crucial role in maintaining normal neuronal physiology^4,36,37^. Elevated extracellular concentrations of potassium can generate neuronal hyperexcitability^4^. For example, increased extracellular potassium chloride is extensively used to induce membrane depolarization in cultured neurons^38,39^. To investigate whether RGEPO1 could detect extracellular K^+^ dynamics in neurons, in combination with GCaMP6f for intracellular calcium imaging, we expressed RGEPO1 on the extracellular side of the plasma membrane in primary hippocampal neurons using the Asp leading peptide, which displayed a clear membrane localization (Fig. 5a; see Supplementary Fig. 17 for more representative images). Simultaneously, GCaMP6f was expressed in the cytoplasm (Fig. 5a). Transient stimulation with 30 mM K^+^ resulted in a robust increase of approximately 63% in RGEPO1 and 630% in GCaMP6f, with time to peak fluorescence of approximately 33s and 6s, respectively, both of which returned to baseline following K^+^ washout (Fig. 5b).Previous studies have demonstrated that the excessive stimulation of glutamate receptors by glutamate causes a significant efflux of potassium ions from neurons^15,40,41^. We examined whether these changes in intracellular K^+^ could be optically detected using RGEPO2. RGEPO2 was expressed in primary mouse neurons and demonstrated even distribution throughout soma and dendrites (Fig. 5c; see Supplementary Fig. 17 for more representative images). The addition of 500 µM glutamate triggered a robust fluorescence decrease of approximately 77% of RGEPO2, indicating potassium efflux with time to peak of 140 s (Fig. 5d; for visualization of fluorescence changes see Supplementary Movie 3). These results collectively suggested that RGEPOs are promising for monitoring the transient dynamics of intracellular and extracellular potassium in cultured hippocampal neurons.

**Figure 5.**
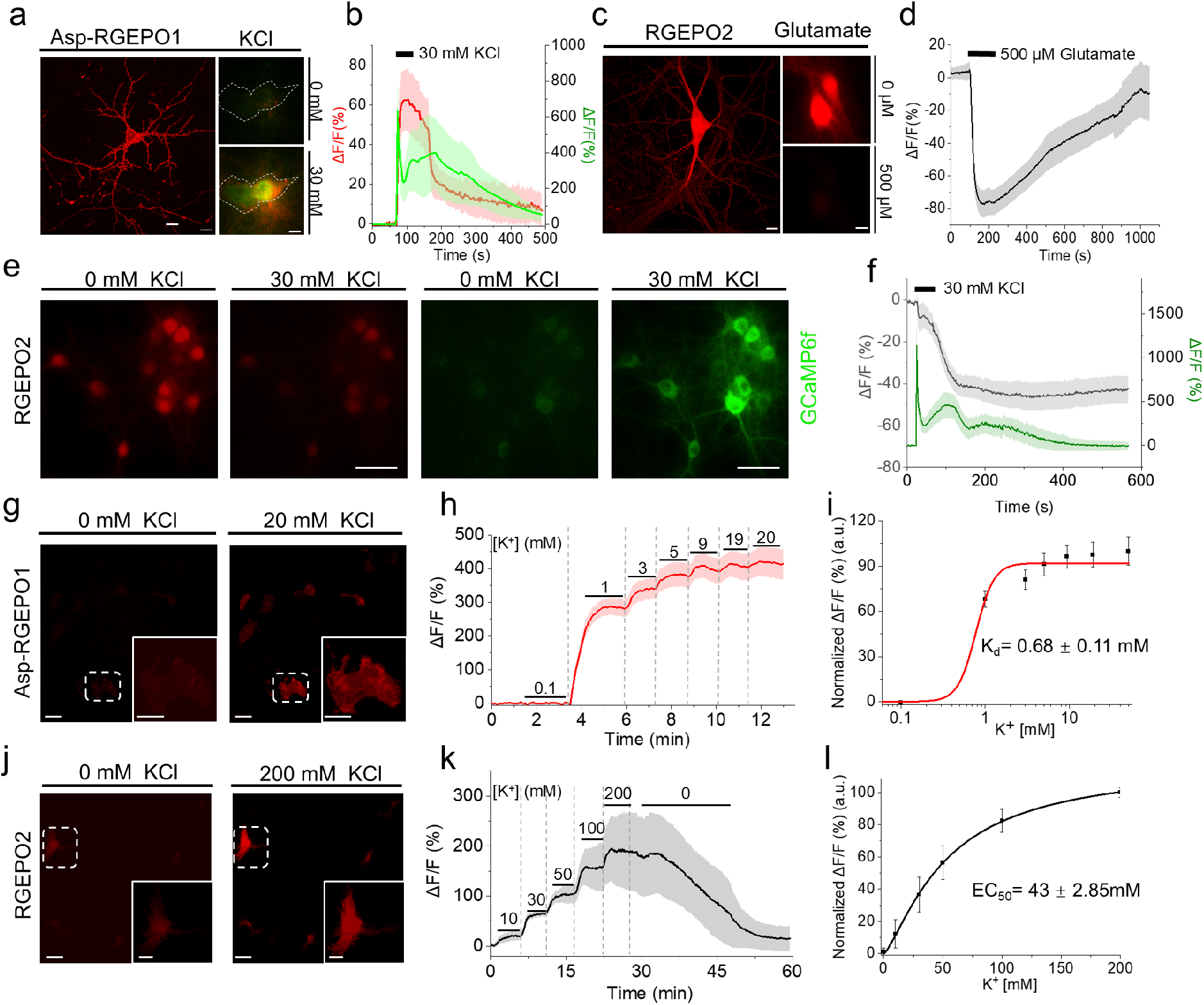
Recording potassium transients in primary neuronal and astrocyte cultures. a. Representative image of hippocampal neurons expressing Asp-RGEPO1, shown in ACSF buffer and in response to 30 mM KCl stimulation (n =6 neurons from 2 independent cultures). b. Time course of fluorescence intensity change of Asp-RGEPO1 and GCaMP6f with stimulation of 30 mM KCl on neurons (n = 13 neurons from 2 wells of 1 transfection), data are expressed as mean ± SD. c. Representative image of a hippocampal neuron expressing RGEPO2, shown in ACSF buffer and in response to 500 μM glutamate (n =19 neurons from 2 independent cultures). d. Time course of fluorescence intensity change of RGEPO2 with stimulation of 500 μM Glutamate (n = 57 neurons from 2 wells of 1 transfection). e. Representative dual-color imaging of K^+^ and Ca^2+^ dynamics in dissociated hippocampal neurons with response to 30 mM KCl. f. Time course of fluorescence intensity changes of RGEPO2 and GCaMP6f with stimulation of 30 mM KCl in neurons (n = 31 neurons from 1 culture, see the other culture data in the supplement Fig.10), data are expressed as mean ± SD. g. Representative fluorescence images of primary astrocyte expressing RGEPO1 in response to 20 mM KCl (n= 25 astrocytes from 2 independent cultures). Scale bars, 100 µm, insert, 20 µm. h. Time course of fluorescence intensity change of RGEPO1 with stimulation by a series of KCl buffers on astrocytes (n= 6 astrocytes from 1 culture), data are expressed as mean ± SD. i. Plot of normalized ΔF/F against different K^+^ concentrations fitted using nonlinear fitting (Hill) for the data shown in panel h. j. Representative fluorescence images of primary astrocyte expressing RGEPO2 in response to 200 mM KCl (n= 17 astrocytes from 2 independent cultures). Scale bars, 100 µm, insert, 10 µm. k. Time course of fluorescence intensity change of RGEPO2 with stimulation by a series of KCl buffer (n= 5 astrocytes from 1 culture), data are expressed as mean ± SD. l. Plot of normalized ΔF/F against different K^+^ concentrations fitted using nonlinear fitting (Hill) for the data shown in panel j.

To explore the simultaneous imaging of K^+^ and Ca^2+^ dynamics in the cytoplasm, we co-expressed RGEPO2 and the green Ca^2+^ indicator GCaMP6f^42^ in cultured hippocampal neurons (Fig. 5e). Brief exposure to 30 mM KCl resulted in a rapid increase in the green fluorescence of GCaMP6f by approximately 1139% due to Ca^2+^ influx induced by neuronal depolarization (Fig. 5f green trace; see Supplementary Fig. 18 for more representative traces and Supplementary Movie 4 for visualization of fluorescence changes). Concurrently, the red fluorescence of RGEPO2 underwent a decrease of approximately 47% (Fig. 5f gray trace; see Supplementary Fig. 18 for more representative traces and Supplementary Movie 4 for visualization of fluorescence changes), suggesting K^+^ efflux.

Compared to neurons, astrocytes are essential for regulating and maintaining stable extracellular potassium levels, ensuring the overall homeostasis of the central nervous system^5,43^. To evaluate the functionality of RGEPOs, we transiently expressed them in primary mouse astrocytes (Fig. 5g-l). RGEPOs displayed clear localization on the extracellular membrane and in the cytoplasm of the astrocytes (Supplementary Fig. 17). The fluorescence increase of extracellularly expressed RGEPO1 was dependent on K^+^ concentration, achieving a ΔF/F of 408% at 20 mM (Fig. 5g,h; for visualization of fluorescence changes see Supplementary Movie 5). The dissociation constant (Kd) at the cell surface was determined to be 0.68±0.11 mM using local K^+^ application with a micropipette, with a Hill coefficient of 1.5 ± 0.36 (Fig. 5i). Similarly, the fluorescence intensity changes of intracellularly localized RGEPO2 were also dependent on K^+^ concentrations, resulting in a ΔF/F of 196% at 200 mM with an EC50 of approximately 43 ± 2.85 mM and a Hill coefficient of 1.33 ± 0.06. (Fig. 5j,k,l; for visualization of fluorescence changes, see Supplementary Movie 6). The observed changes in RGEPO2 were consistent with a previous study using K^+^ fluorescent dye^44^. Additionally, we investigated whether ionophores could also enhance potassium uptake in astrocytes. In the presence of valinomycin and CCCP, the fluorescence change of RGEPO2 increased to 480% with an EC50 of approximately 18 mM and a Hill coefficient of 0.91± 0.19 (Supplementary Fig. 19). The observed difference in affinities under these two conditions can be attributed to the role of valinomycin in facilitating K^+^ influx. This explains why, at the same titration concentrations, the valinomycin-treated group exhibited a larger fluorescence increase, particularly at the initial titration stages, resulting in higher affinity. Moreover, compared to HEK293FT cells, the fluorescence change in astrocytes was 3.5-fold larger at 200 mM K^+^ in the presence of valinomycin and CCCP. This further demonstrates the superior ability of astrocytes to regulate potassium ion concentrations. Collectively, these results suggest that elevated extracellular K^+^ concentration triggers K^+^ uptake by astrocytes, supporting the notion that astrocytes play an important role in involving potassium spatial buffering and thus exhibit differential responses to potassium relative to neurons under similar depolarized conditions.

### Recording potassium transients in neurons in acute brain slices

Encouraged by the high performance of RGEPOs in cultured neurons, we extended our investigation to intact brain tissue. To simultaneously monitor K^+^ dynamics and neuronal activity, we co-expressed Asp-RGEPO1 or RGEPO2 with GCaMP6f in the cerebral cortex neurons of P0 mice, using recombinant adeno-associated virus (rAAV2/9) driven by the human synapsin 1 (hSyn) promoter (Fig. 6a). After 3 to 4 weeks, we prepared acute brain slices and imaged the fluorescence responses to application of extracellular K^+^ (Fig. 6a). Bright fluorescence was detected in the cortex region of the brain slices, confirming the expression of RGEPO1 or RGEPO2 and GCaMP6f (Fig. 6b,c; Supplementary Fig. 20, 21). Stimulation with 20/30 mM KCl elicited a robust and repeatable fluorescence increase of 6.5% for RGEPO1 and 104% for GCaMP6f, both of which returned to baseline after KCl washout (Fig. 6d,e,f). Conversely, the application of 20/30 mM KCl led to a considerable decrease in RGEPO2 fluorescence by 7.0%, accompanied by a corresponding increase in the GCaMP6f signal of 128% (Fig. 6g,h,i). These trends were consistent with the results obtained from cultured neurons, although the ΔF/F values observed in acute brain slices were smaller. This discrepancy is likely due to differences in protein expression levels and the more complex regulatory networks present in intact tissue. Overall, these findings demonstrate that RGEPOs are capable of effectively detecting K^+^ dynamics in both extracellular and intracellular environments in intact brain tissue.

**Figure 6.**
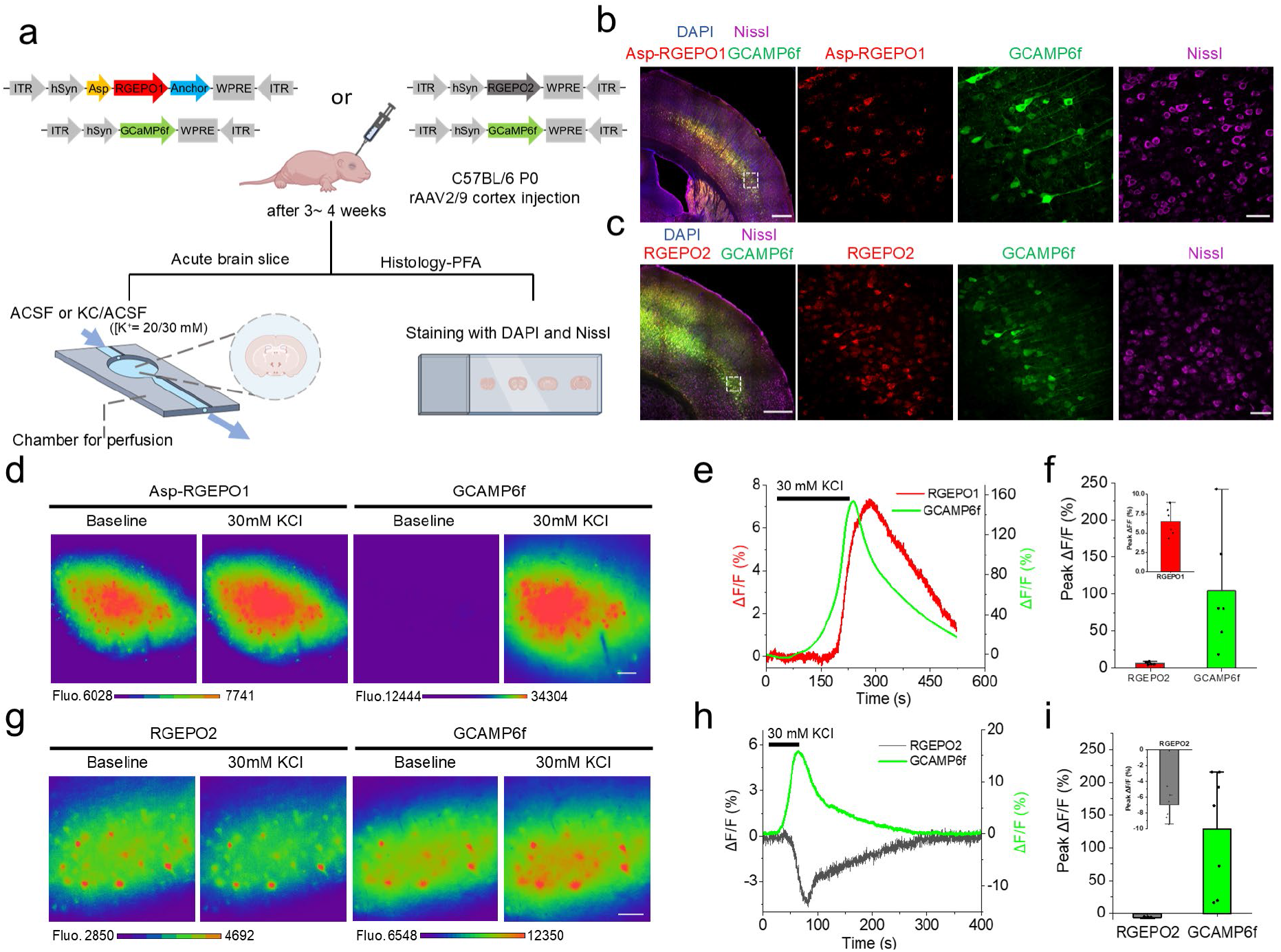
Imaging of RGEPOs in neurons in acute brain slices. a. Schematic illustration depicting the AAV injection of Asp-RGEPO1 or RGEPO2 with GCAMP6f into the cortex for the brain slice experiments. ITR inverted terminal repeat, hSyn human Synapsin I promoter, WPRE woodchuck hepatitis virus posttranslational regulatory element. b. Left, fluorescence image of the cerebral cortex showing the expression of Asp-RGEPO1 (red) and GCAMP6f (green). Scale bar, 500 µm. Right, large magnification of fluorescence images showing relative expression regions. Scale bar, 50 µm. c. Left, fluorescence image of the cerebral cortex showing the expression of RGEPO2 (red) and GCAMP6f (green). Scale bar, 200 µm. Right, large magnification of fluorescence images showing relative expression regions. Scale bar, 50 µm. d. Representative pseudocolor images of RGEPO1 and GCAMP6f expressed in the brain slice in response to 30 mM KCl (n= 4 slices from 2 mice). Color bars, fluorescence intensity. Scale bar, 50 µm. e. Single-trial optical traces of Asp-RGEPO1 and GCAMP6f fluorescence upon stimulation of 30 mM KCl of acute brain slice. f. Quantification maximal ΔF/F of Asp- RGEPO1 and GCAMP6f in response to 20/30 mM KCl (n= 6 slices from 4 mice). Dot, individual data point for each acute brain slice; bar, mean; error bar, SD. g. Representative pseudocolor images of RGEPO2 and GCAMP6f expressed in the brain slice in response to 30 mM KCl(n= 2 slices from 2 mice). Color bars, fluorescence intensity. Scale bar, 50 µm. h. Single-trial optical traces of RGEPO2 and GCAMP6f in response to 20/30 mM KCl. i. Quantification maximal ΔF/F of RGEPO2 and GCAMP6f in response to 30 mM KCl (n= 7 slices from 5 mice). Dot, individual data point for each acute brain slice; bar, mean; error bar, SD.

### Dual color neocortical imaging of Ca^2+^ and K^+^ dynamics during seizure

Epilepsy is a severe neurological disorder marked by defects in potassium channels and disrupted potassium homeostasis, leading to brain hyperexcitability and seizures^45,46^. Previous research using ion-selective probes has documented changes in potassium levels during seizures, but these methods are not scalable to monitor neuronal populations^47–50^. Building on the reliable detection of neuronal activity indicated by the potassium sensors in acute brain slices, we next explored the potential of monitoring potassium dynamics in neuronal populations using Asp-RGEPO1 and RGEPO2 *in vivo* in mice. In this experiment, we co-expressed GCaMP6f with RGEPO variants in neurons of the primary somatosensory cortex and employed two-photon microscopy to capture seizure activities in L2/3 neurons elicited by kainic acid (Fig. 7a,b). Similar to previous findings^51–53^, we observed a synchronous fluorescence wave followed by a propagating wave across the board in the case of calcium (Fig 7c-e, Supplementary movie 7,8), which should indicate the stage of seizure activity and spreading wave of depolarization, respectively. In mice expressing Asp-RGEPO1, we detected a spreading wave of fluorescence increase (median peak ΔF/F = 89%). The signal onset (Fig. 7d,g) for the RGEPO1 wave showed a broader wavefront compared to GCaMP6f, likely due to the slower RGEPO1 kinetics and, thus, varied peaking of RGEPO1. However, analysis of the averaged response from neuronal cell bodies revealed that this wave rose nearly simultaneously with, but approximately 2.5 seconds ahead of, the propagating wave observed with GCaMP6f (median peak ΔF/F 1411%; Supplementary Fig. 22a, Supplementary Movie 7). Together, this observation suggested a significant rise in extracellular potassium following seizure termination. For RGEPO2, we observed a rapid fluorescence decay (median peak ΔF/F = -37%) coinciding with the seizure onset wave seen in GCaMP6f (median peak ΔF/F 136%; Fig. 7e). RGEPO2 also exhibited a subsequent spreading wave (Fig. 7e-h). Due to acute z-plane motion observed in both channels during seizures, the exact latency between the calcium and potassium spreading waves cannot be determined from averaged traces of the cell population (Supplementary Fig. 22b). Nonetheless, the RGEPO2 wave propagates within a similar time frame to GCaMP6f, as evidenced by approximate onset of the signal (Fig. 7e, Supplementary Movie 8) and the delay of fluorescence peaking time in individual cells (Fig. 7h). These findings indicate a decrease in intracellular potassium levels reported by RGEPO2 during seizures. Compared to the results obtained from the acute brain slice data, RGEPOs in the mouse seizures model exhibited similar trends but produced stronger responses. Overall, these results demonstrated the capability of RGEPO1 and RGEPO2 to report potassium activity during seizures in vivo in mice. Although they offer lower temporal resolution and signal amplitude compared to GCaMP6f, RGEPO1, and RGEPO2, they enable the observation of potassium dynamics with single-cell resolution at the population level *in vivo* in drug-induced seizures for the first time.

**Figure 7.**
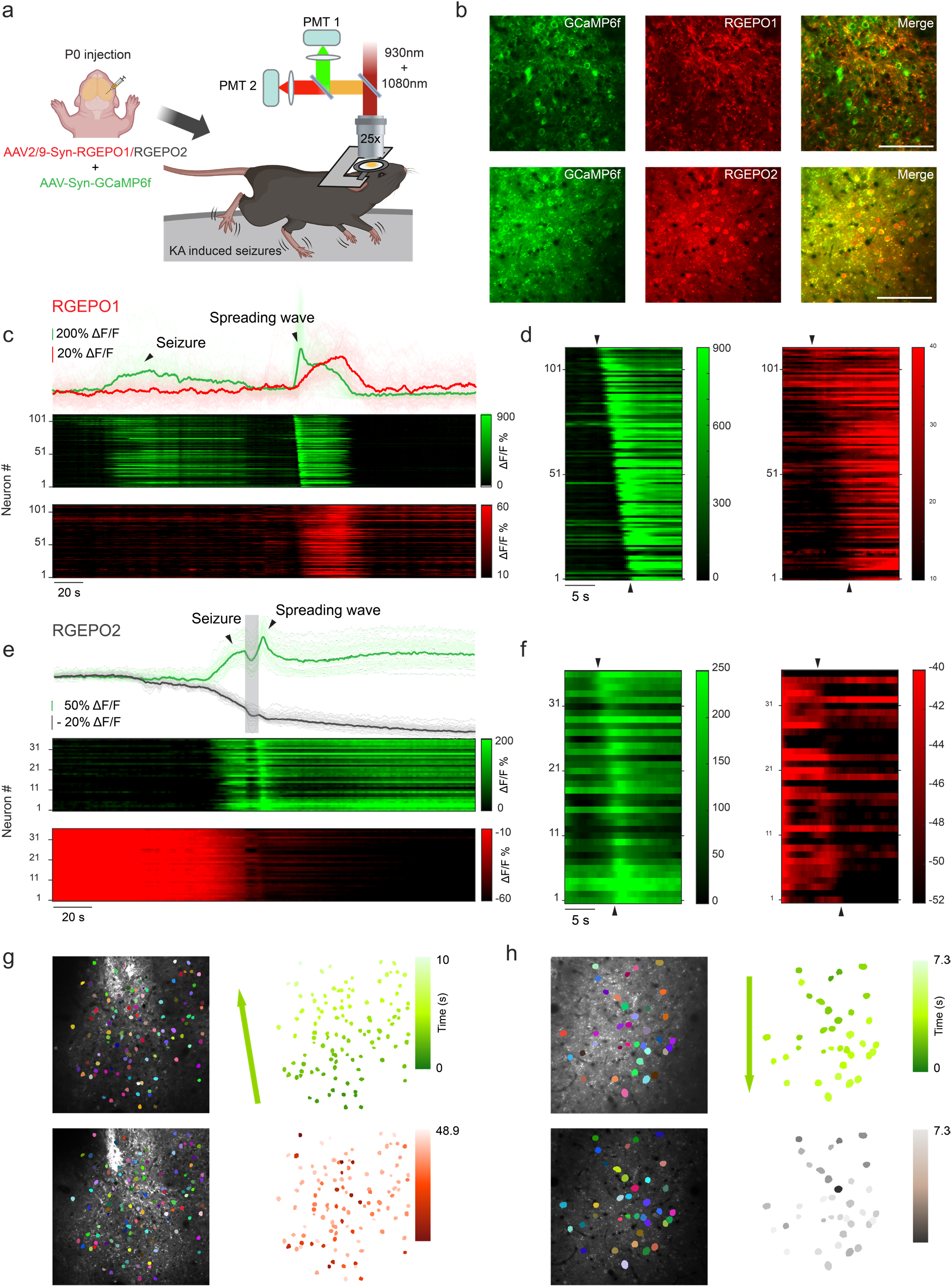
Two-photon imaging of GCaMP6f and RGEPOs in mice with KA-induced seizures. a. Schematic representation of virus injection and imaging in a mouse undergoing KA-induced seizures using a two-photon microscopy. b. Representative single-plane images showing co-expression of RGEPO1/2 and GCaMP6f in layer 2/3 neurons of the primary somatosensory cortex. c. Top panel: Individual (light green and light red) and averaged (green and red) fluorescence responses of GCaMP6f and RGEPO1 during KA-induced seizures and subsequent spreading waves. Middle and bottom panels: Vertical projections of fluorescence profiles from neurons expressing GCaMP6f and RGEPO2 during KA-induced seizures and subsequent spreading waves. d. Zoomed-in view of the vertical line profile of GCaMP6f and RGEPO2 from c, highlighting the onset of the spreading wave. n = 111 neurons from one mouse. e. Top panel: Individual (light green and light red) and averaged (gray and dark gray) fluorescence responses of GCaMP6f and RGEPO2 during KA-induced seizures and subsequent spreading waves. Middle and bottom panels: Vertical projections of fluorescence profiles from neurons expressing GCaMP6f and RGEPO2 during KA-induced seizures and subsequent spreading waves. The gray shaded box indicates the period of acute motion observed along the z-axis. f. Similar to panels d, but for GCaMP6f and RGEPO2. n = 36 neurons from one mouse. g. Temporal analysis of the spreading wave of GCaMP6f and RGEPO1 following seizure activity. Left: pseudocolored mask of individual neurons overlaid on the average projection image. Right: same mask as left, colored based on fluorescence peak time. Statistics are the same as in panel c. h. Similar to panel f, but for GCaMP6f and RGEPO2. Statistics are the same as in panel e. Colored arrow indicates the approximate direction of wave propagation. Scale bar: 100 µm.

## Discussion

In this study, we engineered the first red genetically encoded fluorescent potassium indicators, RGEPO1 and RGEPO2, by incorporating the Hv-Kbp, a novel potassium-binding domain, into the mApple fluorescent protein (Supplementary Fig. 7). Compared to green fluorescent potassium sensors^13–15^ (Supplementary Table 1), RGEPOs offer several advantages, including compatibility with multicolor imaging and low phototoxicity and the ability to be applied in intact tissue environments. Additionally, RGEPOs exhibited a sensitive response to the intracellular and extracellular K^+^ change in the culture neurons and astrocytes, acute brain slices, as well as in awake mice. Our results collectively suggest that RGEPOs are effective tools for detecting intracellular and extracellular potassium during the neuronal activity process.

To engineer RGEPOs, we inserted the potassium binding domain into the red fluorescent protein mApple at position 151 on the 7^th^ β-strand next to the chromophore (Fig. 1c). Majority of the previously reported red genetically encoded fluorescent probes have been based on the fusion of circularly permuted fluorescent protein (cpFP) with binding domain^54,55^. A similar molecular design has been used to develop glutamate and calcium biosensors, which were also constructed by inserting corresponding binding domains into the 7^th^ β-strand of mApple^20,56^. However, the applicability of these biosensors was not demonstrated beyond cultured mammalian cells. In contrast, RGEPOs displayed a strong response in the cultured neurons and astrocytes (Fig. 5) and were also capable of detecting K^+^ dynamics in both extracellular and intracellular environments of intact brain tissue (Fig. 6). This opens new avenues for the development of red fluorescent probes, making RGEPOs a successful example of building red fluorescence protein-based indicators using the inserted-into-red FP topology.

RGEPOs were optimized through a combination of directed molecular evolution in *E. coli* and mammalian cells (Fig. 1,2). This dual-system approach addresses the common challenge of transferring biosensor prototypes optimized in *E. coli* to mammalian cells^57^, where they often perform suboptimally. By initially using the *E. coli* system combined with random mutant generation for high-throughput screening, we were able to identify promising prototypes, which were then validated in mammalian cells. Compared to optimization in a single system alone, this combined optimization strategy increases the throughput of sensor screening while overcoming the issue of prototype incompatibility between systems, thereby enhancing overall efficiency.

RGEPOs were successfully employed to detect intracellular and extracellular potassium dynamics in cultured neurons and astrocytes, as well as in acute brain slices. RGEPO1 exhibited sensitive responses to extracellular K^+^ across different models. RGEPO2 showed a negative response in primary neurons and a positive response in astrocytes, consistent with their physiological functions^4,5,43^ (Fig. 5). Furthermore, we observed that cultured neurons took longer to return to baseline after washing out K^+^ compared to astrocytes (Fig. 5f, k), which support that astrocytes have a stronger capacity to regulate potassium ions^5,43^. This finding aligns with the understanding that while both neurons and astrocytes depend on K⁺ channels, the types and roles of these channels differ significantly^36,58^. In acute brain slices, we observed a similar response pattern to that seen in cultured neurons, though the response amplitudes were less pronounced, indicating the more complex K^+^ dynamics in the actual physiological environment.

To further extend the application of our potassium probes, we examined RGEPOs in vivo in mice using two-photon imaging. In mice with kainic acid (KA)-induced seizures, we observed a spreading wave in both RGEPO1 and RGEPO2, with delay time of wave arrival at cells observed across the board (Fig. 7c-h, Supplementary Movie 7, 8). Interestingly, a previous study using GCaMP and ion-selective microelectrodes reported a latency in the rise of Ca^2+^ fluorescence relative to the increase in extracellular K^+^ during KCl-induced spreading waves^59^. In our study, by fully adopting an optical approach, we found consistency with that study, but at a population level: the spreading wave detected by GCaMP6f occurred 2.5 seconds after that of RGEPO1. This indicates that RGEPO1 can reliably report potassium dynamics during spreading waves, similar to those demonstrated previously by electrophysiology. It is known that both seizure activity and the subsequent spreading wave of depolarization drive strong calcium influx and potassium efflux intracellularly and lead to a significant rise in extracellular potassium levels^60–62^. We found that RGEPO2 is capable of sensing and reporting intracellular potassium drops during both seizure activity and the spreading depolarization stage. While we did not observe a significant increase in RGEPO1 signal upon Ca^2+^ wave onset during seizures, this may be limited by the current sensitivity level of the probe. In addition, the suboptimal sensitivity and kinetics of RGEPOs observed in acute brain slices and in mice *in vivo* also suggest opportunities for improving RGEPOs in future research. In summary, our results demonstrated the capability of RGEPO1 and RGEPO2 to capture key physiological changes in potassium levels with single-cell resolution, which, to our knowledge, is the first report of optical measurements of potassium transients during seizure-related activities in vivo in mice. Through molecular dynamics simulations, we gained insights into the molecular mechanisms underlying potassium binding within RGEPO1 and RGEPO2. Our simulations revealed distinct potassium-binding pockets in the two indicators. RGEPO1 was found to have two potassium-binding pockets: a main binding pocket and a transient pocket similar to the potassium-binding site in Ec-Kbp. RGEPO2 has one potassium binding pocket, which is close to the chromophore. These findings suggest that linker content changes can impact substrate binding, especially in proteins with simpler structures. The MD simulations also indicated that RGEPO1’s surface is more electronegative, facilitating the retention of positively charged ions and supporting the notion that RGEPO1 performs better as a fluorescent sensor than RGEPO2, consistent with our solution-based results.

The primary limitations of RGEPOs lie in their relatively low dynamic response range and limited fluorescence change (ΔF/F) within intracellular environments, particularly in intact brain tissue. This issue was notably pronounced with RGEPO2, which exhibited lower ΔF/F in HEK293FT cells and neurons compared to green fluorescent potassium indicators. A key factor contributing to this reduced performance is likely the high basal potassium concentration in the cytoplasm, which attenuates sensor sensitivity. To address these limitations, in our future studies, we plan to develop RGEPO variants with greater sensitivity and optimized binding affinity for improving the dynamics range. Moreover, we are working on engineering next-generation RGEPO variants with faster K^+^ kinetics, enhancing their ability to detect K^+^ pulses more effectively. Furthermore, we also observed puncta formation at high expression levels of RGEPOs, which we aim to mitigate by optimizing expression levels and binding domains. Finally, future efforts will focus on two-photon imaging experiments in awake mice to track cellular and subcellular K^+^ dynamics under physiological and pathological conditions.

In summary, we have developed the first red genetically encoded indicators for imaging the intracellular and extracellular potassium dynamic in cultured mammalian cells, acute brain slices, and mice *in vivo*. RGEPOs represent significant advancements in the field of potassium imaging, offering new tools for the real-time visualization of K^+^ dynamics in both extracellular and intracellular environments. Future work will focus on enhancing their sensitivity and expanding their applicability to other biological systems, ultimately contributing to a deeper understanding of potassium’s role in cellular physiology.

## Methods

### Genome mining for the Ec-Kbp homologous domains

The BLASTp tool from the National Center for Biotechnology Information (NCBI) was utilized to identify alternative potassium-binding domains. We initiated our search with the amino acid sequence of Hv-Kbp (NCBI Sequence ID: VAV91021.1) as the query, targeting the env_nr metagenomic database. The search was executed with default parameters aside from the specified database. The acquired results were filtered based on the percentage similarity of amino acid sequences (above 40%), and manual selection was applied to refine the results. The following proteins were selected for further investigation: VAV (GenBank: VAV99455.1; hypothetical protein from hydrothermal vent metagenome); GAI (GenBank: GAI29416.1; unnamed protein product from marine sediment metagenome); MND (GenBank: MND59673.1; LysM domain/BON superfamily protein from compost metagenome); MPM (GenBank: MPM71561; putative protein YgaU from bioreactor metagenome); MNT (GenBank: MNT23082.1; LysM domain/BON superfamily protein from compost metagenome).

### Molecular cloning and mutagenesis

The DNA sequences encoding identified potassium binding domains, MNT (list all of them; sequence IDs above), and KRaION1 were synthesized by Tsingke Biotech Co., Ltd (China) and swapped with Electra1 gene in the pHybrid expression vector^63^ (pHybrid-Electra1, WeKwikGene plasmid #0000202) for screening in *E.coli* bacteria and cytoplasmic expression in HEK cells. The secretion peptides used in Fig. 2d were synthesized de novo with mammalian codon optimization based on the amino acid sequences of the N-termini of aspartic proteinase nepenthesin-1 (Asp, 1-24 aa from Sequence ID: BAF98915.1), azurocidin (Azu, 1-19 aa from Sequence ID: NP_001691.1), corticotropin-releasing factor receptor 1 (CRFR, 1-24 aa from Sequence ID: XP_040124593.1), endothelin receptor type B (EtBR, 1-26 aa from Sequence ID: CAD30645.1). For extracellular-cell-surface expression in HEK cells, the corresponding genes were cloned and swapped with the Thyone gene in the pDisplay-Thyone plasmid (WeKwikGene plasmid #0000366). For generating prototypes of red potassium indicators, the genes of the selected potassium binding domains were swapped with the gene of the CaM-M13 calcium-binding domain in the FRCaMPi gene, preserving the original linkers between FP and binding domains. For expression in primary mouse cell cultures, the RGEPOs genes were swapped with iRFP670-P2A-GFP in the pAAV-CAG-iRFP670-P2A-GFP plasmid (WeKwikGene plasmid# 0000008) and with GRAB_5-HT1.0 in the pAAV-hSyn-GRAB_5- HT1.0 plasmid (WeKwikGene plasmid# 0000123). All DNA fragments and plasmids were sequenced and verified by Healthy Creatures Biotec Co., Ltd (China).

The custom DNA oligonucleotides were synthesized by Tsingke Biotechnology Co., Ltd. PrimeStar Max master mix (Takara, Japan) was used for routine polymerase chain reaction (PCR) amplification. The routine Gibson assembly was conducted using Hieff Clone Plus One-step cloning kit (YEASEN Biotech, China). Site-directed mutagenesis of RGEPOs was performed with overlap extension PCR using custom DNA oligonucleotides with degenerated NNS codons. Random mutagenesis was performed using error-prone PCR (GENSTAR, China) with the mutation rate 1-6 bp per kbp according to the manufacturer’s instructions. Following purification on a 1% agarose gel, PCR products obtained using site-directed or random mutagenesis were enzymatically digested with KpnI and XbaI (New England BioLabs, USA), and ligated with a similarly digested pHybrid backbone vector via T4 DNA ligase (New England BioLabs, USA). Electroporation facilitated the introduction of ligation products into electrocompetent NEB 10-beta cells (10^10^ cfu/µg; Biomed, China), which were then cultured in LB medium supplemented with 0.1 mg/ml ampicillin (Amp^+^) and 0.06% w/v L-Rhamnose (Rha^+^) (Sangon Biotech Co., Ltd., China) for 24 h at 37°C followed by culturing for 18 h at 18°C. The BD FACS Melody™ Cell Sorter (BD Biosciences, USA) was utilized for bacterial library sorting, employing 561 nm excitation wavelength and 585/42 nm filter, from which approximately 30,000 cells with the highest fluorescence were sorted and cultured overnight on Amp^+^/Rha^+^ agar plates at 37°C. Selected bright colonies were subsequently cultured in duplicates of 24 deep-well RB blocks (Invitrogen, USA), utilizing 4 mL of LB medium supplemented with ampicillin for plasmid extraction and another 4 mL of LB medium supplemented with ampicillin and rhamnose for protein expression. The cultures were initially incubated at 37°C for 24 hours, followed by a lower temperature incubation at 18°C for 18 hours. Post three freeze-thaw cycles at -80°C to disrupt the cells, proteins were extracted using the Bacterial Protein Extraction Kit (Sangon Bitech, China). The lysates, titrated with KCl solution in the range of 0 to 200 mM, were assessed for fluorescence response in 96-well plates using the Varioskan LUX Plate reader (ThermoFisher Scientific, USA) with excitation at 560 nm and emission at 580-640 nm. Mutants, selected based on the highest fluorescence changes (ΔF/F) and relative baseline brightness, were further sequenced and revalidated as described above.

### 3. Protein purification and in vitro characterization

Protein expression and purification were carried out utilizing the rhamnose-inducible pHybrid system in TOP10 cells (Biomed, China) cultured in Amp^+^/Rha^+^ LB medium at 37°C overnight followed by a subsequent culturing at 18°C for 18 hours as outlined previously. *E. coli* cells were harvested at 4,500 rpm, 4°C for 20 minutes. The cell pellets were lysed in lysozyme buffer with sonication and then clarified by ultracentrifugation at 18,000 rpm, 4°C for 30 minutes. Next, the supernatants were incubated with Ni-NTA agarose resin (YEASEN Biotech, China) for 20 minutes at 4°C, before being processed through an affinity chromatography column (Beyotime, China) for protein elution. The eluted proteins were dialyzed against K^+^-free 25 mM Tris/MES buffer (pH 7.4) overnight at 4°C and subsequently stored at 4°C until used.

The Absorption spectra of RGEPOs were measured using a UV-Vis-NIR Spectrophotometer (Shimadzu, Japan), covering a wavelength range from 260 to 700 nm. The fluorescence spectra were collected with FLS1000 Spectrofluorometer controlled by Fluoracle spectrometer operating software (Edinburgh Instruments, UK). The Extinction coefficients (EC) were calculated employing the alkaline denaturation method, as previously described^14^. Briefly, the biosensor solution was rapidly mixed with 2M NaOH at a 1:1 volume ratio. The absorption spectrum was then promptly recorded for subsequent comparison with the solution before adding NaOH. The absolute fluorescence quantum yield (QY) and fluorescence lifetime measurements were conducted using the FLS1000 spectrometer (Edinburgh Instruments, UK) which was outfitted with an integrating sphere accessory, strictly adhering to the guidelines provided by the manufacturer.

Rapid kinetic measurements detailing the interactions between RGEPOs and potassium ions were conducted utilizing a Stopped Flow Spectrometer (SX 20, Applied Photophysics Ltd., UK). The fluorescence detection parameters included an excitation wavelength of 560 nm, with a bandwidth of 1 nm, and an emission filter set to 580/10 nm. Fluorescence was recorded for 5000 time points over 250 s in 3 sequential repeats respectively.

To conduct pH titrations, protein solutions were adjusted to various pH levels (from 3 to 10) using Hydrion Buffer Chemvelopes (Hydrion, USA) in the presence and absence of 150 mM KCl. To assess K^+^ specificity, proteins were introduced into buffers containing distinct ions: LiCl (5 mM), NaCl (15 mM), KCl (150 mM), RbCl (5 mM), CsCl (5 mM), NH4Cl (5 mM), MgCl2 (20 mM), CaCl2 (20 mM) and ZnCl2 (10 µM) (all from Sigma-Aldrich, USA). For evaluating K^+^ interactions under non-isotonic conditions, protein solution was diluted into a series of K^+^ buffers ranging from 0 mM to 500 mM within Tris/MES buffers (pH 7.4). For isotonic Kd measurement, protein solutions were added into isotonic solutions characterized by varied ratios of KCl to N-methyl-D-glucamine (NMDG, Sigma-Aldrich, USA), all prepared in Tris/MES buffers (pH 7.4), with K^+^ concentrations examined from 0 to 250 mM. NMDG was employed to maintain consistent osmotic pressure throughout the experiments. The fluorescence responses of these protein solutions were measured using a Thermo Scientific Varioskan LUX spectrophotometer, setting the excitation wavelength at 560 nm and measuring emission between 580 and 640 nm.

### 4. Imaging of RGEPOs in HEK293FT cells

HEK293FT cells (ATCC® ACS-4500™, USA) were cultured in Dulbecco’s modified Eagle’s medium (DMEM, Gibco) supplemented with 10% fetal bovine serum (FBS, CellMAX, China) and 200 U/mL penicillin–streptomycin (YEASEN Biotech, China) at 37°C with 5% CO2. Cells were seeded on 14 mm coverslips (Biosharp, China) pre-coated with Matrigel (Animal Blood Ware, China) in 24-well plates before transfection. Transfection of HEK293FT cells was performed employing Hieff Trans (YEASEN Biotech, China) according to the manufacturer’s protocol. Briefly, 500 ng of plasmids and 1 µl of transfection reagent were added to each well. Imaging was conducted 24-36 h post-transfection, utilizing an inverted wide-field Nikon Eclipse Ti2 microscope, equipped with a SPECTRA III light engine (Lumencor, USA) and an Orca Flash4.0v3 camera (Hamamatsu, USA), and operated via NIS-Elements AR software, utilizing 20x NA0.75 or 10x NA0.45 objective lens.

In the process of intracellular validation, the transfected cells were initially incubated in a basal buffer solution composed of 1.5 mM CaCl2, 1.5 mM MgSO4, 1.25 mM NaH2PO4, 26 mM NaHCO3 and 10 mM D-Glucose (pH 7.4) with 15 μM valinomycin (Aladdin Biochemical Technology Co., Ltd., China) and 5 μM CCCP (Shanghai Macklin Biochemical Co., Ltd., China) for 15 minutes at 22-25°C. Following incubation, cells were subjected to perfusion with a series of potassium-enriched buffers, with concentrations varying from 0 to 200 mM, each prepared in the basal buffer via an automated perfusion system, operating at a flow rate of 1 mL/min. The methodology ensured a stable extracellular environment for the cells, negating the need for manual intervention in applying the solutions, as previously described^14^. For the purpose of extracellular validation, cell culture medium was replaced with basal buffer before prior to imaging. A gradient of potassium ion solutions, with concentrations ranging from 0 to 50 mM, was carefully introduced to the cells using precision pipetting, to assess the effects of varying extracellular potassium levels.

### 5. Culture and imaging in primary neurons and astrocytes

All animal maintenance and experimental procedures for mice were conducted according to the Westlake University animal care guidelines, and all animal studies were approved by the Institutional Animal Care and Use Committee of Westlake University under animal protocol (AP#19-044-KP-2).

Cortical astrocytes and hippocampal neurons were harvested from postnatal day 0 C57BL/6 pups, without distinction of sex, obtained from the Animal Facility at Westlake University. Dissected tissues were digested using 0.25% Trypsin-EDTA (YEASEN Biotech, China) for 10 minutes at 37°C, a process that was subsequently halted by the addition of advance medium (Gibco, USA) containing 10% fetal bovine serum (FBS, Gibco, USA).

The dissociated cortical astrocytes were seeded on 60 mm culture dishes (Thermo Fisher Scientific, USA) pre-coated with Matrigel (Animal Blood Ware, China). Half of the culture medium was refreshed every 3-4 days with DMEM (Gibco, USA) supplemented with 10% FBS (CellMAX, China) every 3-4 days. For transfection procedures, astrocytes were further sub-cultured in 24-well plates (NEST, China), also pre-coated with Matrigel (Animal Blood Ware, China), followed by transfection with the pAAV-CAG-RGEPO1/2 plasmids using Hieff Trans (YEASEN Biotech, China) according to the manufacturer’s instructions.

The dissociated neurons were plated on 14 mm coverslips (Biosharp, China) pre-coated with Matrigel (Animal Blood Ware, China). Neuronal cultures were maintained in Neurobasal-A medium (Gibco, USA) supplemented with 10% FBS (Gibco, USA) and 2% B27 supplement (Gibco, USA), with medium half changes every 2-3 days. At 3 days in vitro (DIV), AraC (0.002 mM, Sigma-Aldrich, USA) was introduced to selectively inhibit astrocytes proliferation. Cultured neurons were transduced with rAAV2/9 virus (rAAV2/9-hSyn-GCaMP6f, titer, >10^12^ viral genome per ml, Shanghai Sunbio Medical Biotechnology; rAAV2/9-hSyn- RGEPO1, titer, >10^12^ viral genome per ml, BrainVTA (Wuhan) Co., Ltd; rAAV2/9-hSyn- RGEPO2, titer, >10^12^ viral genome per ml, Shanghai Sunbio Medical Biotechnology) at 5 DIV. Measurements on neurons were conducted between DIV 10 and 13. The cell culture conditions and imaging setup for astrocytes and neurons followed the protocols outlined in the section describing HEK293FT cells.

### Immunohistochemistry

To characterize the expression of RGEPO1 and RGEPO2 in fixed tissue, the mice at 1-1.5 months old were deeply anesthetized using 1% sodium pentobarbital and transcardically perfused with pre-chilled PBS followed by 4% paraformaldehyde (PFA; Beyotime, China). Brain tissue was harvested and postfixed in 4% PFA for 4-6 h at 4°C. Followed by rinsing in PBS three times, the coronal brain tissue sections at a thickness of 50 μm were sliced using a VT1200S Vibratome (Leica Microsystems). The free-floating slices were incubated with Nissl (dilution ratio 1:1000; Invitogen, USA) diluted in PBS for 20 min at 22-25°C and then washed three times for 5 min each with PBS. Following mounted with prolonged gold antifade reagent (ThermoFisher; USA), the brain slices were imaged at Nikon Spinning Disk Field Scanning Confocal Systems (CSU-W1 SoRa) with 10x and 40x objective lenses.

### Virus injections

The neonatal intraventricular injections of rAAV2/9 (rAAV2/9-hSyn-GCaMP6f, titer, >10^12^ viral genome per ml, Shanghai Sunbio Medical Biotechnology; rAAV2/9-hSyn- RGEPO1, titer, >10^12^ viral genome per ml, BrainVTA (Wuhan) Co., Ltd; rAAV2/9-hSyn- RGEPO2, titer, >10^12^ viral genome per ml, Shanghai Sunbio Medical Biotechnology) were performed for characterization in brain tissue^64^. Briefly, viruses were injected pan-cortically into pups at postnatal day 0, regardless of sex, with a Hamilton microliter syringe. To induce hypothermic anesthesia, the pups were put on foil with ice underneath until they stopped responding to gentle squeezing on the limbs. 0.5 μL virus solution (titer: >10^12^ v.g. mL^-1^) supplemented with 0.1% FastGreen dye (F7252, Sigma-Aldrich) was injected into each hemisphere. The pups were then moved onto a heating pad maintained at 37°C for a 5 min recovery. After regaining response to gentle squeezing, the pups were returned to the home cages. To prevent cannibalism, the whole injection process for each pup was strictly controlled within a few minutes.

### Acute brain slice preparation and imaging

Neonatal AAV injected mice were deeply anesthetized with 1% pentobarbital sodium and then perfused transcardially with cold (2-4°C) dissecting solution containing (in mM): 213 sucrose, 5 KCl, 1.4 NaH2PO4, 26 NaHCO3, 1 CaCl2, 3 MgCl2, and 10 D-(+)-glucose saturated with 95% O2 and 5% CO2, pH 7.4 adjusted with NaOH, 300-310 mOsm/L. After the brain tissue was swiftly trimmed, coronal slices with 350 µm thick were cut off using a vibratome (VT1000S, Leica Microsystems, Germany) and then transferred into a prewarmed chamber (BSC-PC, Harvard Apparatus, USA) for one-hour incubation at 34°C with oxygenated regular aCSF containing (in mM): 124 NaCl, 5 KCl, 1.4 NaH2PO4, 26 NaHCO3, 2.4 CaCl2, 1.2 MgCl2, and 10 D-(+)-glucose saturated with 95% O2 and 5% CO2 (pH 7.4 adjusted with NaOH, 300-310 mOsm/L), followed by at least 1 h recovery at room temperature (21-25°C) before imaging. The imaging setup for acute brain slices followed the protocols described for HEK293FT cells at 22-25°C. All perfusion buffers were saturated with 95% O2 and 5% CO2.

### Virus injection and cranial window implantation

Mice received cortical injection of a mixture of rAAV2/9-hSyn-GCaMP6f and rAAV2/9-hSyn- RGEPO1/2 virus on day P0. The procedure and conditions were similar to that described for acute brain slice imaging. Seven to eight weeks post-injection, the mice were ready for craniotomy. Using a skull drill, a 3.0-mm diameter craniotomy was created above the center of the primary somatosensory cortex (AP: −0.9mm, ML: −3.0mm approximately). The skull was carefully removed with surgical forceps. A coverslip was then implanted onto the craniotomy region and a custom titanium headplate was then secured to the skull with cement. To reinforce the window, a layer of denture base resin was applied over the dental cement and allowed to dry for 10 minutes. The mice were then returned to their home cages and given at least four days to recover post-surgery.

### Two photon in vivo imaging

For two-photon imaging, awake animals were head-fixed onto a spherical treadmill after recovery. Imaging was performed using a commercial Olympus FVMPE-RS microscope with a 25 X water immersion objective lens. To induce seizures, kainic acid (20 mg/kg; cat#487-79- 6, MedChemExpress, USA) was diluted in PBS and injected intraperitoneally into the mice. Functional images (512 × 512 pixels, 0.994 μm/pixel) were acquired at 3 Hz using a resonant scanner to capture fluorescence dynamics. The laser operated at 930 nm and 1080 nm, with emission light filtered through band-pass filter BA495-540 and BA575-645, respectively.

### Molecular dynamics

We carried out classical molecular dynamics (MD) simulations for both RGEPO1 and RGEPO2 systems at different ion concentrations, as described below.

In order to prepare the systems, we used AlphaFold2 version 1.5.5, where the NMR structure of the potassium binding domain (PDBID: 7PVC) and the crystal structure of mCherry (PDB ID: 2H5Q) were used to validate the final result^14,65,66^.After that, MolProbity version 4.5.2^67^ and PDB2PQR web server version 3.5.2^68^ were used to obtain the proper protonation states for protein residues in each system at pH 7. Then we arranged simulations under different conditions as described below.

To run free diffusion simulations in 20 mM ionic concentration conditions for RGEPO1 and RGEPO2, we randomly added 11 potassium ions to neutralize the systems and achieve a 20 mM concentration. We then solvated the systems using TIP3P water model with a 14 Å water pad from the surface of the protein^69^. To generate the topology and coordinate files we used tleap from AMBER20^70^. The force field parameters used were ff14SB for the protein^71^ and custom-made parameters for the chromophore using AutoParams^72^. After heating to 300 K, all systems were equilibrated by restraining the protein and chromophore while leaving the solvent unrestrained. We performed a series of iterative relaxations starting at 100 kcal/mol and incrementally decreasing the restraints towards 0, with the density approaching 1.0 g/cm^-3^. A 200 ns classical MD simulation was then run on the equilibrated system using the Langevin thermostat^73^ with a 5 ps^-1^ friction parameter in the NVT ensemble with pmemd.cuda in Amber20. The trajectories were run an additional two times for 100 ns each to confirm statistical convergence.

To run in 300 mM ionic concentration system simulations, we carried out MD simulations for both RGEPO1 and RGEPO2 with a 300 mM ionic concentration, where we randomly added 38 potassium ions and the appropriate amount of chloride ions to neutralize the system (27). The simulations were conducted following the same procedure as in 20 mM ionic concentration systems.

To run in 300 mM ionic concertation system with bound K^+^ at Kbp site (PDB ID -7PVC), we carried out MD simulations for both RGEPO1 and RGEPO2 with same procedure. The only difference is that one of the randomly added K^+^ ions was placed at the initially reported binding site of Kbp (PDB ID -7PVC) to see the stability and the potential effect on the sensitivity of fluorescence. In total, we ran 1600 ns of MD.

## Data analysis and statistics

Data were analyzed using NIS Elements Advance Research software (versions 5.21.00 and 5.30.00), Excel (Microsoft 2021MSO), OriginPro (2019b, OriginLab), Fiji ImageJ (2.9.01/1.53t.), Clustal Omege(online version) and ESPrint 3.0^74^. Analysis of all fluorescence traces was conducted as follows: cells and a neighboring cell-free region were manually selected using NIS Elements Advance Research software, and fluorescence measurements were performed for each region of interest (ROI). Fluorescence from an RGEPO-free region was subtracted from the cell fluorescence to account for background, except for the data from acute brain slices. Fluorescence change (ΔF/F) was then calculated using the formula [(Fmax- F)/F], in which the F is the baseline fluorescence intensity. In addition, for acute brain slice data (Figure 6f, i), RGEPOs fluorescence traces were corrected for photobleaching by subtracting baseline fluorescence traces that were fit to an exponential function. Dose-response curves for different K^+^ were fitted with Hill equation (F=Fmax*[ligand]^n/(EC50^n+[ligand]^n)), while F is the fluorescence intensity for a defined concentration of ligand, Fmax is the maximum fluorescence intensity, EC50 is the half-maximum effective concentration, and n is the Hill coefficient. The data are represented as mean ± SD. Computational data of the systems were analyzed using cpptraj^75^(Root mean square deviation, hydrogen bonding percentages, distance measurements). For the visualization, and to calculate electrostatic charge distribution of the surface of systems we used ChimeraX^76^. To create the movie of potassium bound RGEPO systems we used VMD^77^.

For in vivo two-photon imaging of RGEPOs, we used NormCorre^78^ for piece-wise rigid motion correction of the images. Neuronal ROI segmentation was performed on the average projection image of GCaMP6f using CellPose^79^. Fluorescence traces were then extracted from the ROIs using ImageJ by averaging pixel intensities for each frame. The fluorescence response of cells was represented as a ΔF/F trace, where F is the average of the first 100 frames from the recording. A sliding window was then applied to smooth the ΔF/F trace. For temporal analysis of wave arrival in cell populations (Fig. 7g,h), we identified the frame number at which the ROI reaches peak fluorescence intensity. The ROIs were then colored using a gradient palette according to the frame number. For RGEPO1, analysis of peak fluorescence intensity did not reveal a clear pattern of wave propagation as observed in Fig. 7d. Therefore, we analyzed the time at which 37% of the peak ΔF/F was reached in the ROI for RGEPO1 as an approximation to demonstrate wave propagation in the neuronal population.

## Data availability

All essential raw datasets including source files for Supplementary Figures and raw unprocessed images, will be available at FigShare. All plasmids used in this study will be available from WeKwikGene (https://wekwikgene.wllsb.edu.cn/). Source data files are available from FigShare (https://doi.org/10.6084/m9.figshare.28067942.v1). Raw files for time-lapse imaging of Asp-RGEPO1 and GREPO2 expressed in the cultured neurons and astrocytes are available from FigShare (https://doi.org/10.6084/m9.figshare.27101941.v1). The computational inputs used to run simulations of systems, including topology files, initial coordinates of all the systems, last frames of the free diffusion simulations of both RGEPO1 and RGEPO2, the force field parameters created for chromophore, as well as a short simulation video of potassium diffusion in RGEPO1, have been deposited to Zenodo (https://doi.org/10.5281/zenodo.13824581).

## Acknowledgments

We are grateful to the Flow Cytometry Core Facility at Westlake University and the Laboratory Animal Resources Center. We also thank Zhong Chen from Westlake University for their assistance with spectroscopic measurements; M.H. Liao and G.C. Fang from Imaging Core at Westlake University for their technical support with the Nikon Confocal microscope. We thank the WSU Supercomputing Grid for computational support. We thank Minho Eom from the Korea Advanced Institute of Science and Technology for his advice on the analysis of *in vivo* two-photon images. The work was supported by start-up funding from the Foundation of Westlake University, Westlake Laboratory of Life Sciences and Biomedicine, National Natural Science Foundation of China grant 32171093, and “Pioneer” and “Leading Goose” R&D Program of Zhejiang 2024SSYS0031 to K.D.P. The authors thankfully acknowledge support from the National Science Foundation−NSF through the grant NSF CHE2338804 to A.R.W. We thank Wayne State University and Department of Chemistry for the Thomas C. Rumble University Graduate Fellowship to V. P.

## Author contributions

L.Y. and K.D.P. initiated the project and interpreted the data. L.Y., F.V.S., and K.D.P. performed the initial design of the sensor prototype. L.Y., with help from S.Z., performed the direct evolution of sensors in *E. coli* and HEK293FT cells. L.Y., with help from C.L., characterized RGEPOs in vitro. L.Y, with help from H.Z., performed the characterization of RGEPOs in cultured neurons and astrocytes. L.Y. and X.S performed the acute brain slice and imaging. V.P. and A.R.W. performed MD simulations. S.Z. performed the two-photon image in awake mice.

C.G. set up the perfusion system. L.Y., S.Z., and K.D.P. analyzed the data and, together with V.P. and A.R.W., wrote the manuscript. K.D.P. made high-level designs and plans and oversaw all aspects of the project.

---

**Supplementary Figure 1.**
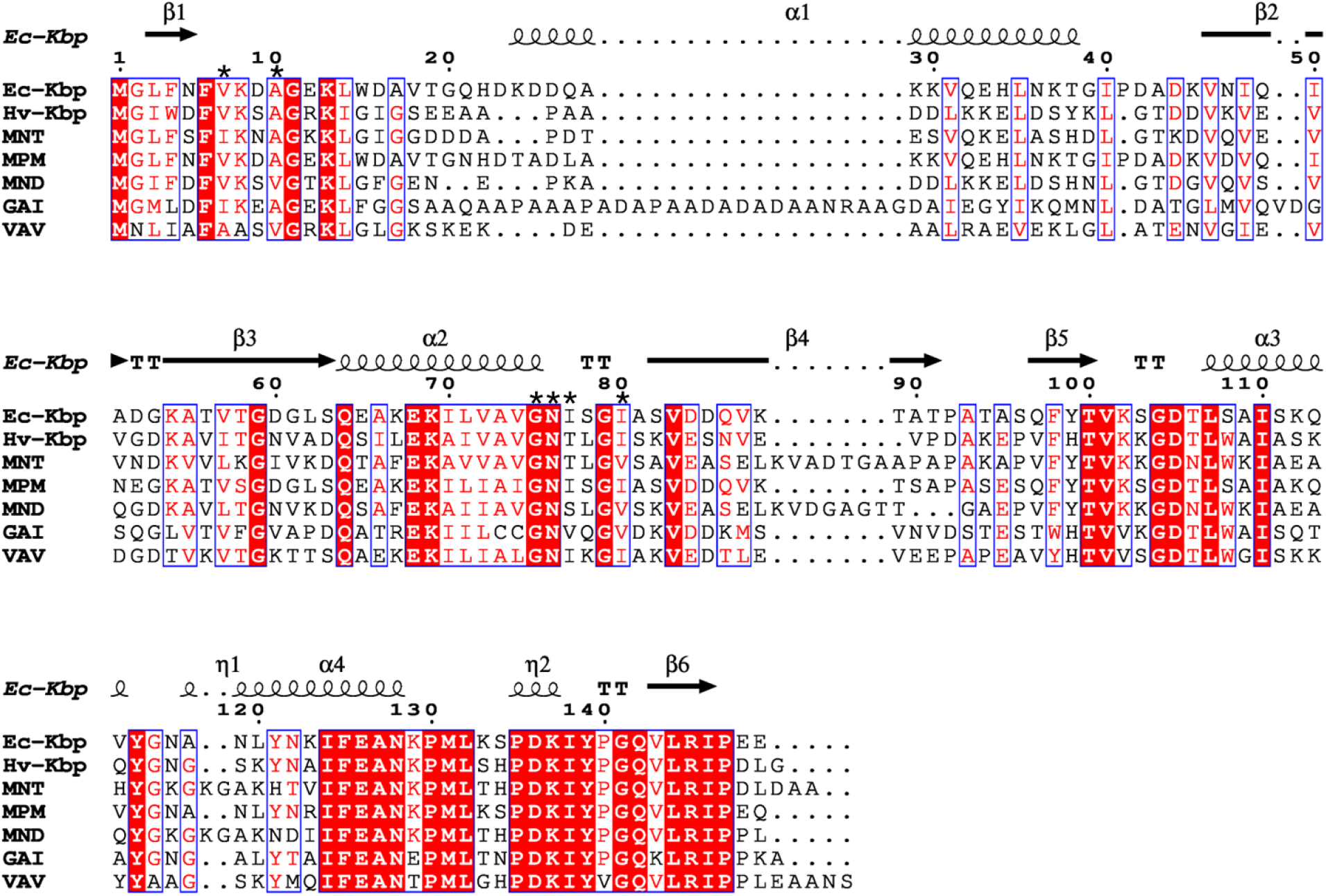
Alignment of amino acid sequences of Ec-Kbp and Hv-Kbp with five homologs obtained from a metagenomic BLAST search. Residues within the red boxes with white text indicate the strict identity of amino acids, while those within a blue frame denote similarity among the homologs. Labels for β-sheet-forming regions and α-helix-forming regions are provided as arrows/straight lines and curved lines, respectively. The symbol ’η’ denotes a 310 helix, and ’TT’ indicates strict β-turns. Residues located within 3.0 Å of the potassium ion in the Ec-Kbp structure (PDB: 7PVC) are marked with asterisks (N = 6 amino acids). The amino acid numbering follows that of Ec-Kbp. Alignment of protein sequence was conducted via Clustal Omega and subsequently shown with Ec-Kbp structure by ESPript 3.0.

**Supplementary Figure 2.**
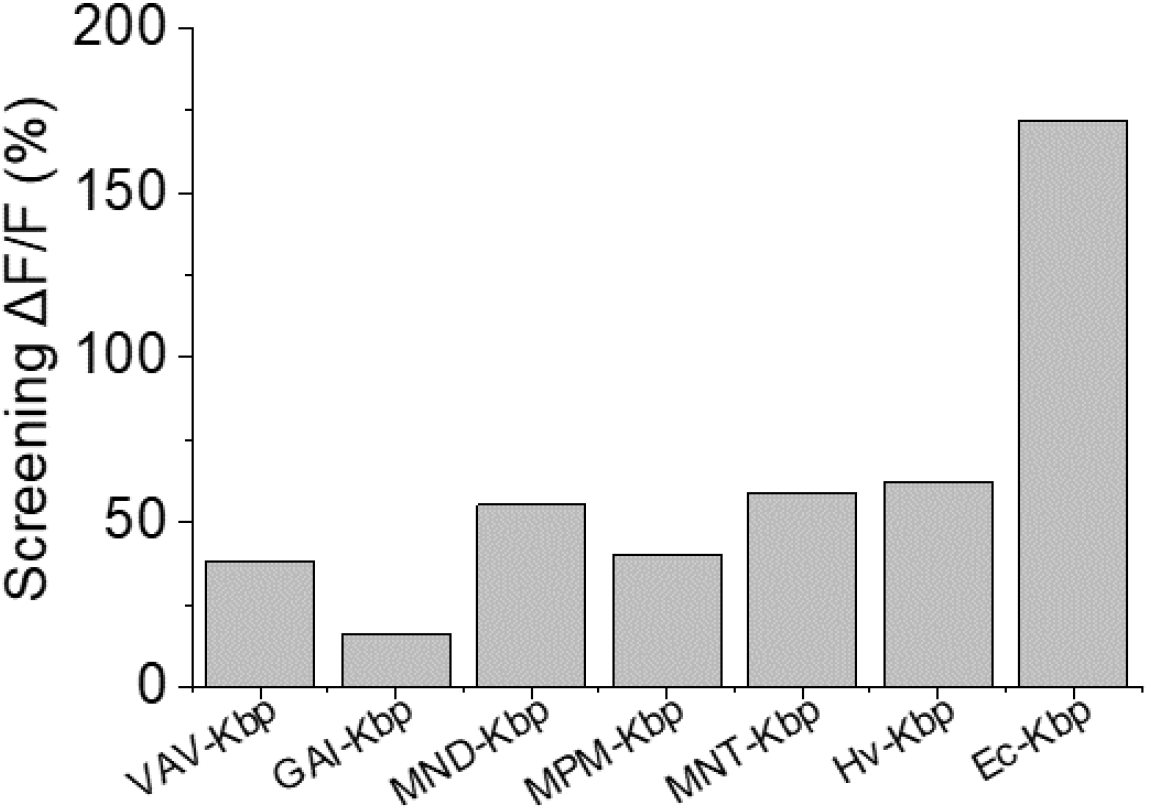
Fluorescence response of potassium green fluorescence sensors generated by swapping Ec-Kbp with Hv-Kbp and five newly identified homologs in solution. The homologs were inserted into mNeoGreen following the previous KRaION1’s design^4^ and titrated with 230 mM K^+^ in solution.

**Supplementary Figure 3.**
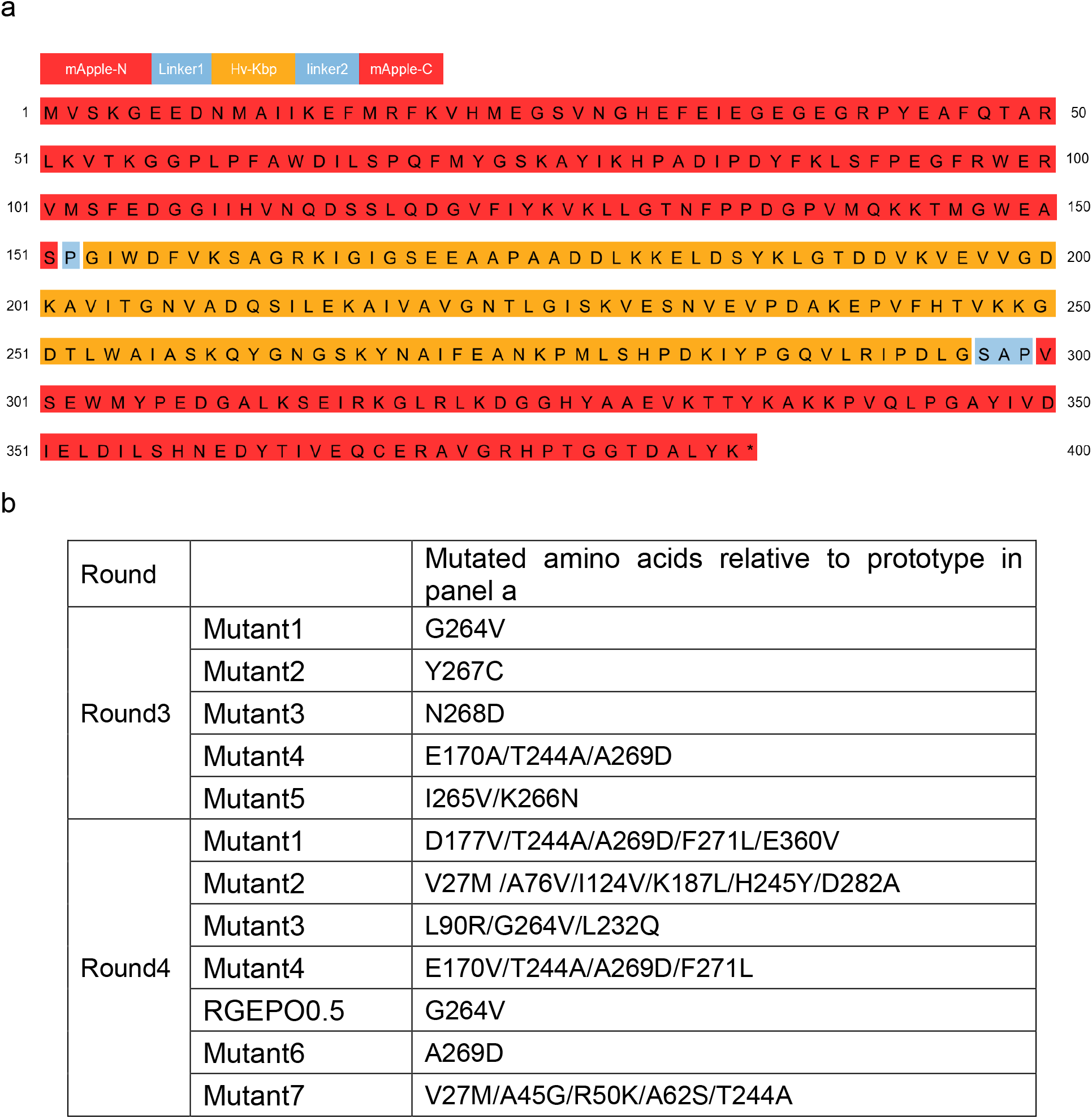
Amino acids sequence of prototype and relative mutants. (a) Amino acids sequence of RGEPO-prototype sensor with linker sequences Linker1: P and Linker2: SAP. (b) The mutants were validated in mammalian cells, corresponding to the red- highlighted dots in Figure 1e.

**Supplementary Figure 4.**
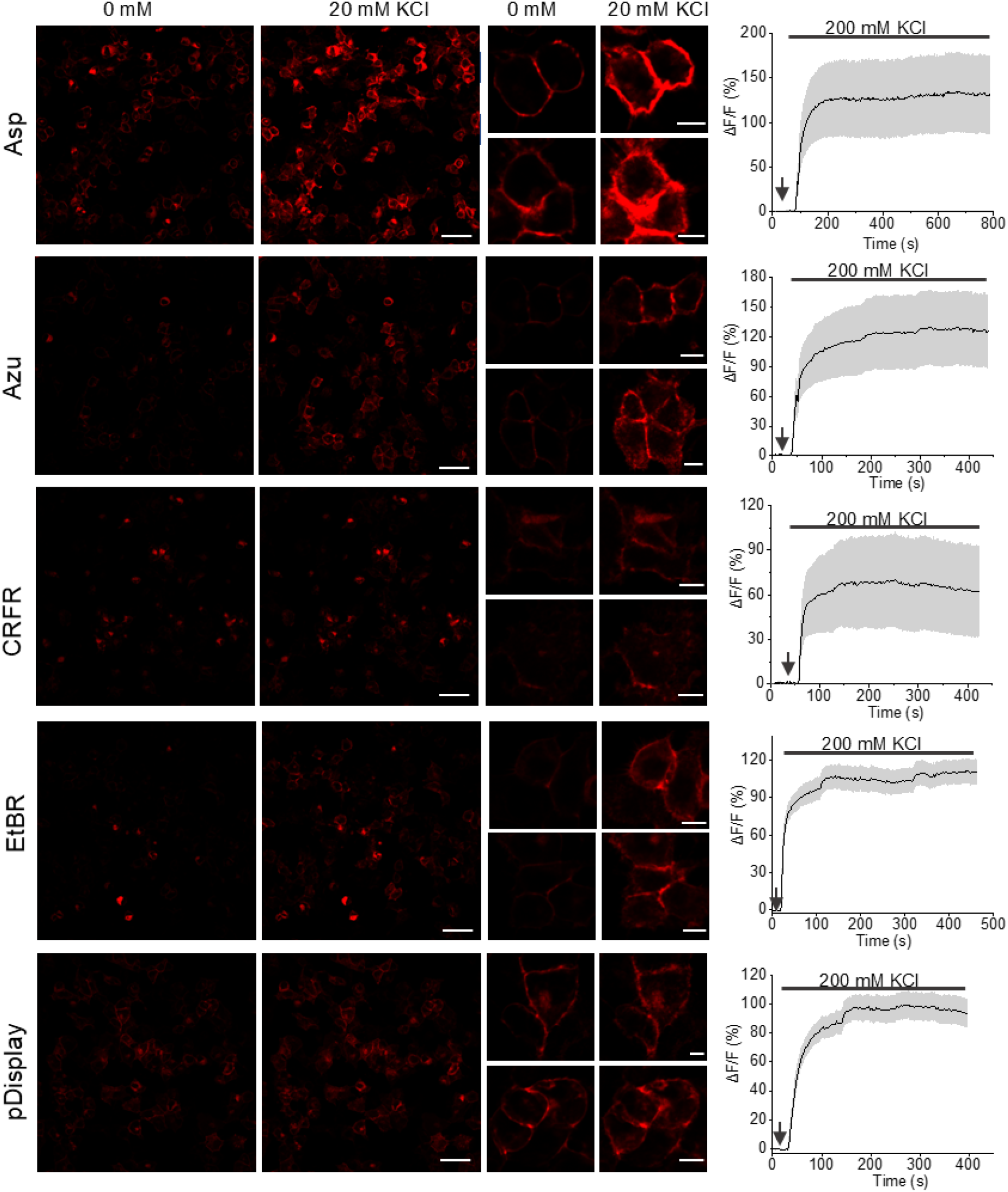
Representative single-plane confocal fluorescence images of leader- sequence variants and single-trial optical traces. Left, single plane fluorescence images of RGEPOs with 5 different leader sequences expressed in HEK293FT cells in response 20 mM KCl (n=2 field of views (FOVs) from 1 culture). Scale bars, 50 µm. Middle, magnified images in K^+^ free and K^+^ binding state (n= 3 cells from 2 FOVs over 1 culture for each). Scale bars, 10 µm. Right, single-trial optical traces of RGEPOs with different leader sequence upon stimulation of 200 mM KCl, which is shown in the Fig. 2e (n= 31, 13, 22, 22, 24 cells from 1 culture for Asp, Azu, CRFR, EtbR and Igκ, respectively).

**Supplementary Figure 5.**
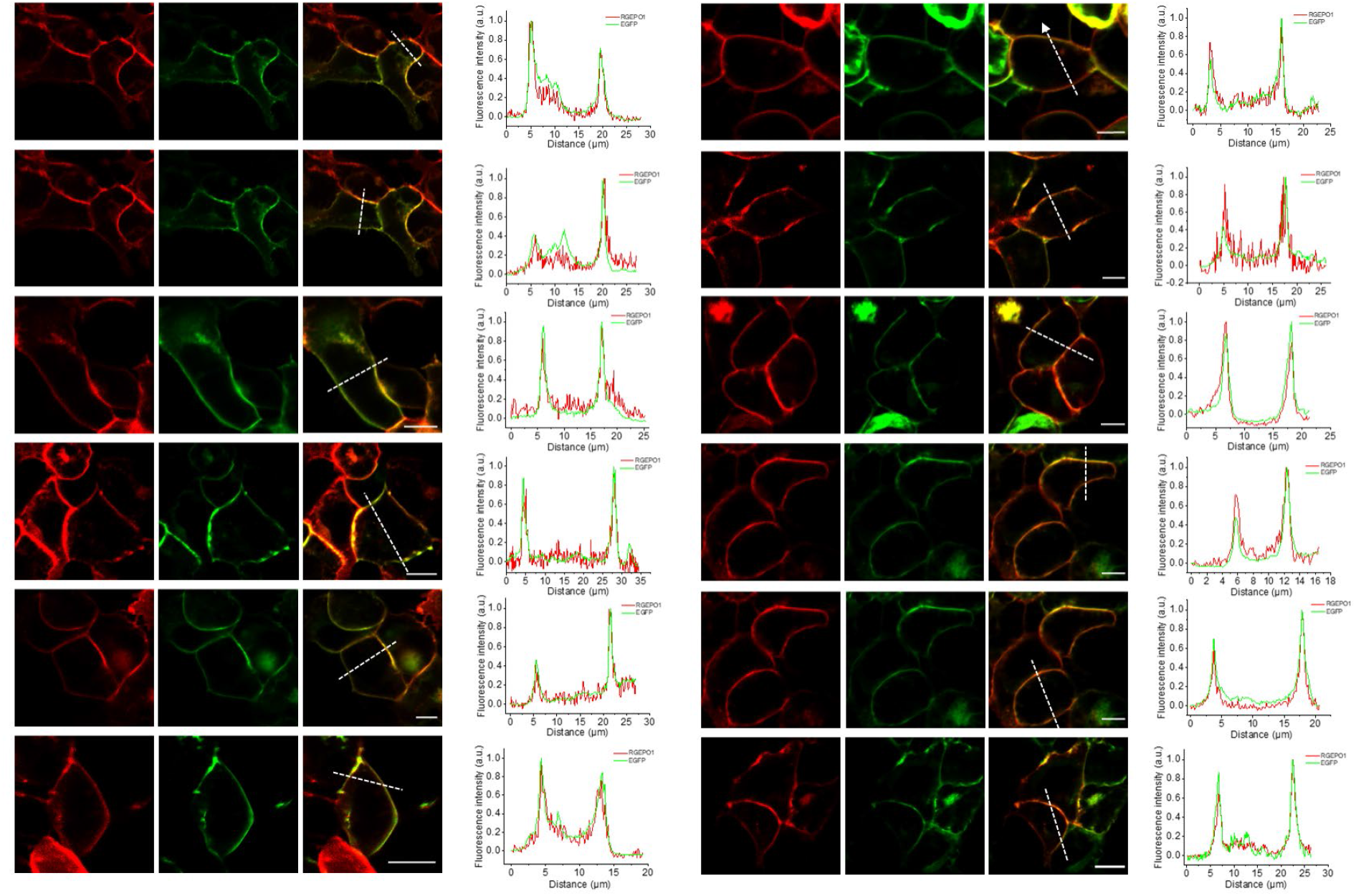
Validation of membrane localization of Asp-RGEPO0.5 expressed in HEK293T cells. Fluorescence images and membrane localization analysis of Asp-RGEPO0.5 expressed in HEK293FT cells. Membrane-targeted GFP (Igk-GFP) was co-expressed to label the plasma membrane. Left, single-plane confocal fluorescence images of HEK293T cells expressing the RGEPO0.5 (red) and EGFP (green) (n= 12 cells from 2 independent transfections from one culture). Right, normalized linecut (shown as white dashed line on the left) plots of fluorescence signals measured in both the red and green channels. Scale bars, 10µm

**Supplementary Figure 6.**
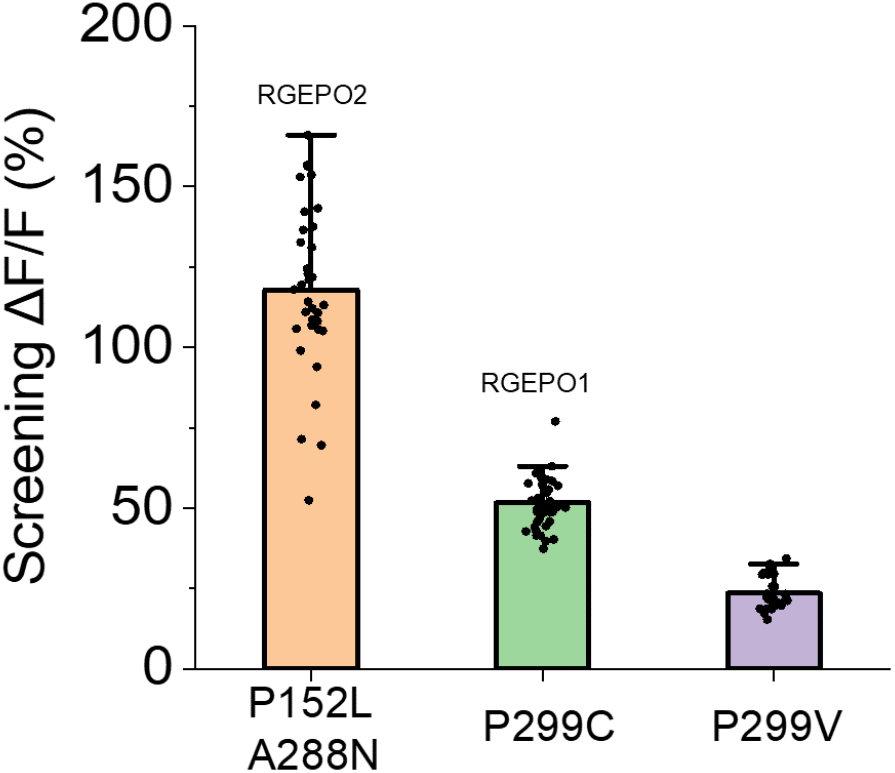
Validation of the top three mutants in the cytoplasm of HEK293FT cells. Validation of the top three mutants, which exhibited the highest extracellular responses, in the intracellular environment using a perfusion system (n= 34, 44, 33 cells from 1 culture each, respectively). Dot, individual data point for single cells; bar, mean; error bar, SD.

**Supplementary Figure 7.**
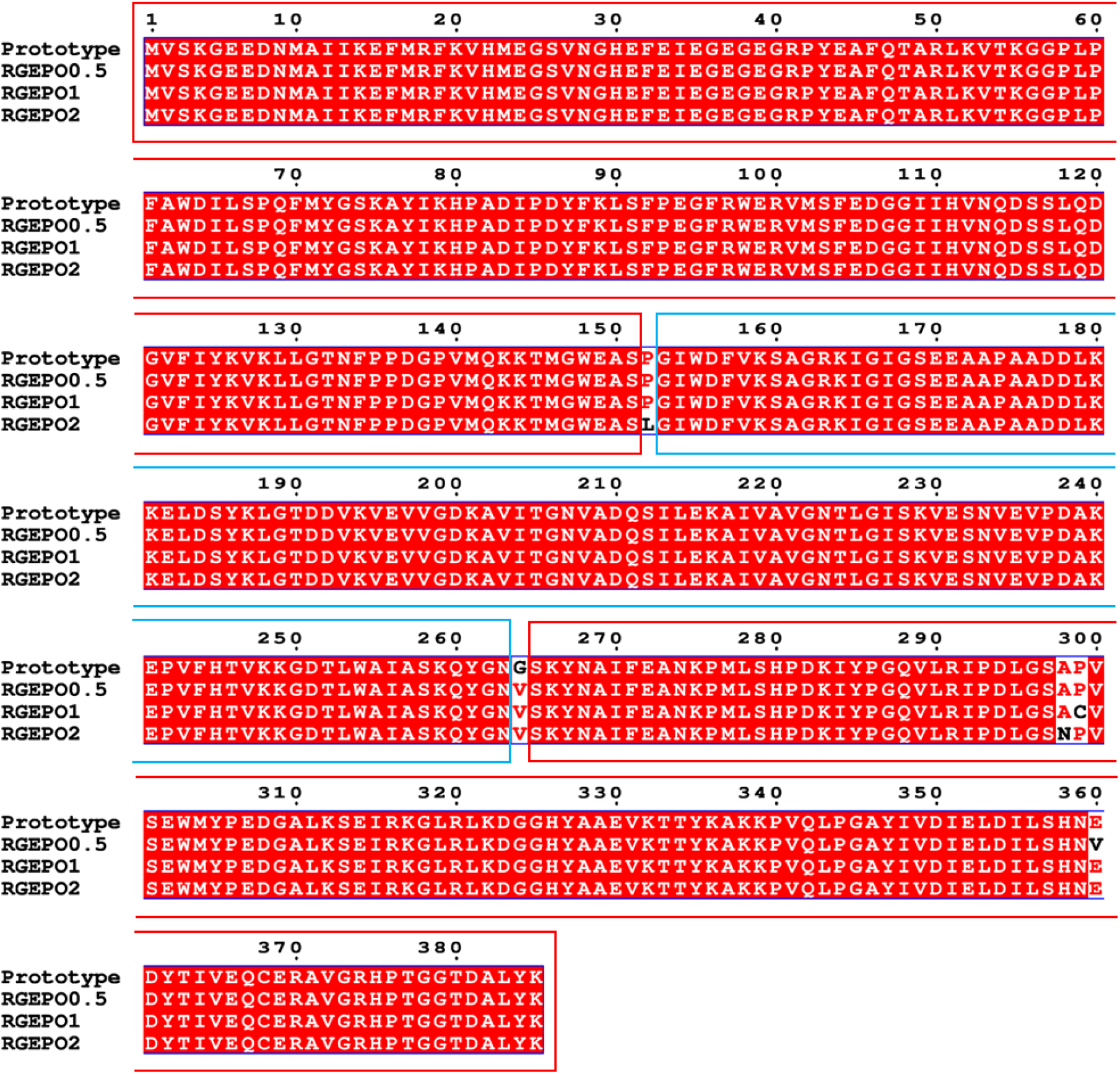
Amino acids sequence of prototype, RGEPO0.5, RGEPO1 and RGEPO2. The sequences derived from mApple and the potassium-binding domain are highlighted in red and blue boxes, respectively. The non-red shaded areas indicate mutations acquired during the engineering process.

**Supplementary Figure 8.**
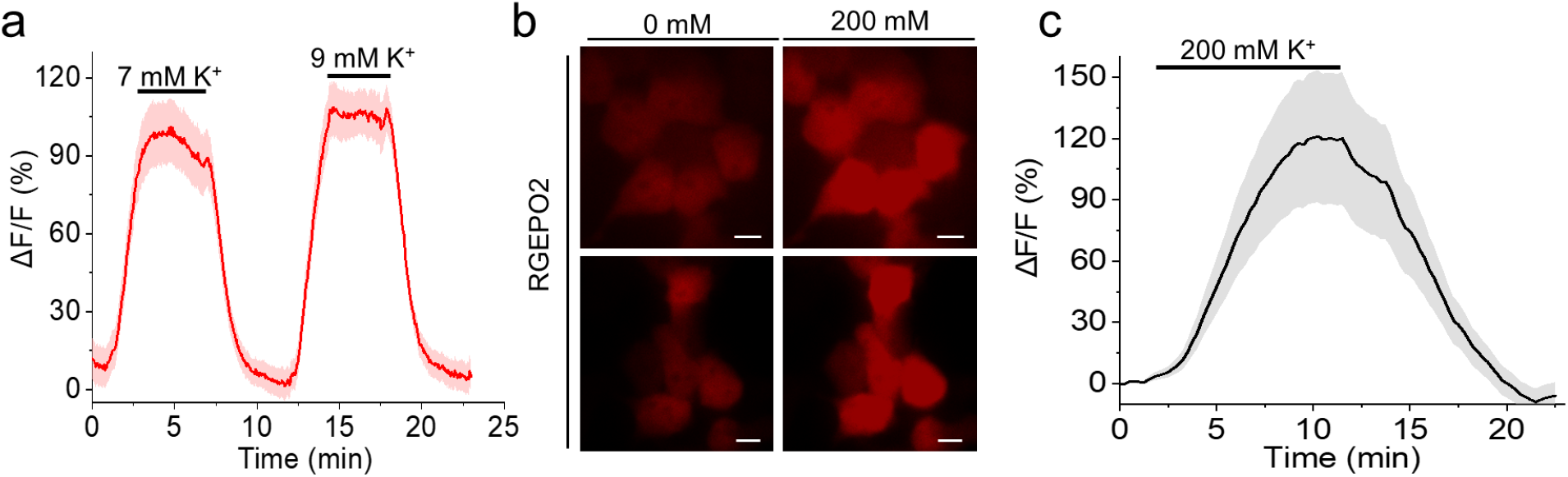
Validation of RGEPOs’ reversibility and representative images of RGEPO2 expressed in the cytoplasm of HEK cells. (a) Fluorescence intensity change (ΔF/F0) time course of RGEPO1 with stimulation by a series of K^+^ buffer on HEK293T cells (n= 5 cells from 1 culture), data are expressed as mean (solid line) and SD (shaded area). (b) Representative images of RGEPO2 expressed in the HEK293FT cells (n= 32 cells from 1 cultures). Scale bar, 10 µm. Images were obtained via an inverted wide-field Nikon Eclipse Ti2 microscope with 20x NA0.75 objective lens. (c) Fluorescence intensity change (ΔF/F) time course of RGEPO1 with stimulation by a series of K^+^ buffer on HEK293T cells using valinomycin and CCCP (n= 34 cells from 1 culture), data are expressed as mean (solid line) and SD (shaded area).

**Supplementary Figure 9.**
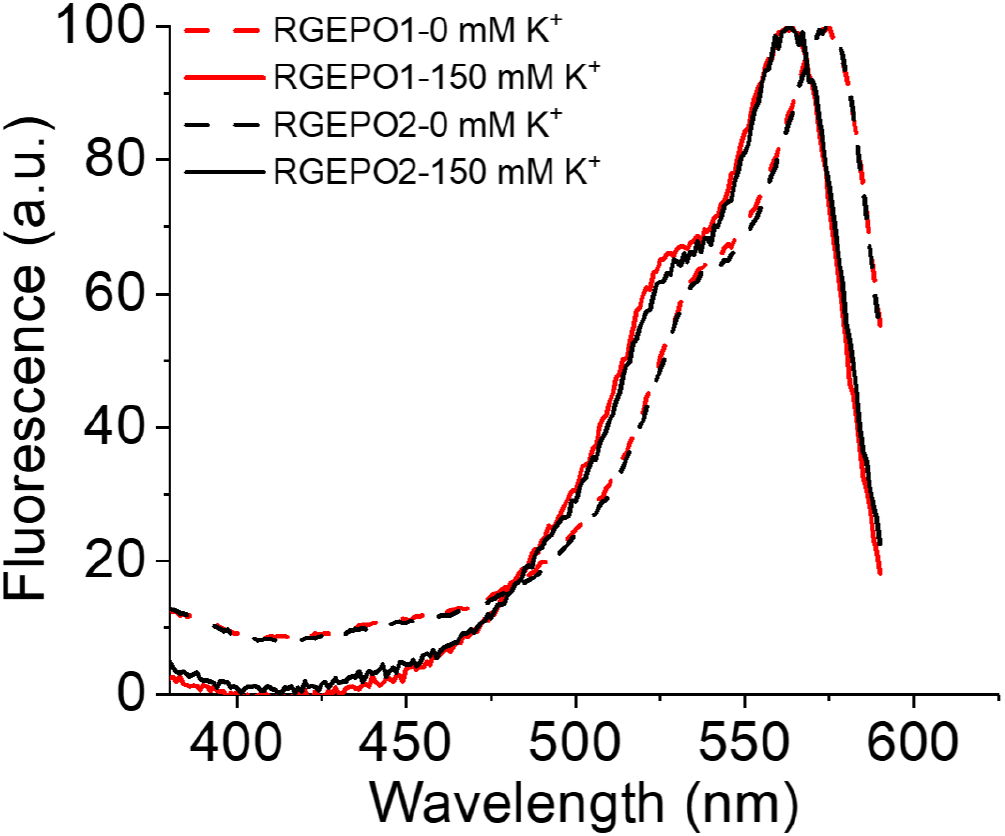
Steady-state excitation fluorescence spectra of RGEPOs at 0 and 150 mM potassium at pH = 7.4.

**Supplementary Figure 10.**
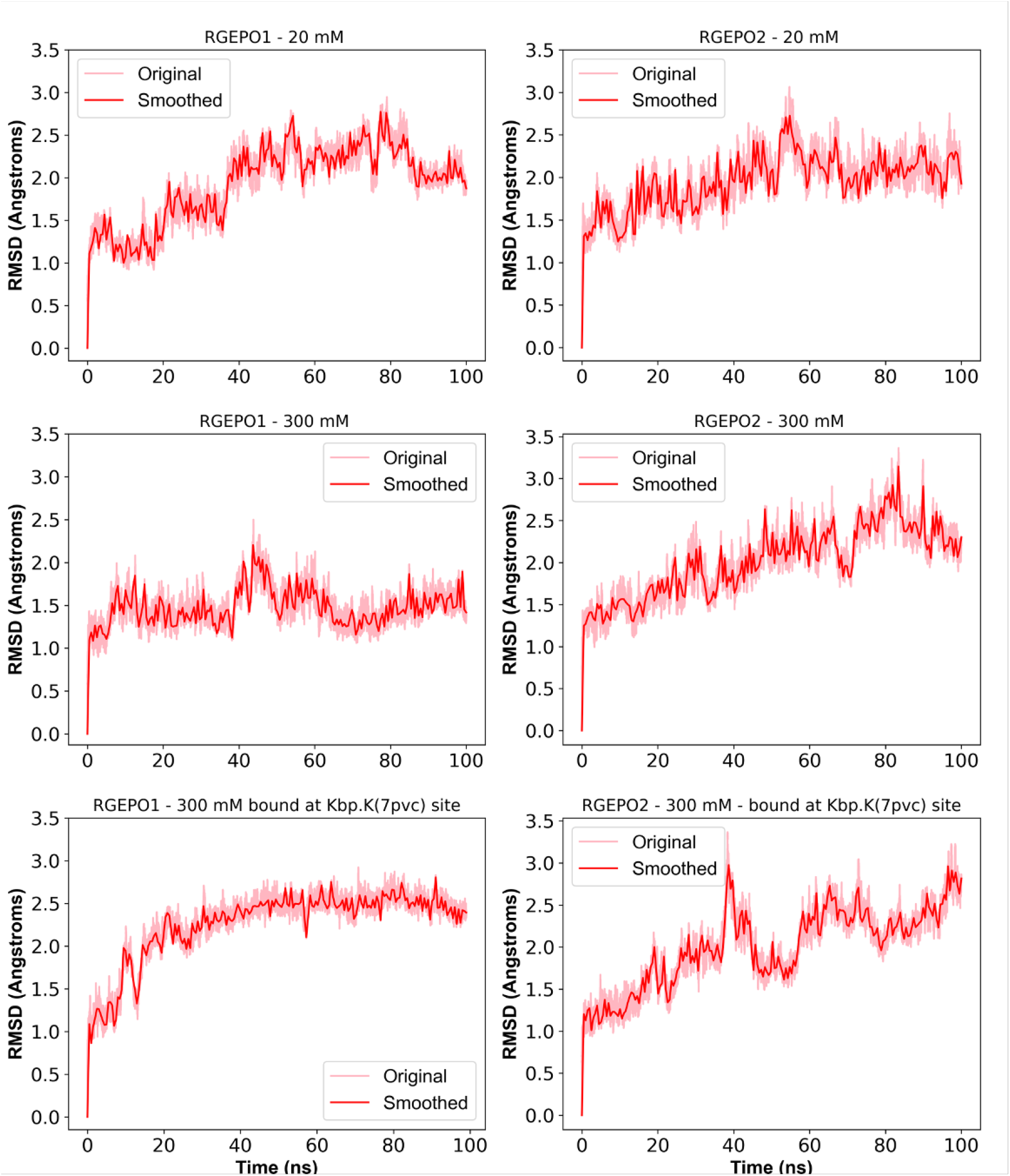
Change in root mean square deviation (RMSD) of RGEPO1 and RGEPO2 systems with time over first 100 ns. The first two row graphs show the free diffusion simulations of RGEPO1 (left column) and RGEPO2 (right column) at their respective ionic concentrations. The last row graphs represent simulations of RGEPO1 (left) and RGEPO2 (right) with a 300 mM ionic concentration, initiated with a K^+^ bound at the originally reported Kbp.K binding site (PDB ID: 7PVC).

**Supplementary Figure 11.**
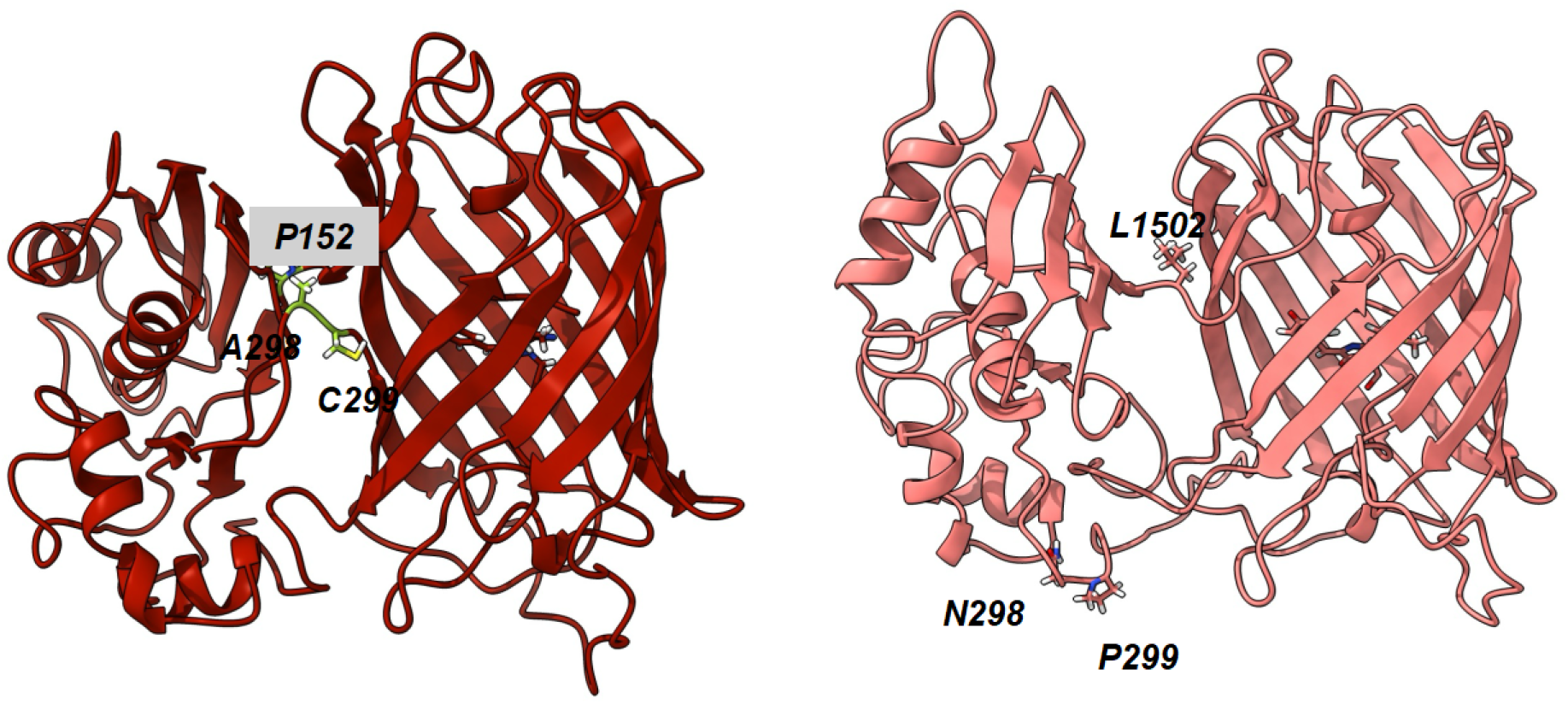
Comparison of mutation positions with time. Structure of RGEPO1 (in red) and RGEPO2 (in salmon pink) with mutated positions highlighted in sticks.

**Supplementary Figure 12.**
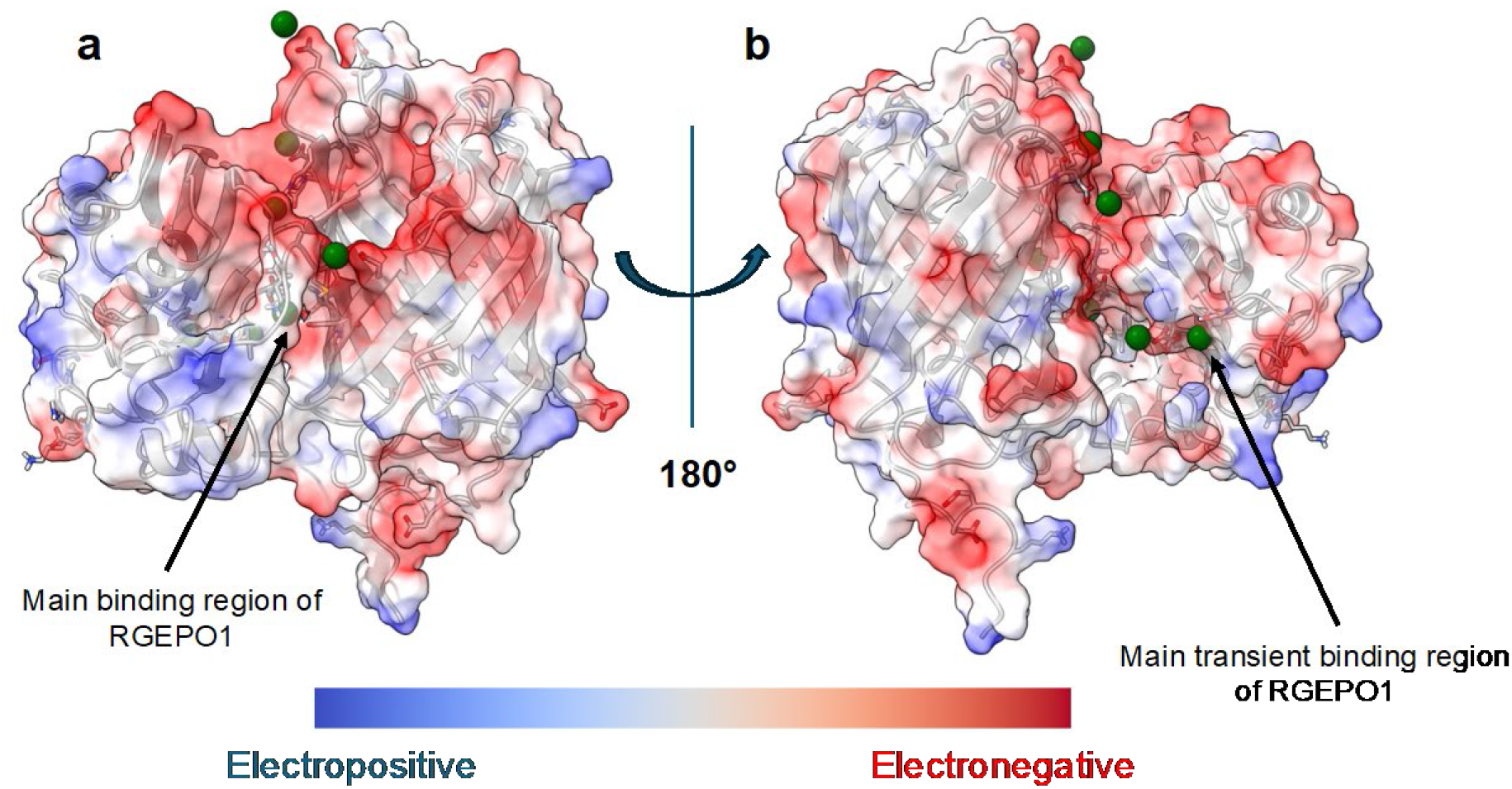
Possible transient sites of RGEPO1. Figures a and b show the electrostatic charge distribution on the surface of RGEPO1 (a: front view, b: back view after a 180-degree rotation). In these figures, red represents more negatively charged regions, while blue represents more positively charged regions, including both potential transient sites and main binding sites (labeled with an arrow) as observed in free diffusion simulations with K^+^ shown in green spheres.

**Supplementary Figure 13.**
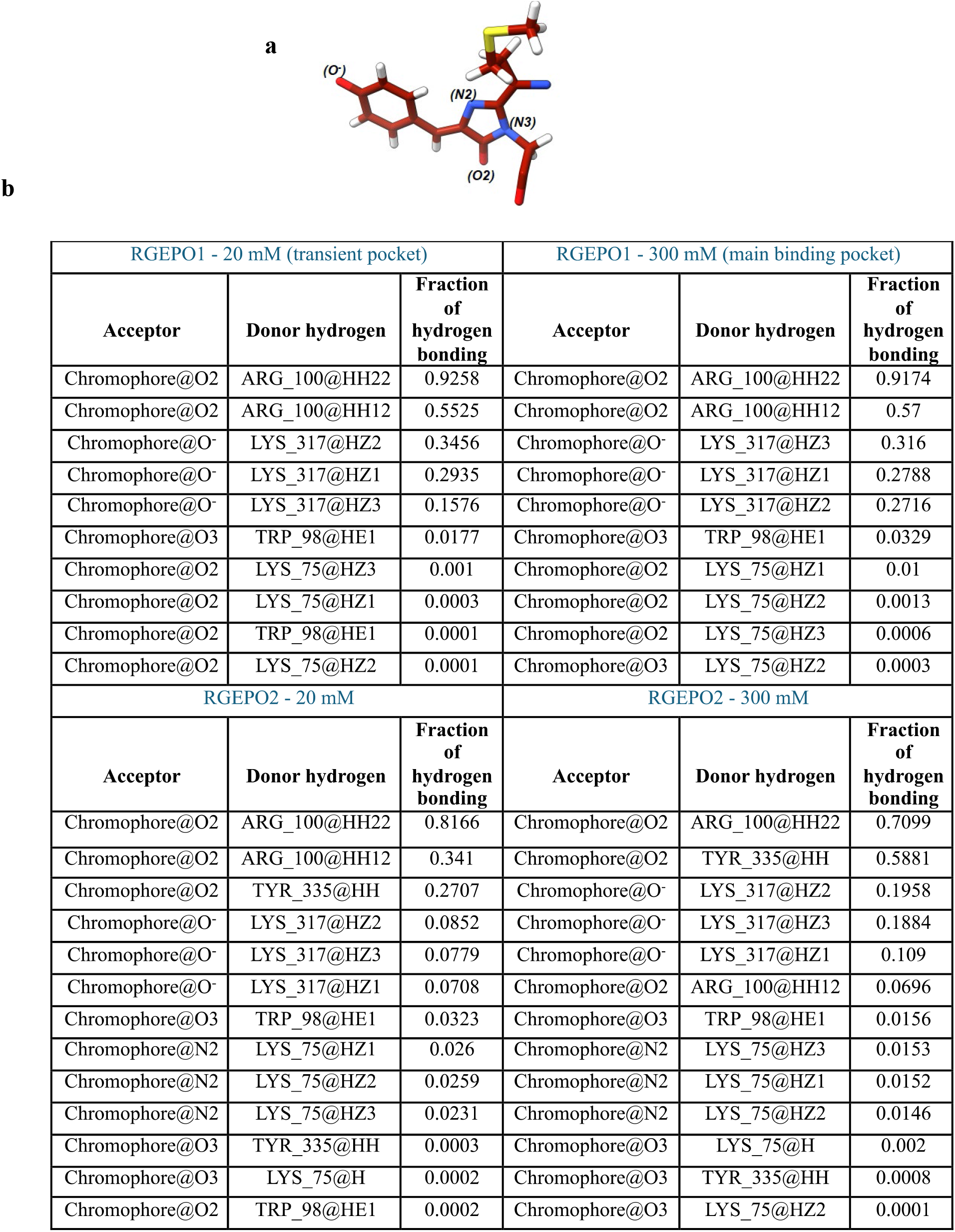
(a) Atom names of the chromophore and (b) table of hydrogen bonding analysis.

**Supplementary Figure 14.**
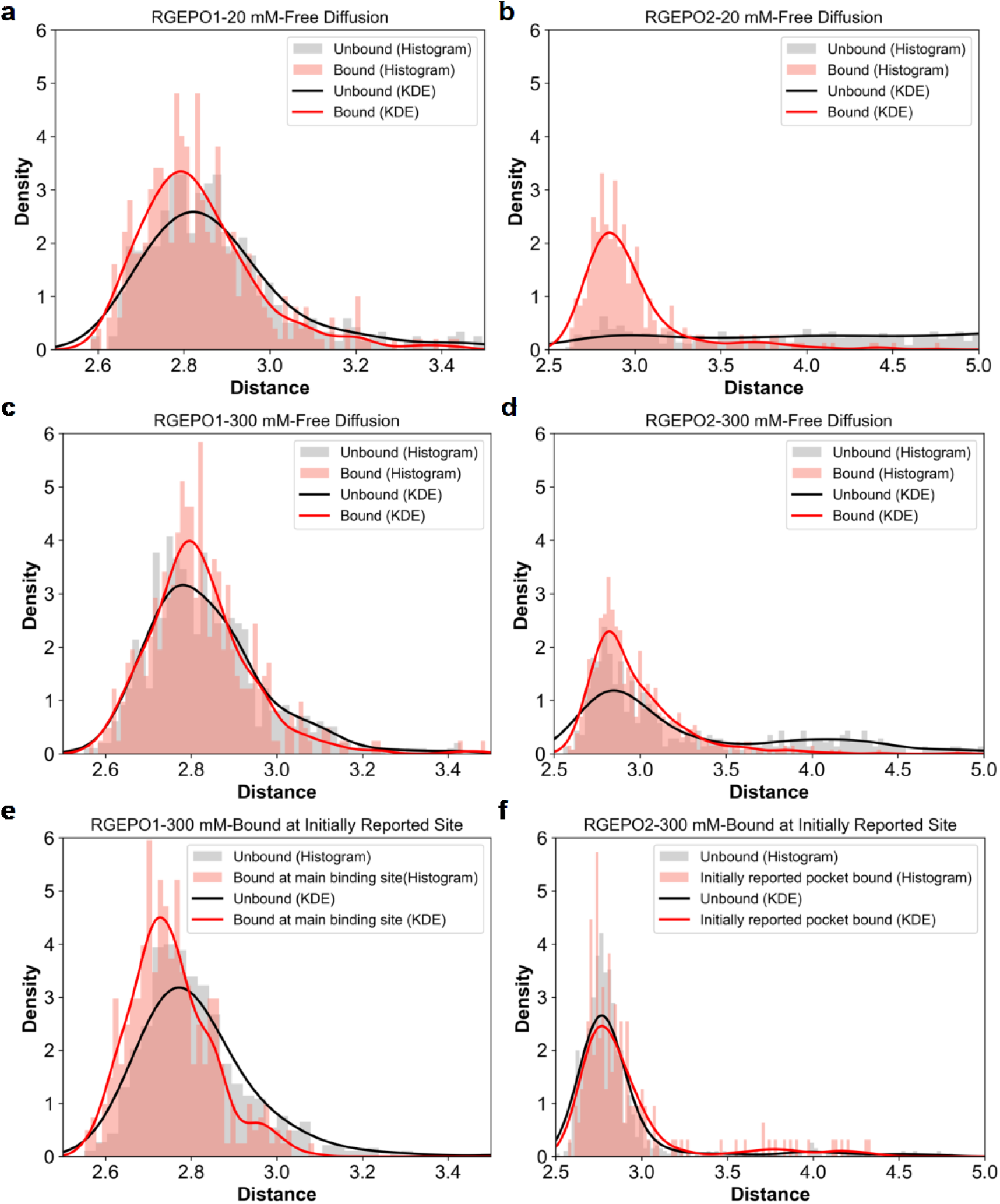
Variation of distance between K317 side chain nitrogen atom and the O^-^ atom of the chromophore during K^+^ bound and unbound states. Graph (a) shows the density plot for the variation in distance between the K317 side chain nitrogen and the O^-^ of the chromophore, where free diffusion of K^+^ occurs at the transient binding site of RGEPO1 at 20 mM concentration. Graphs (b) to (d) display the density plots for the free diffusion simulations of K^+^ at the main binding sites of RGEPO1 and RGEPO2, as labelled. Graphs (e) and (f) correspond to K^+^ bound simulations at the initially reported Kbp.K site. In graph (e), the analysis focuses on the free diffusion of K^+^ at the main binding site, while in graph (f), it pertains to K^+^ at the initially reported site.

**Supplementary Figure 15.**
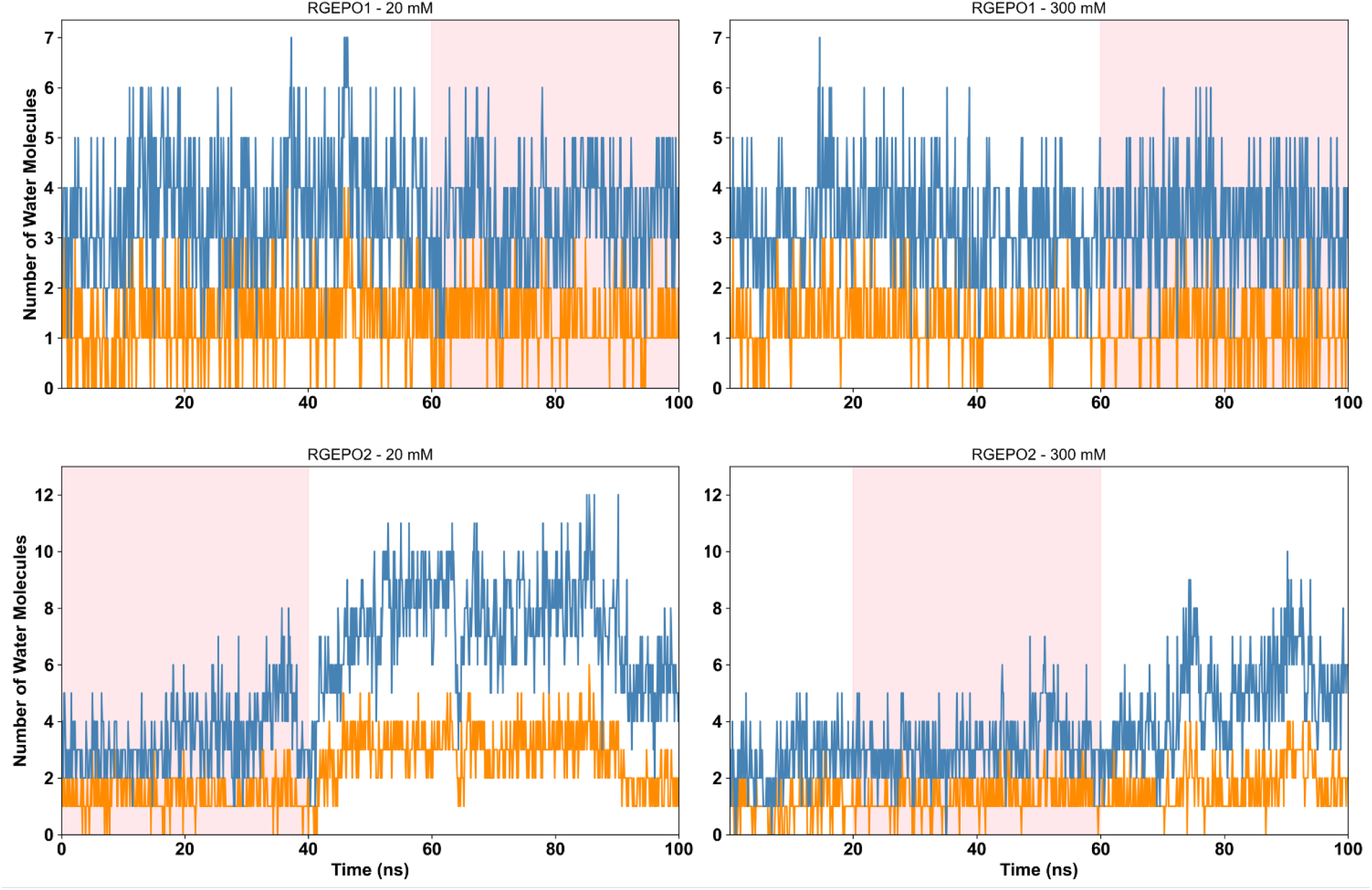
Variation of number of water molecules at the O^-^ end of the chromophore. These graphs depict the change in the number of water molecules around the O^-^ position within 3.5 Å (orange) and 5 Å (blue) throughout the trajectory. The time periods during which a K^+^ ion is bound are highlighted in pink. All simulations are for free diffusion of K+, with binding occurring at the main binding site. However, the simulation for RGEPO1 at 20 mM concentration specifically focuses on the transient binding of K^+^.

**Supplementary Figure 16.**
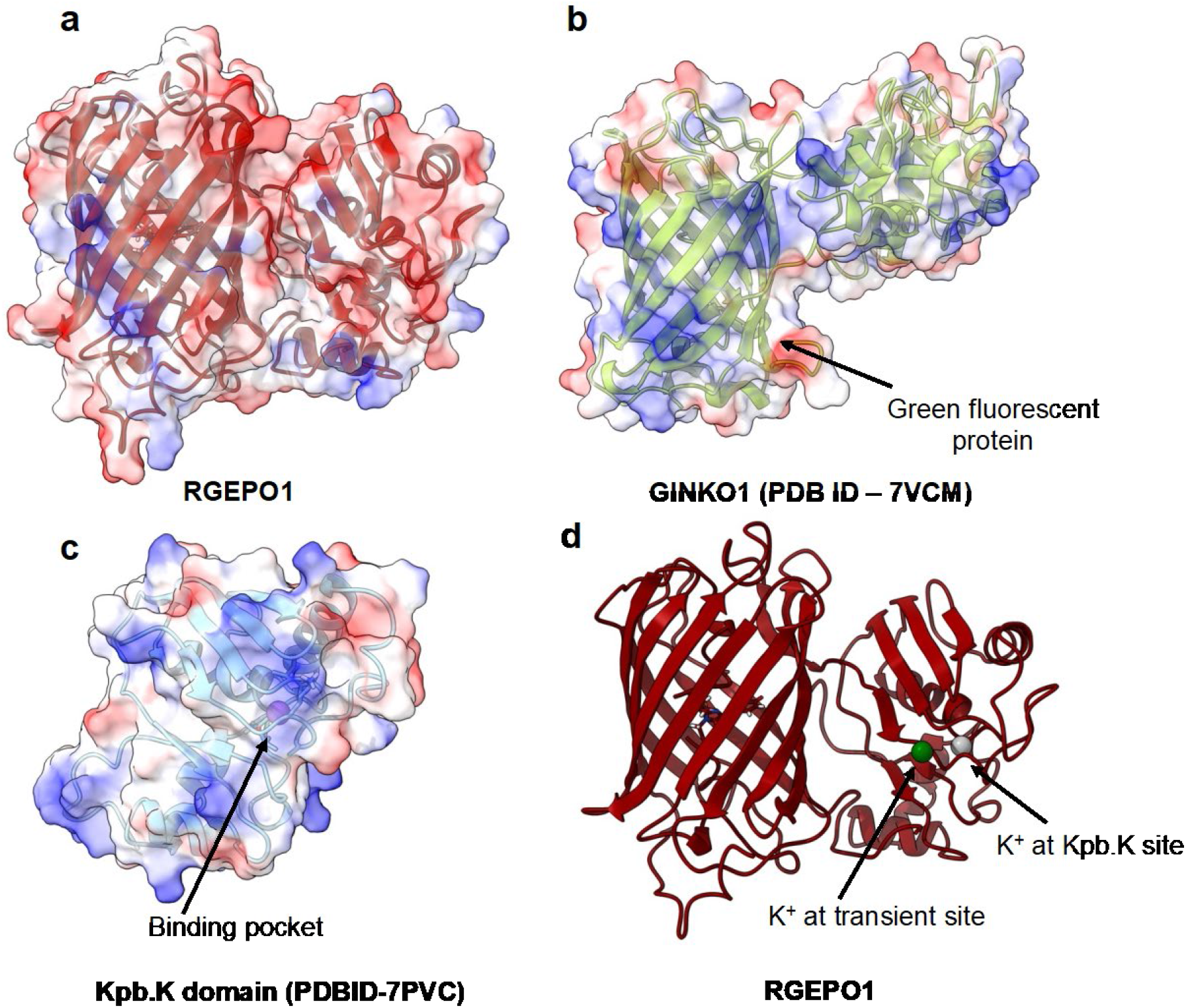
The electrostatic charge distribution of potassium indicators. (a) The electrostatic charge distribution of RGEPO1 is shown, with red indicating more electronegative regions and blue representing more electropositive regions. (b) The electrostatic charge distribution of GINKO1 (genetically-encoded potassium ion biosensor) (PDB ID – 7VCM) is shown, with red indicating more electronegative regions and blue representing more electropositive regions. (c) The electrostatic charge distribution of the Kbp.K domain (PDB ID: 7PVC) is shown, with red indicating more electronegative regions and blue representing more electropositive regions. The bound K^+^ is depicted as a purple sphere. (d) The structure of RGEPO1 is shown in red, with the transiently bound K^+^ in a green sphere and the K^+^ at the Kbp.K site (PDB ID: 7PVC) in a grey sphere.

**Supplementary Figure 17.**
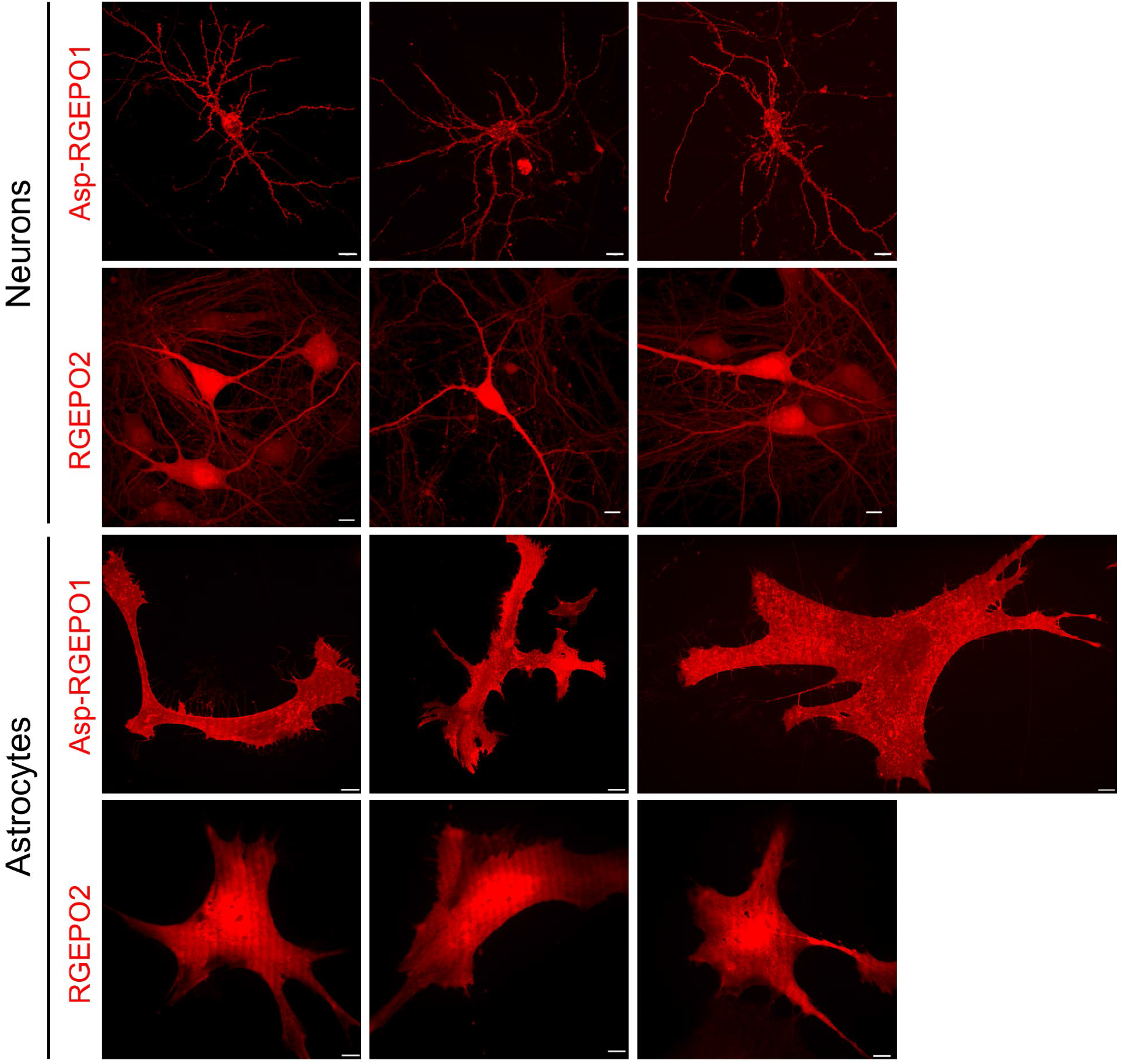
Representative images of live mouse hippocampal neurons expressing Asp-RGEPO1 and RGEPO2. Maximum intensity projection confocal images of Asp-RGEPO1 on the extracellular surface and RGEPO2 in the cytoplasm of live cultured neurons and astrocytes in ACSF buffer (n= 4 and 17 neurons from 2 independent cultures; n=25 and 17 astrocytes from 2 independent cultures). Scale bars, 10 µm (scale bar of the second image of Asp-RGEPO1on the astrocyte is 30 µm).

**Supplementary Figure 18.**
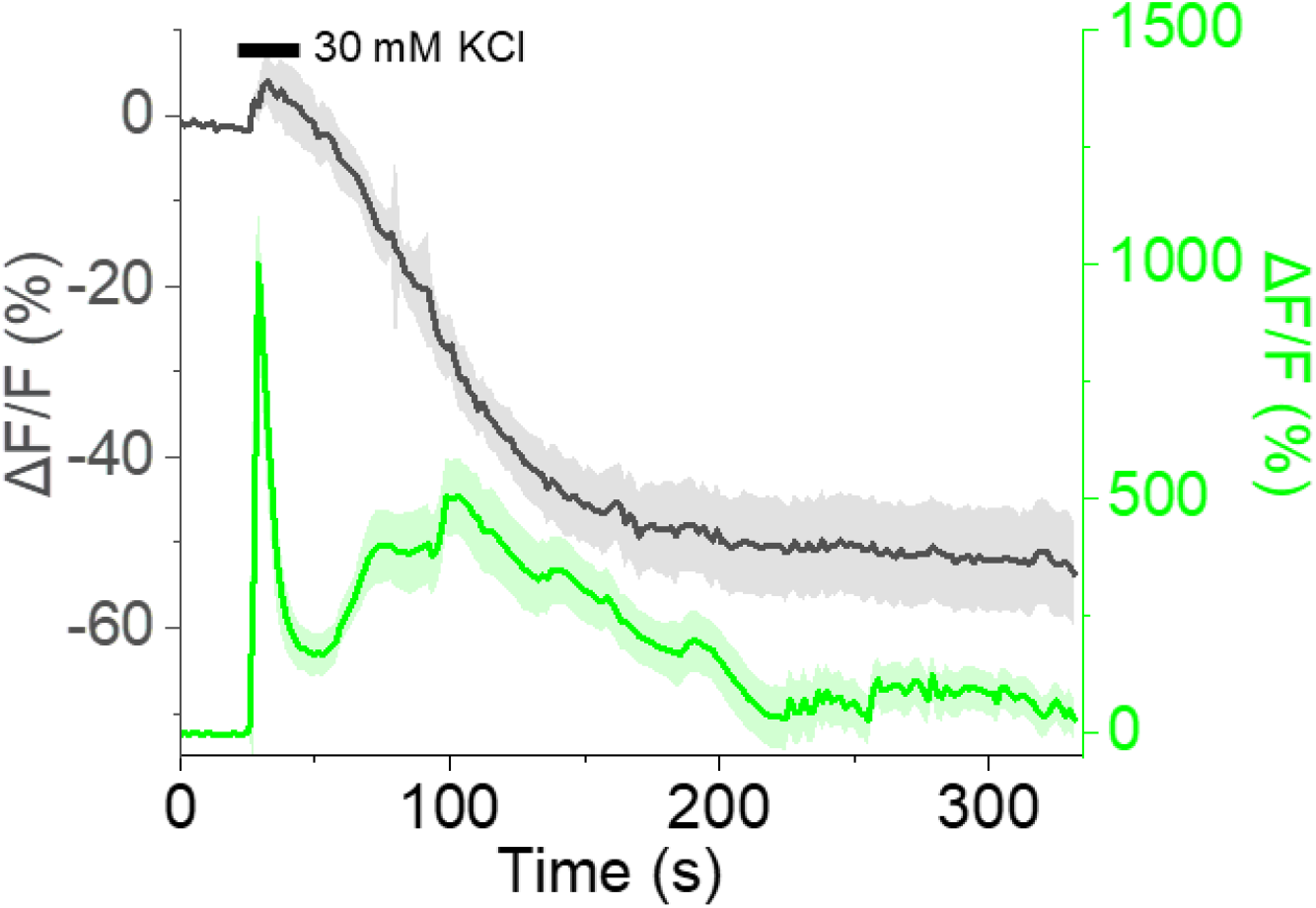
Time course of fluorescence intensity changes of Asp-RGEPO1 and GCaMP6f. Time course of fluorescence intensity changes for Asp-RGEPO1 and GCaMP6f in response to 30 mM KCl stimulation in neurons using a perfusion system. n= 38 neurons from 1 culture.

**Supplementary Figure 19.**
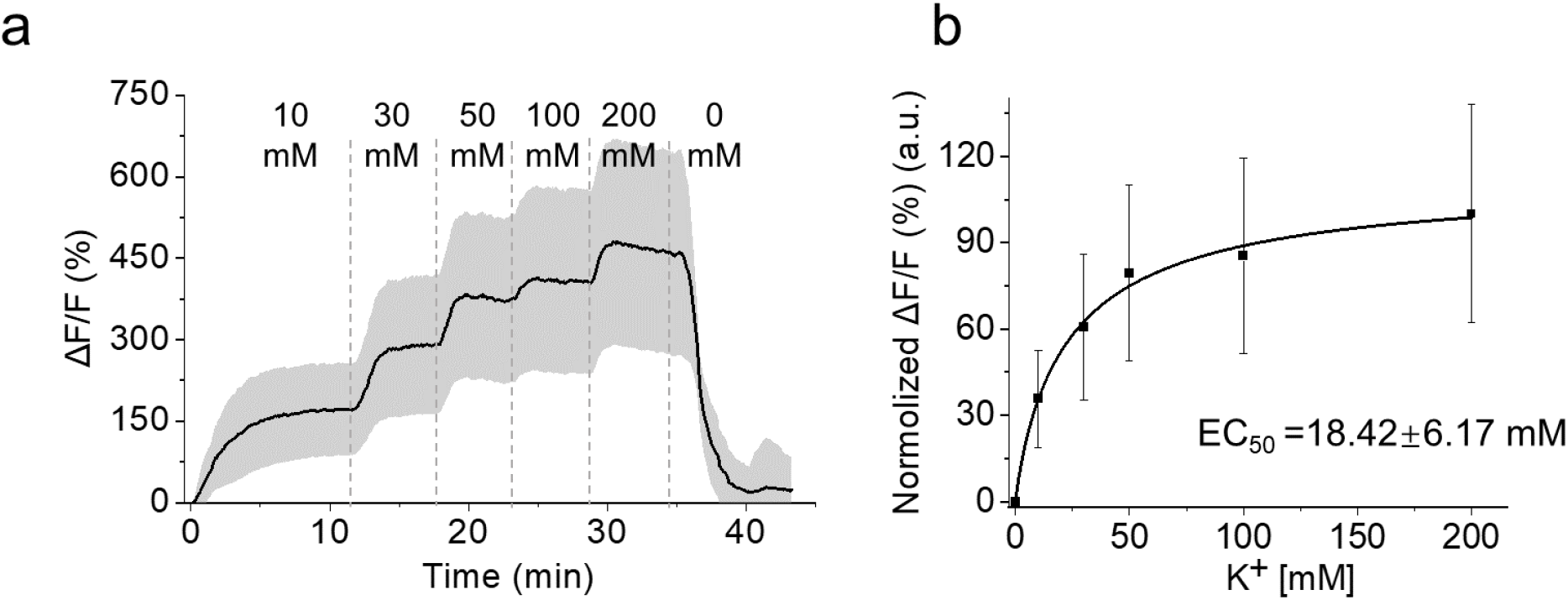
Time course of fluorescence intensity change of RGEPO2 titrated with a series of KCl buffer. (a) Time course of fluorescence intensity change of RGEPO2 with stimulation by a series of KCl buffer using valinomycin and CCCP (n= 12 cells from one culture), data are expressed as mean ± SD. (b) Plot of normalized ΔF/F against different K^+^ concentrations fitted using nonlinear fitting (Hill) for the data shown in panel a.

**Supplementary Figure 20.**
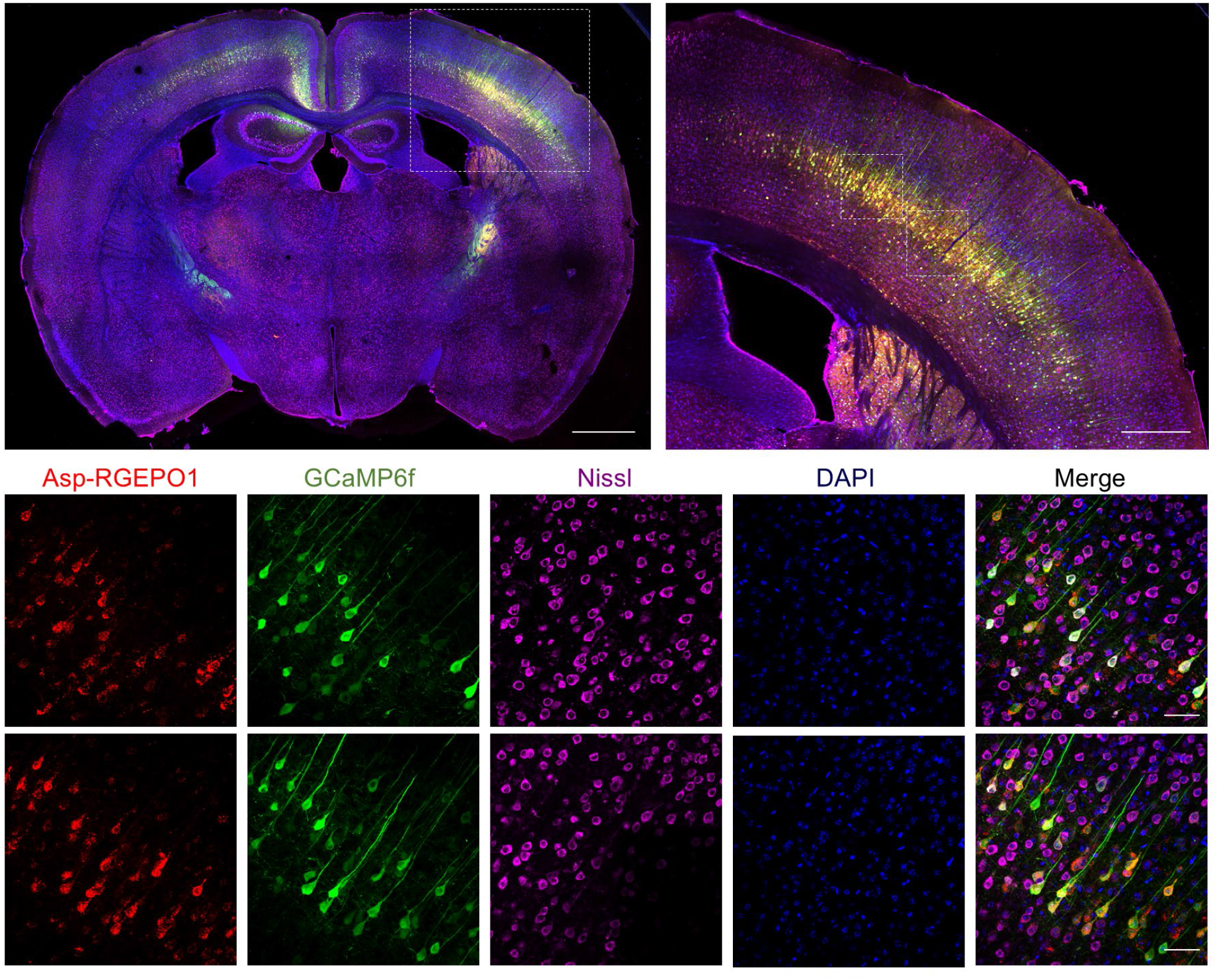
Image of the cerebral cortex showing the expression of Asp-RGEPO1. Upper left, single panel confocal fluorescence image of the cerebral cortex displaying the expression of Asp-RGEPO1 (red) (scale bar = 1000 µm). Upper right, higher magnification of single panel confocal fluorescence images highlighting relative expression regions (scale bar = 500 µm). Lower, further magnified multi-panel confocal fluorescence images showing detailed expression regions (scale bar = 50 µm).

**Supplementary Figure 21.**
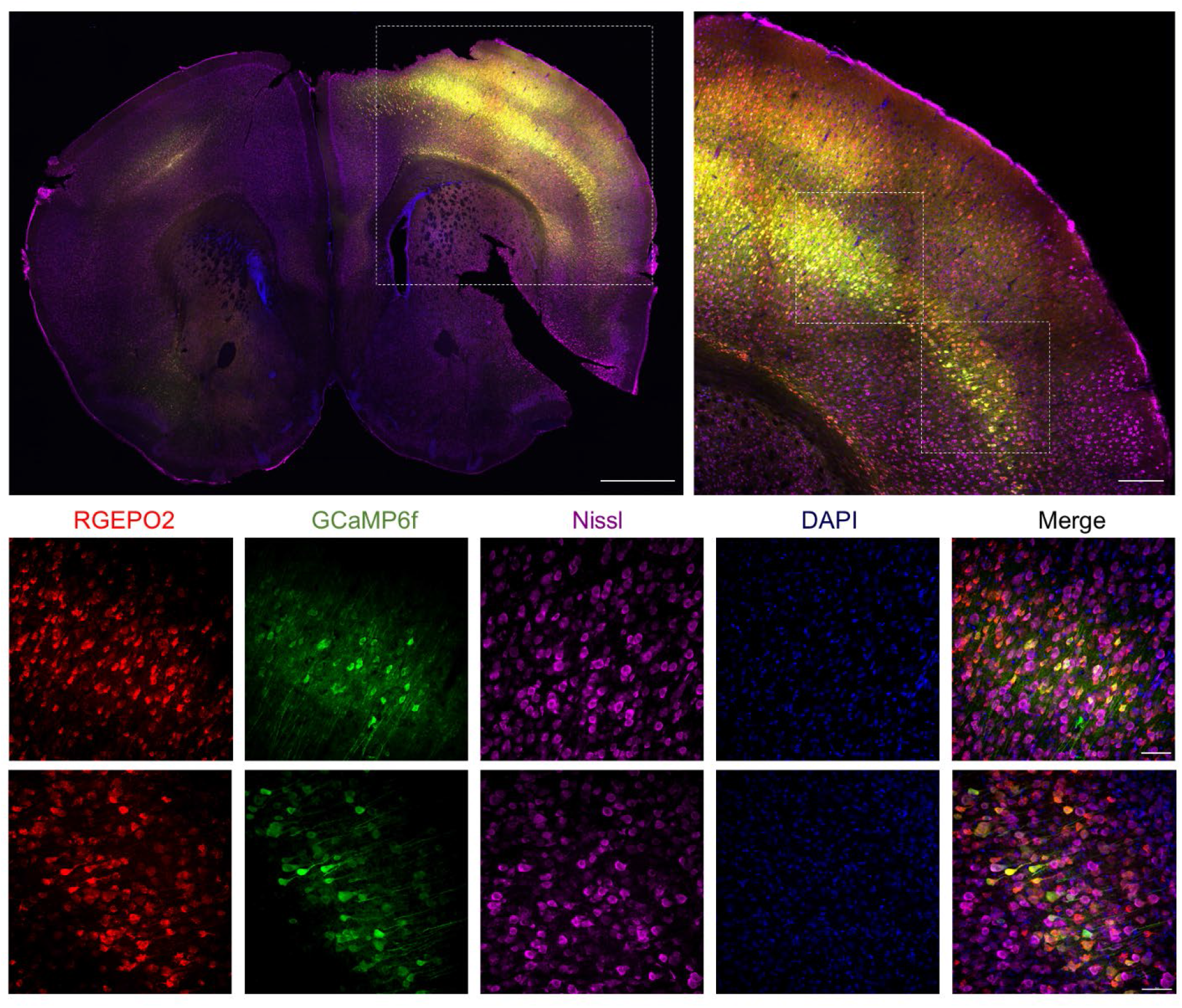
Image of the cerebral cortex showing the expression of RGEPO2 and GCaMP6f. Upper left, single panel confocal fluorescence image of the cerebral cortex displaying the expression of RGEPO1 (red) and GCAMP6f (green) (scale bar = 1000 µm). Upper right, higher magnification of single panel confocal fluorescence images highlighting relative expression regions (scale bar = 200 µm). Lower, further magnified multi-panel confocal fluorescence images showing detailed expression regions (scale bar = 50 µm).

**Supplementary Figure 22.**
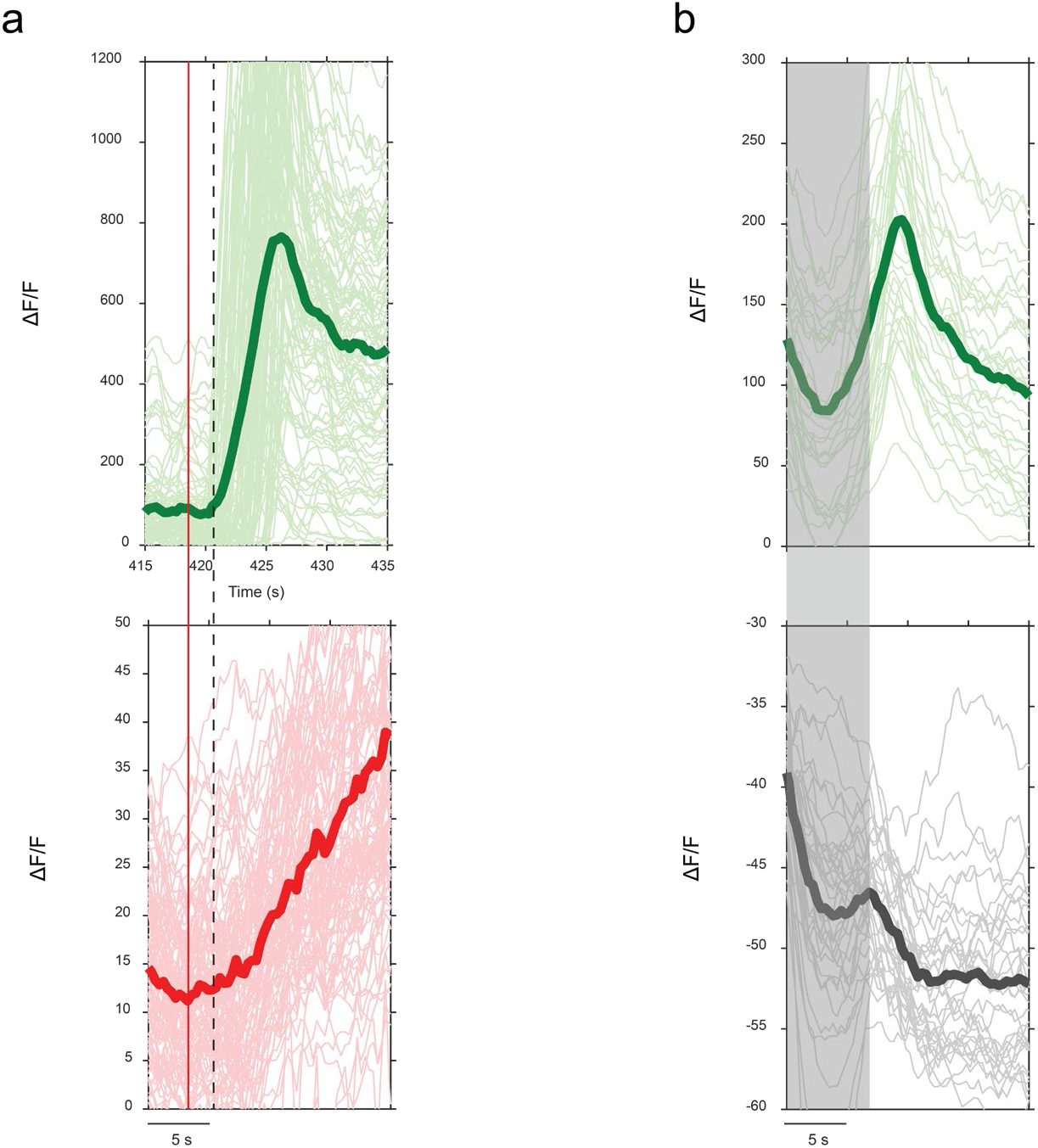
Estimated spreading wave onset for GCaMP6f and RGEPOs in mice with KA-induced seizures. a. Zoomed-in view of individual (light green and light red) and averaged (green and red) fluorescence responses of GCaMP6f and RGEPO1 from Fig. 7c. Statistics are the same as in Fig .7c. Red line indicates the rise of RGEPO1 fluorescence. Dashed line indicates the rise of GCaMP6f fluorescence. b. Zoomed-in view of individual (light green and light gray) and averaged (green and dark gray) fluorescence responses of GCaMP6f and RGEPO2 from Fig. 7e. The gray shaded box indicates the period of acute motion observed along the z-axis. Statistics are the same as in Fig .7e.

**Supplementary Table 1.**
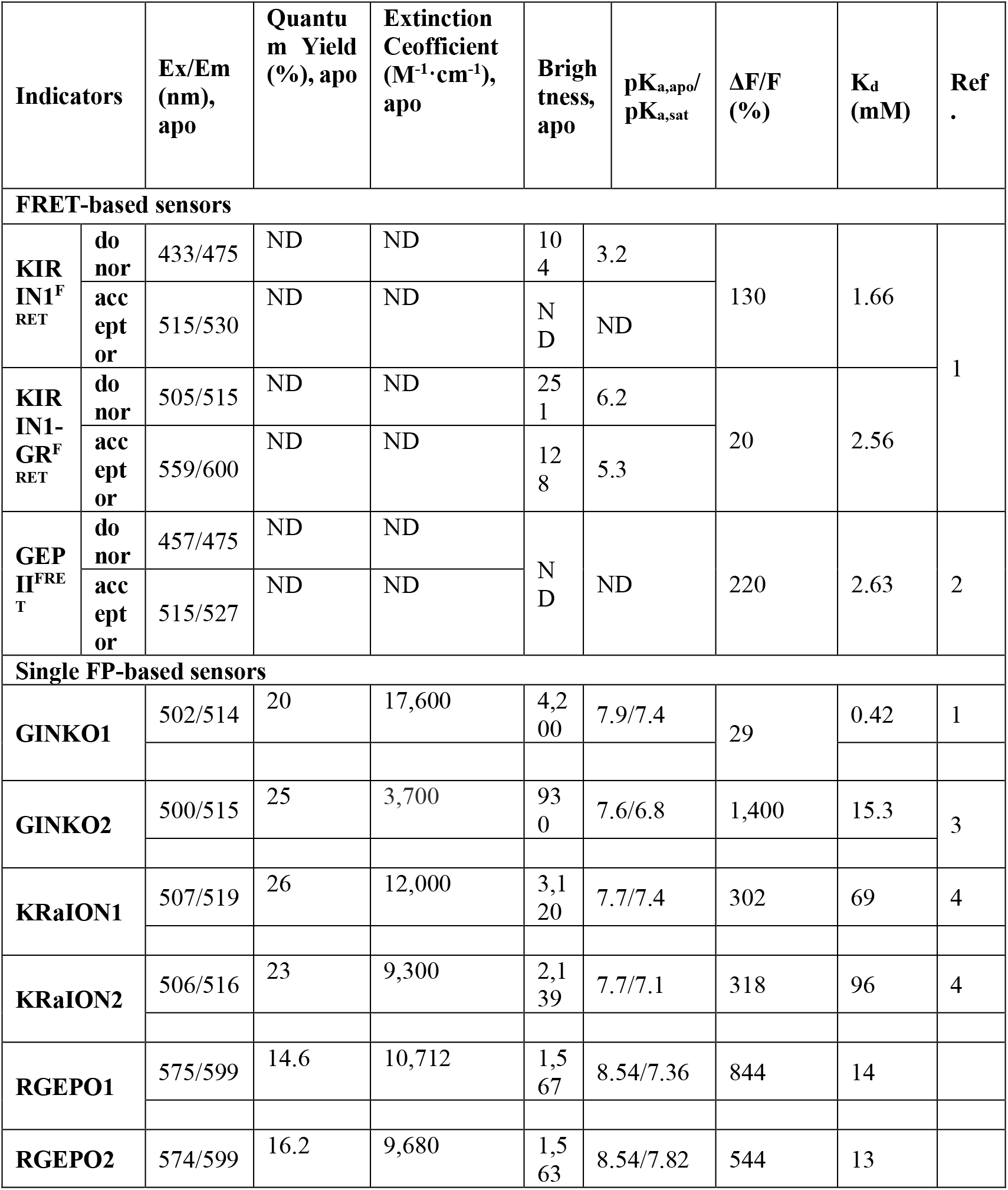
Properties of genetically encoded fluorescent potassium biosensors.

## Description of supplementary movies

**Supplementary Movie 1:** Time-lapse imaging of Asp-RGEPO1 on the extracellular surface of HEK293FT cells in response to 20 mM KCl. Imaging was conducted at 20°C-25°C using confocal microscopy.

**Supplementary Movie 2:** Time-lapse imaging of RGEPO2 expressed in the cytoplasm of HEK239FT cells in response to 200 mM KCl in the presence of valinomycin and CCCP. Imaging was conducted at20°C-25°C using an inverted wide-field Nikon Eclipse Ti2 microscope.

**Supplementary Movie 3:** Time-lapse imaging of RGEPO2 in the cytoplasm of cultured neurons in response to 500 µM glutamate. Imaging was conducted at 20°C-25°C using an inverted wide- field Nikon Eclipse Ti2 microscope.

**Supplementary Movie 4:** Time-lapse imaging of dual-color fluorescence of RGEPO2 and GCaMP6f in the neuronal cytoplasm in response to 30 mM KCl stimulation. Imaging was conducted at 20°C-25°C using an inverted wide-field Nikon Eclipse Ti2 microscope, capturing dynamic changes in both potassium and calcium signals over time.

**Supplementary Movie 5:** Time-lapse imaging of Asp-RGEPO1 on the extracellular surface of cultured astrocytes in response to a series of KCl buffer ranging from 0 mM to 20 mM. Imaging was performed at 20°C-25°C using an inverted wide-field Nikon Eclipse Ti2 microscope.

**Supplementary Movie 6:** Time-lapse imaging of RGEPO2 localized in the cytoplasm of cultured astrocytes in response to a series of KCl buffer ranging from 0 mM to 200 Mm without valinomycin or CCCP. Imaging was performed at 20°C-25°C using an inverted wide-field Nikon Eclipse Ti2 microscope.

**Supplementary Movie 7:** Single-plane dual-color imaging of GCaMP6f and RGEPO1 activity from L2/3 neurons of primary somatosensory cortex in awake mouse during KA-induced seizures. Images were acquired using Olympus FVMPE-RS 2P microscope. Imaging conditions: 512 × 512 pixels, 0.994 μm/pixel, acquired at 3Hz. Scale bar, 100μm.

**Supplementary Movie 8**: Single-plane dual-color imaging of GCaMP6f and RGEPO2 activity from L2/3 neurons of primary somatosensory cortex in awake mouse during KA-induced seizures. Images were acquired using Olympus FVMPE-RS 2P microscope. Imaging conditions: 512 × 512 pixels, 0.994 μm/pixel, acquired at 3Hz. Scale bar, 100μm.

## References

1. Palmer, B. F. Regulation Of Potassium Homeostasis. Clin. J. Am. Soc. Nephrol. (2014) doi:10.2215/CJN.08580813.

2. Grafe, P. & Ballanyi, K. Cellular mechanisms of potassium homeostasis in the mammalian nervous system. Can. J. Physiol. Pharmacol. (1987) doi:10.1139/y87-164.

3. Bean, B. P. The action potential in mammalian central neurons. Nature Reviews Neuroscience at 10.1038/nrn2148 (2007).

4. Durand, D. M., Park, E. H. & Jensen, A. L. Potassium diffusive coupling in neural networks. Philosophical Transactions of the Royal Society B: Biological Sciences at 10.1098/rstb.2010.0050 (2010).

5. Verkhratsky, A. & Nedergaard, M. Physiology of astroglia. Physiol. Rev. (2018) doi:10.1152/physrev.00042.2016.

6. Chrysafides, S. M. & Sharma, S. Physiology, Resting Potential. StatPearls (2019).

7. Veech, R. L., Kashiwaya, Y. & King, M. T. The resting membrane potential of cells are measures of electrical work, not of ionic currents. Integr. Physiol. Behav. Sci. (1995) doi:10.1007/BF02691602.

8. Frant, M. S. & Ross, J. W. Potassium ion specific electrode with high selectivity for potassium over sodium. Science (80-.). (1970) doi:10.1126/science.167.3920.987.

9. van de Velde, L., d’Angremont, E. & Olthuis, W. Solid contact potassium selective electrodes for biomedical applications – a review. Talanta at 10.1016/j.talanta.2016.06.050 (2016).

10. Meuwis, K., Boens, N., De Schryver, F. C., Gallay, J. & Vincent, M. Photophysics of the fluorescent K+ indicator PBFI. Biophys. J. (1995) doi:10.1016/S0006-3495(95)80428-5.

11. Golchini, K. et al. Synthesis and characterization of a new fluorescent probe for measuring potassium. Am. J. Physiol. - Ren. Fluid Electrolyte Physiol. (1990) doi:10.1152/ajprenal.1990.258.2.f438.

12. Zhou, X. et al. A new highly selective fluorescent K+ sensor. J. Am. Chem. Soc. (2011) doi:10.1021/ja207345s.

13. Wu, S. Y. et al. A sensitive and specific genetically encoded potassium ion biosensor for in vivo applications across the tree of life. PLoS Biol. (2022) doi:10.1371/journal.pbio.3001772.

14. Torres Cabán, C. C., et al. Tuning the Sensitivity of Genetically Encoded Fluorescent Potassium Indicators through Structure-Guided and Genome Mining Strategies. ACS Sensors 7, 1336–1346 (2022).

15. Shen, Y. et al. Genetically encoded fluorescent indicators for imaging intracellular potassium ion concentration. *Commun*. Biol. (2019) doi:10.1038/s42003-018-0269-2.

16. Bischof, H. et al. Novel genetically encoded fluorescent probes enable real-Time detection of potassium in vitro and in vivo. Nat. Commun. (2017) doi:10.1038/s41467-017-01615-z.

17. Ashraf, K. U. et al. The Potassium Binding Protein Kbp Is a Cytoplasmic Potassium Sensor. Structure (2016) doi:10.1016/j.str.2016.03.017.

18. Mehta, S. et al. Single-fluorophore biosensors for sensitive and multiplexed detection of signalling activities. Nat. Cell Biol. (2018) doi:10.1038/s41556-018-0200-6.

19. Subach, O. M. et al. Frcamp, a red fluorescent genetically encoded calcium indicator based on calmodulin from schizosaccharomyces pombe fungus. Int. J. Mol. Sci. (2021) doi:10.3390/ijms22010111.

20. Wu, J. et al. Genetically Encoded Glutamate Indicators with Altered Color and Topology. ACS Chem. Biol. (2018) doi:10.1021/acschembio.7b01085.

21. Baird, G. S., Zacharias, D. A. & Tsien, R. Y. Circular permutation and receptor insertion within green fluorescent proteins. Proc. Natl. Acad. Sci. U. S. A. (1999) doi:10.1073/pnas.96.20.11241.

22. Nasu, Y. et al. A genetically encoded fluorescent biosensor for extracellular l-lactate. Nat. Commun. (2021) doi:10.1038/s41467-021-27332-2.

23. Gräwe, A. & Stein, V. Linker Engineering in the Context of Synthetic Protein Switches and Sensors. Trends Biotechnol. 39, 731–744 (2021).

24. Nasu, Y., Shen, Y., Kramer, L. & Campbell, R. E. Structure- and mechanism-guided design of single fluorescent protein-based biosensors. Nature Chemical Biology at 10.1038/s41589-020-00718-x (2021).

25. Zacchia, M., Abategiovanni, M. L., Stratigis, S. & Capasso, G. Potassium: From Physiology to Clinical Implications. Kidney Dis. (2016) doi:10.1159/000446268.

26. Beerli, R. R. et al. Isolation of human monoclonal antibodies by mammalian cell display. Proc. Natl. Acad. Sci. U. S. A. (2008) doi:10.1073/pnas.0805942105.

27. Romani, A. M. P. Magnesium homeostasis in mammalian cells. Front. Biosci. (2007) doi:10.2741/2066.

28. Sterns, R. H. Disorders of Plasma Sodium — Causes, Consequences, and Correction. N. Engl. J. Med. (2015) doi:10.1056/nejmra1404489.

29. Palmer, B. F. & Clegg, D. J. Physiology and pathophysiology of potassium homeostasis. Adv. Physiol. Educ. (2016) doi:10.1152/advan.00121.2016.

30. Yamagata, N. The Concentration of Common Cesium and Rubidium in Human Body∗. J. Radiat. Res. (1962) doi:10.1269/jrr.3.9.

31. Protasov, E. S. et al. Erythrocytes as bioreactors to decrease excess ammonium concentration in blood. Sci. Rep. (2019) doi:10.1038/s41598-018-37828-5.

32. Xu, N., Francis, M., Cioffi, D. L. & Stevens, T. Studies on the resolution of subcellular free calcium concentrations: A technological advance. Focus on ‘Detection of differentially regulated subsarcolemmal calcium signals activated by vasoactive agonists in rat pulmonary artery smooth muscle cells’. American Journal of Physiology - Cell Physiology at 10.1152/ajpcell.00046.2014 (2014).

33. Chang, K. L. et al. Zinc at pharmacologic concentrations affects cytokine expression and induces apoptosis of human peripheral blood mononuclear cells. Nutrition (2006) doi:10.1016/j.nut.2005.11.009.

34. Amdisen, A. Clinical and serum-level monitoring in lithium therapy and lithium intoxication. J. Anal. Toxicol. (1978) doi:10.1093/jat/2.5.193.

35. Barondeau, D. P., Kassmann, C. J., Tainer, J. A. & Getzoff, E. D. Structural chemistry of a green fluorescent protein Zn biosensor. J. Am. Chem. Soc. 124, 3522–3524 (2002).

36. Yellen, G. The voltage-gated potassium channels and their relatives. Nature at 10.1038/nature00978 (2002).

37. Boyle, Y., Johns, T. G. & Fletcher, E. V. Potassium Ion Channels in Malignant Central Nervous System Cancers. Cancers at 10.3390/cancers14194767 (2022).

38. Kato, H., Gotoh, H., Kajikawa, M. & Suto, K. Depolarization triggers intracellular magnesium surge in cultured dorsal root ganglion neurons. Brain Res. (1998) doi:10.1016/S0006-8993(97)01232-8.

39. Michod, D. et al. Calcium-Dependent Dephosphorylation of the Histone Chaperone DAXX Regulates H3.3 Loading and Transcription upon Neuronal Activation. Neuron (2012) doi:10.1016/j.neuron.2012.02.021.

40. Choi, D. W. NMDA receptor-mediated K+ efflux and neuronal apoptosis. Science (80-.). (1999) doi:10.1126/science.284.5412.336.

41. Hösli, L., Hösli, E., Landolt, H. & Zehntner, C. Efflux of potassium from neurones excited by glutamate and aspartate causes a depolarization of cultured glial cells. Neurosci. Lett. (1981) doi:10.1016/0304-3940(81)90062-8.

42. Chen, T. W. et al. Ultrasensitive fluorescent proteins for imaging neuronal activity. Nature (2013) doi:10.1038/nature12354.

43. Clarke, L. E. & Barres, B. A. Emerging roles of astrocytes in neural circuit development. Nature Reviews Neuroscience at 10.1038/nrn3484 (2013).

44. Rimmele, T. S. & Chatton, J. Y. A novel optical intracellular imaging approach for potassium dynamics in astrocytes. PLoS One (2014) doi:10.1371/journal.pone.0109243.

45. Köhling, R. & Wolfart, J. Potassium channels in epilepsy. Cold Spring Harbor Perspectives in Medicine at 10.1101/cshperspect.a022871 (2016).

46. de Curtis, M., Uva, L., Gnatkovsky, V. & Librizzi, L. Potassium dynamics and seizures: Why is potassium ictogenic? Epilepsy Research at 10.1016/j.eplepsyres.2018.04.005 (2018).

47. Krishnan, G. P. & Bazhenov, M. Ionic dynamics mediate spontaneous termination of seizures and postictal depression state. J. Neurosci. (2011) doi:10.1523/JNEUROSCI.6200-10.2011.

48. Uva, L. et al. A novel focal seizure pattern generated in superficial layers of the olfactory cortex. J. Neurosci. (2017) doi:10.1523/JNEUROSCI.2239-16.2016.

49. Librizzi, L. et al. Interneuronal network activity at the onset of seizure-like events in entorhinal cortex slices. J. Neurosci. (2017) doi:10.1523/JNEUROSCI.3906-16.2017.

50. Gnatkovsky, V., Librizzi, L., Trombin, F. & De Curtis, M. Fast activity at seizure onset is mediated by inhibitory circuits in the entorhinal cortex in vitro. Ann. Neurol. (2008) doi:10.1002/ana.21519.

51. Dong, A. et al. A fluorescent sensor for spatiotemporally resolved imaging of endocannabinoid dynamics in vivo. Nat. Biotechnol. (2022) doi:10.1038/s41587-021-01074-4.

52. Farrell, J. S. et al. In vivo assessment of mechanisms underlying the neurovascular basis of postictal amnesia. Sci. Rep. (2020) doi:10.1038/s41598-020-71935-6.

53. Stern, M. A., Dingledine, R., Gross, R. E. & Berglund, K. Epilepsy insights revealed by intravital functional optical imaging. Front. Neurol. (2024) doi:10.3389/fneur.2024.1465232.

54. Dana, H. et al. Sensitive red protein calcium indicators for imaging neural activity. Elife (2016) doi:10.7554/eLife.12727.

55. Shen, Y. et al. A genetically encoded Ca2+ indicator based on circularly permutated sea anemone red fluorescent protein eqFP578. BMC Biol. (2018) doi:10.1186/s12915-018-0480-0.

56. Hara, Y. et al. High-affinity tuning of single fluorescent protein-type indicators by flexible linker length optimization in topology mutant. *Commun*. Biol. 7, 705 (2024).

57. Klein, S. G. et al. Toward Best Practices for Controlling Mammalian Cell Culture Environments. Frontiers in Cell and Developmental Biology at 10.3389/fcell.2022.788808 (2022).

58. McNeill, J., Rudyk, C., Hildebrand, M. E. & Salmaso, N. Ion Channels and Electrophysiological Properties of Astrocytes: Implications for Emergent Stimulation Technologies. Frontiers in Cellular Neuroscience at 10.3389/fncel.2021.644126 (2021).

59. Enger, R. et al. Dynamics of ionic shifts in cortical spreading depression. Cereb. Cortex (2015) doi:10.1093/cercor/bhv054.

60. Dreier, J. P., Lemale, C. L., Kola, V., Friedman, A. & Schoknecht, K. Spreading depolarization is not an epiphenomenon but the principal mechanism of the cytotoxic edema in various gray matter structures of the brain during stroke. Neuropharmacology at 10.1016/j.neuropharm.2017.09.027 (2018).

61. Oliveira-Ferreira, A. I., et al. Spreading depolarization, a pathophysiological mechanism of stroke and migraine aura. Future Neurology at 10.2217/fnl.11.69 (2012).

62. Raimondo, J. V., Burman, R. J., Katz, A. A. & Akerman, C. J. Ion dynamics during seizures. Frontiers in Cellular Neuroscience at 10.3389/fncel.2015.00419 (2015).

63. Papadaki, S. et al. Dual-expression system for blue fluorescent protein optimization. Sci. Rep. (2022) doi:10.1038/s41598-022-13214-0.

64. Zhang, H. et al. Quantitative assessment of near-infrared fluorescent proteins. Nat. Methods (2023) doi:10.1038/s41592-023-01975-z.

65. Jumper, J. et al. Highly accurate protein structure prediction with AlphaFold. Nature (2021) doi:10.1038/s41586-021-03819-2.

66. Shu, X., Shaner, N. C., Yarbrough, C. A., Tsien, R. Y. & Remington, S. J. Novel chromophores and buried charges control color in mFruits. Biochemistry (2006) doi:10.1021/bi060773l.

67. Chen, V. B. et al. MolProbity: All-atom structure validation for macromolecular crystallography. Acta Crystallogr. Sect. D Biol. Crystallogr. (2010) doi:10.1107/S0907444909042073.

68. Jurrus, E. et al. Improvements to the APBS biomolecular solvation software suite. Protein Sci. (2018) doi:10.1002/pro.3280.

69. Jorgensen, W. L., Chandrasekhar, J., Madura, J. D., Impey, R. W. & Klein, M. L. Comparison of simple potential functions for simulating liquid water. J. Chem. Phys. (1983) doi:10.1063/1.445869.

70. D.A. Case, K. Belfon, I.Y. Ben-Shalom, S.R. Brozell, D.S. Cerutti, T.E. Cheatham, III, V.W.D. Cruzeiro, T.A. Darden, R.E. Duke, G. Giambasu, M.K. Gilson, H. Gohlke, A.W. Goetz, R Harris, S. Izadi, S.A. Iz- mailov, K. Kasavajhala, A. Kovalenko, R. Krasny, D. M. Y. and P. A. K. (2020). Amber 2020. University of California, San Francisco.

71. Maier, J. A. et al. ff14SB: Improving the Accuracy of Protein Side Chain and Backbone Parameters from ff99SB. J. Chem. Theory Comput. (2015) doi:10.1021/acs.jctc.5b00255.

72. Hix, M. A. & Walker, A. R. AutoParams: An Automated Web-Based Tool To Generate Force Field Parameters for Molecular Dynamics Simulations. J. Chem. Inf. Model. (2023) doi:10.1021/acs.jcim.3c01049.

73. Loncharich, R. J., Brooks, B. R. & Pastor, R. W. Langevin dynamics of peptides: The frictional dependence of isomerization rates of N-acetylalanyl-N′-methylamide. Biopolymers (1992) doi:10.1002/bip.360320508.

74. Robert, X. & Gouet, P. Deciphering key features in protein structures with the new ENDscript server. Nucleic Acids Res. (2014) doi:10.1093/nar/gku316.

75. Roe, D. R. & Cheatham, T. E. PTRAJ and CPPTRAJ: Software for processing and analysis of molecular dynamics trajectory data. J. Chem. Theory Comput. (2013) doi:10.1021/ct400341p.

76. Pettersen, E. F. et al. UCSF ChimeraX: Structure visualization for researchers, educators, and developers. Protein Sci. (2021) doi:10.1002/pro.3943.

77. Humphrey, W., Dalke, A. & Schulten, K. VMD: Visual molecular dynamics. J. Mol. Graph. (1996) doi:10.1016/0263-7855(96)00018-5.

78. Pnevmatikakis, E. A. & Giovannucci, A. NoRMCorre: An online algorithm for piecewise rigid motion correction of calcium imaging data. J. Neurosci. Methods (2017) doi:10.1016/j.jneumeth.2017.07.031.

79. Stringer, C., Wang, T., Michaelos, M. & Pachitariu, M. Cellpose: a generalist algorithm for cellular segmentation. Nat. Methods (2021) doi:10.1038/s41592-020-01018-x.

## Supplement References

1. Shen, Y., et al Genetically encoded fluorescent indicators for imaging intracellular potassium ion concentration. Commun. Biol. (2019).

81. Bischof, H., et al. Novel genetically encoded fluorescent probes enable real-Time detection of potassium in vitro and in vivo. Nat.Commun. (2017)

3. Shen, Y., et al A sensitive and specific genetically encoded potassium ion biosensor for in vivo applications across the tree of life. Plos Biology (2022)

83. Cristina C, et al. Tuning the Sensitivity of Genetically Encoded Fluorescent Potassium Indicators through Structure-Guided and Genome Mining Strategies. ACS sensors (2022)

